# PeroxiHUB: a modular cell-free biosensing platform using H_2_O_2_ as signal integrator

**DOI:** 10.1101/2022.03.16.484621

**Authors:** Paul Soudier, Thomas Duigou, Peter L. Voyvodic, Ana Zúñiga, Kenza Bazi-Kabbaj, Manish Kushwaha, Jerome Bonnet, Jean-Loup Faulon

**Affiliations:** Université Paris-Saclay, INRAE, AgroParisTech, Micalis Institute, 78352 Jouy-en-Josas, France; Université de Montpellier, INSERM, CNRS, Centre de Biologie Structurale, 34090 Montpellier, France

**Keywords:** synthetic biology, cell-free systems, biosensor, hydrogen peroxide, H_2_O_2_, enzymatic transducer, Computer Aided Design

## Abstract

Cell-free systems have great potential for delivering robust, cheap, and field-deployable biosensors. Many cell-free biosensors rely on transcription factors responding to small molecules, but their discovery and implementation still remain challenging. Here we report the engineering of PeroxiHUB, an optimized H_2_O_2_-centered sensing platform supporting cell-free detection of different metabolites. H_2_O_2_ is a central metabolite and a by-product of numerous enzymatic reactions. PeroxiHUB uses enzymatic transducers to convert metabolites of interest into H_2_O_2_, enabling rapid reprogramming of sensor specificity using alternative transducers. We first screen several transcription factors and optimize OxyR for the transcriptional response to H_2_O_2_ in cell-free, highlighting the need for pre-incubation steps to obtain suitable signal-to-noise ratios. We then demonstrate modular detection of metabolites of clinical interest – lactate, sarcosine, and choline – using different transducers mined via a custom retro-synthesis workflow publicly available on the SynBioCAD Galaxy portal. We find that expressing the transducer during the pre-incubation step is crucial for optimal sensor operation. Finally, we show that different reporters can be connected to PeroxiHUB, providing high adaptability for various applications. Given the wide range of enzymatic reactions producing H_2_O_2_, the PeroxiHUB platform will support cell-free detection of a large number of metabolites in a modular and scalable fashion.

## INTRODUCTION

Detection and quantification of metabolites and other small molecules is an important area of research with applications in many fields, such as disease diagnostics and prognostics^1^, pollutant or pathogen detection, agricultural or industrial process monitoring, and fundamental research methodologies. Most of these challenges are currently being addressed by a combination of analytical physics and chemistry techniques, including chromatography, spectrometry, titrimetry, and optical and electrochemical methods^2, 3^.

Biological systems and related devices have the potential to replace some of these time-, cost- and equipment-expensive methods. Living cells enclose machinery capable of interacting with particular small molecules, including substrate specific enzymes or metabolite-binding transcriptional factors (TFs). These systems have been successfully repurposed into highly responsive whole-cell biosensors able to detect a wide diversity of molecules^4^. Cell-free transcription translation (TX-TL) systems are abiotic, cell-derived biological mixtures that are able to emulate some biological reactions and features *in vitro*. TX-TL systems have followed a continuous development since the 1960s, from their use in the deciphering of the genetic code^5^ to their repurposing into platforms integrating synthetic biology devices over the last two decades^6^. TX-TL systems can integrate various types of biosensors from riboswitches to TF-mediated systems^7, 8^.

Cell-free biosensors present a variety of advantages over whole-cell systems that support their broad use as point-of-use sensing devices. They are abiotic, and thus not subjected to the same GMO regulations as living sensors, and can be freeze-dried for long-term room temperature storage^9^. Moreover, the absence of an intact living and reproducing cellular compartment enables the sensing of molecules that are deleterious for cell growth or those that do not cross the cell membrane.

Researchers have engineered cell-free biosensors to detect nucleic acids and small molecules^10^. While the modular nature of Watson-Crick base pairing supports the engineering of tailor-made sensors for different nucleic acid sequences, metabolite detection follows mostly an *ad hoc* approach, in which specific transcription factors known to respond to small-molecules are co-opted. Compared to other methods, sensing systems derived from transcription factors have multiple advantages, including good specificity and response versatility, thanks to the variety of possible gene expression outputs. They are also suited to carry out complex computational behavior and calculate an output as a function of the concentrations of multiple input molecules^11^.

However, the development of new TF-based sensors in cell-free systems has been hindered by a variety of factors. First, the number of documented transcriptional effectors binding desired chemicals is limited; second, the complexity of their regulatory mechanisms sometimes prevents their implementation in a simplified cell-free system; and third, most cell-free systems use *E. coli* extracts, which can limit the effectiveness of transcription-promoting mechanisms derived from other species.

Recently, we devised an alternative approach to extend the number of potential cell-free metabolite biosensors by using enzymatic transducers to transform a non-detectable molecule into a ligand for a known, characterized transcription factor^12, 13^. Yet, even using metabolic transducers, the small number of TF-based sensors functional in cell-free, and the complexity of the molecules resulting from enzymatic reactions, limits the number of compounds detectable via this approach. In order to circumvent this issue, we aimed to design a signal integration system in which many metabolic transducers modifying several different molecules produce a common metabolite that can be detected by a single transcription factor. To develop this sensing “hub”, we chose hydrogen peroxide (H_2_O_2_), a central metabolism molecule and a ubiquitous product of several enzymatic reactions, as a common signaling molecule.

Here we develop an optimized, TF-based, cell-free H_2_O_2_ sensing platform that, coupled with computer-predicted enzymatic transducers, is able to detect a wide range of small molecules through the activation of various reporter genes. We identify TFs and promoter combinations with the best response to H_2_O_2_ in a cell-free environment and optimize the reaction conditions for high-signal/low-noise hydrogen peroxide detection. We then build a computational tool implemented as a Galaxy workflow to help identify enzymatic transducer candidates for custom metabolite sensing. We determine and optimize key factors enabling these enzymes to mediate the sensor response in the cell-free reaction. As a proof-of-concept, we built sarcosine, lactate and choline biosensors. Importantly, connecting the metabolic transducers to our cell-free H_2_O_2_ sensing hub requires little additional optimization. In addition, we show that our platform can accommodate various output modalities, expanding the range of possible applications. The highly modular sensing platform presented here will enable fast development of new biosensors for custom metabolite detection with reduced screening efforts.

## RESULTS

H_2_O_2_ is a suitable candidate to act as a metabolic hub for several reasons. First, multiple H_2_O_2_-responsive transcription factors and target promoters have been identified, providing an appropriate space for exploring and optimizing an H_2_O_2_ transcriptional response system. Second, unlike many other metabolites(e.g. amino acids) or cofactors (e.g.NAD, Coenzyme A), H_2_O_2_ is not part of the cell-free buffer, limiting interference with the sensor. Finally, H_2_O_2_ is a central metabolite and a byproduct of many enzymatic reactions, which enables connection with many metabolites. This is demonstrated by the Rhea database^14^ that documents more than 350 different enzymatic reactions producing H_2_O_2_.

### A Galaxy webtool for custom H_2_O_2_ transducer mining

To map the space of molecules potentially detectable through enzymatic reactions producing H_2_O_2,_ we developed a Galaxy workflow, named BioSensor (**Figure 1A, Supplementary Figure S1**), combining the RetroPath2.0 software with rp2biosensor, a new bioinformatic tool. The BioSensor workflow can be used on the SynBioCAD Galaxy platform^15^ (accessible at https://galaxy-synbiocad.org/workflows/list_published). It is also available on the Galaxy ToolShed, which enables its installation on any other Galaxy server.

**Figure 1:**
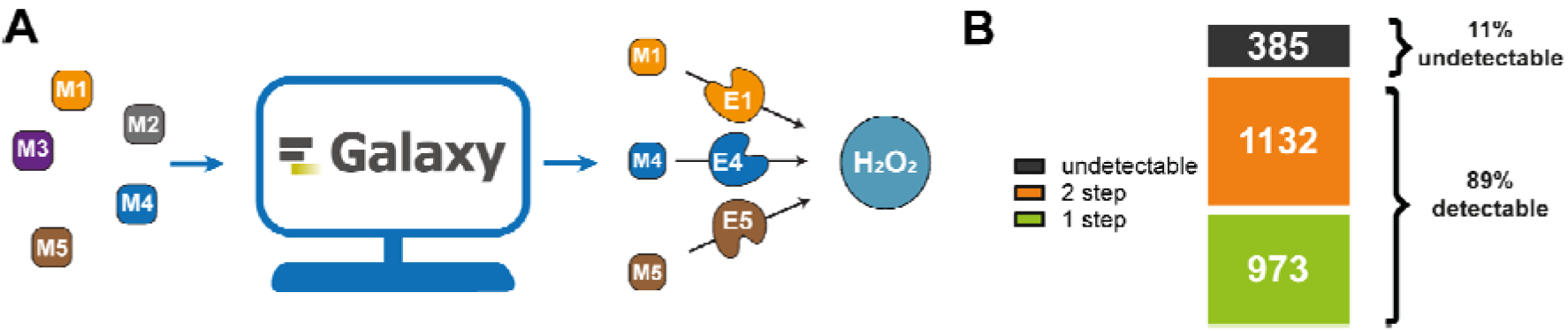
Development and evaluation of a Galaxy workflow for predicting H_2_O_2_-producing reactions. **(A)** Principle of the workflow for custom H_2_O_2_-producing transducer mining. M: Metabolites of interest; E: enzymatic transducers. **(B)** Results of the Biosensor Galaxy workflow for the prediction of metabolic pathways connecting disease-related molecules to H_2_O_2_.

The BioSensor workflow enables the prediction of metabolic reactions connecting any query metabolite to H_2_O_2_. When given the InChI identifier of the molecule to detect, the workflow returns an interactive web page showing, if they are known, the potential pathways connecting the target to the chosen detectable molecule and provide additional information, such as MetaNetX^16^ reaction IDs or EC numbers, to facilitate the identification of potentials enzymatic transducers (**Supplementary Figure S2**).

A new feature of this newly developed tool over the previously released biosensor prediction tool, SensiPath^17^, is the integration of potential promiscuous activity of the predicted enzymatic transducers.

Formalization of enzyme promiscuity using reaction rules has previously been described with RetroPath^18^ and RetroRules^19^. Briefly, promiscuity is modeled by the atomic environment around reaction centers. Increasing the scope of this description -- the diameter surrounding reaction centers -- leads to a more restrictive description about what the substrate should look like, hence increasing the modeled enzyme specificity.

By reducing the diameter constraint of the reaction in the query panel, it is possible to identify new potential sensing routes that take advantage of potential promiscuous activities of enzymes to expand the solution space.

As a pilot study, we used the Galaxy Biosensor workflow to identify molecules of interest that could be converted to H_2_O_2_ with 1 or 2 enzymatic steps, focusing on disease-associated metabolites according to the HMDB database^20^ (**Figure 1B**). We found that of a total of 2490 molecules, 2105 were potentially detectable through enzymatic reactions producing H_2_O_2_, 973 with one step and 1132 with two enzymatic steps. Another encouraging metric is that out of the 1965 sensing enabling metabolic pathways, 1788 rely on reactions with a diameter at least equal to 12, the highest one tested, which suggest good specificity and applicability of them as metabolic transducers. Together these numbers confirm the high connectivity of H_2_O_2_ in metabolic reactions networks. Convinced by this large potential sensing space, we then started to develop the PeroxiHUB platform, a cell-free H_2_O_2_ biosensor able to detect this large variety of compounds on demand through the production of various output signals (**Figure 2A**).

**Figure 2:**
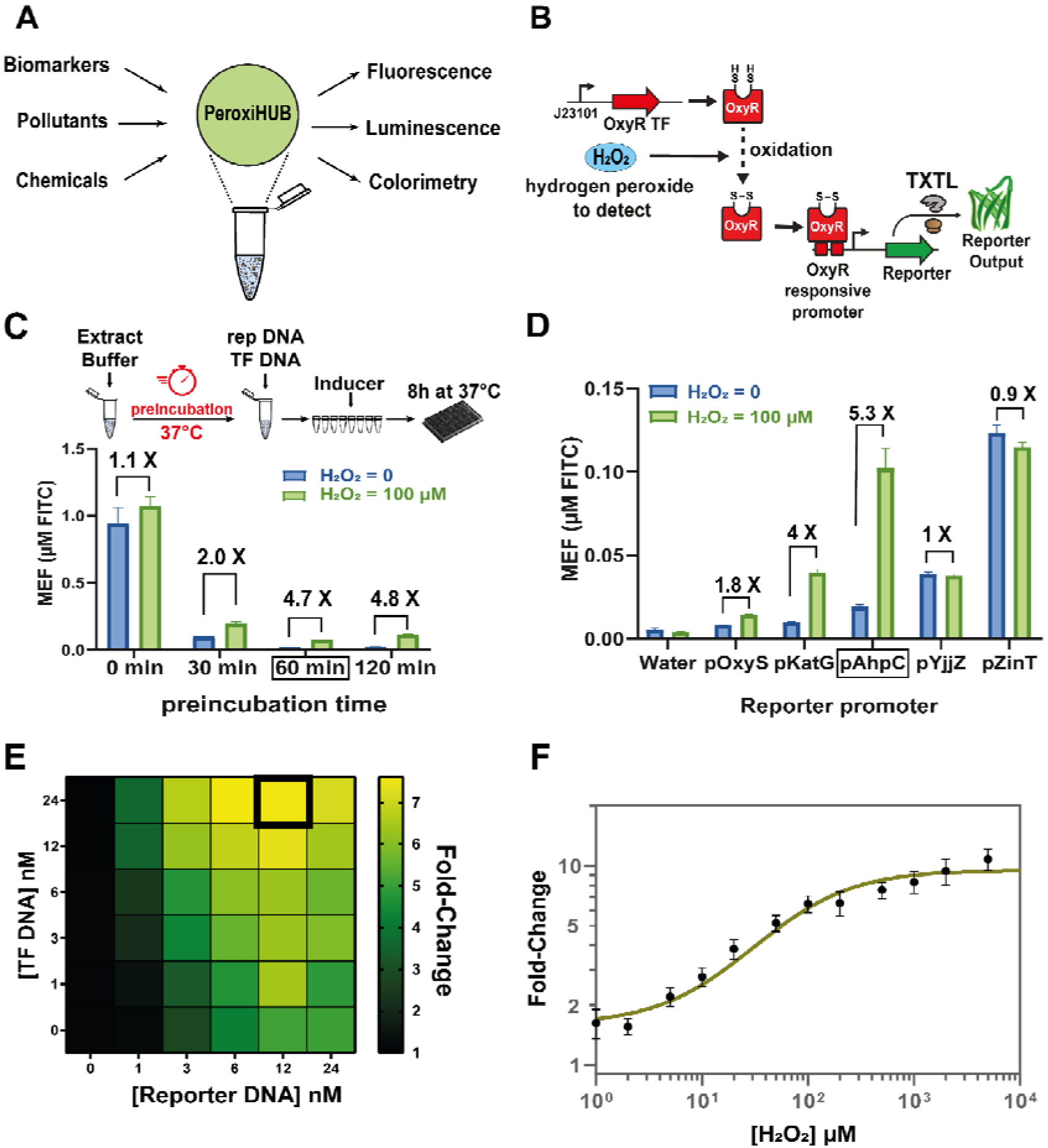
Implementation and optimization of an H_2_O_2_ sensor in cell-free systems. **(A)** Concept of the PeroxiHUB sensor: an optimized cell-free H_2_O_2_ biosensor is used as a hub to detect various molecules with custom output. **(B)** Implementation of an H_2_O_2_ sensor in cell-free. The sensor is composed of two plasmids: one constitutively producing the transcription factor OxyR that reacts with H_2_O_2_ before activating an inducible promoter producing the reporter on the second plasmid. **(C)** Preincubation of cell-free extract and buffer alone at 37 °C for various times before addition of the other components (DNA and inducer) strongly modulates the sensor response, increasing the fluorescence fold-change represented by the number above bars. pAhpC-GFP and J23101-OxyR plasmids were used at a concentration of 10 nM. **(D)** Screening of multiple OxyR interacting promoters reveals various responses to H_2_O_2_ induction after 8h of incubation at 37°. **(E)** DNA concentration gradient test for the transcription factor and the reporter expressing plasmids enables fine tuning of these protein expression levels and optimizes the response of the sensor by increasing the fold change of fluorescence between 0 and 100μM of H_2_O_2_ to more than 7 after 8h of incubation at 37°. **(F)** Final H_2_O_2_ sensing dose response curve evaluated after 8h of incubation at 37° using all the previously optimized conditions(1 hour preincubation, [J23101-OxyR DNA] at 24nM and [pAhpC-sfGFP DNA] at 12 nM,. The fit of the curve was obtained from the mean of three different cell-free reactions. Boxes in (**C**)-(**E**) indicated the selected optimized condition. Error bars represent the standard deviation calculated from 3 individual cell-free reactions. MEF (Mean Equivalent Fluorescence) quantifies the fluorescence measured by the plate reader as equal to the one generated by a certain amount of FITC.

### Implementation and optimization of an H_2_O_2_ biosensor in cell-free

To implement an H_2_O_2_ transcriptional biosensor operating in cell-free, we adapted the design from Rubens et al.^21^ previously used in bacterial cells. This biosensor relies on the OxyR transcription factor, a master regulator involved in the response to oxidative stress in multiple bacterial species, including *E. coli*. OxyR switches from an inactive, reduced state to an active, oxidized one upon reaction with H_2_O_2_, becoming a transcriptional activator^22^. To implement OxyR in a cell-free environment, we used a two-plasmid design (**Figure 2B**): one expressing the OxyR gene under the control of a strong constitutive promoter J23101 (available as a biobrick in the iGEM repository), the other expressing sfGFP under the control of an OxyR-responsive promoter.

Initial implementation of the biosensor according to the reported *in vivo* design^21^ showed no significant response to H_2_O_2_ (**Supplementary Figure S3**), mostly because of a high transcriptional noise, even in the absence of the inducer. We thus focused on identifying and optimizing the parameters influencing the sensor response for a cell-free reaction.

We hypothesized that the high background was due to endogenous H_2_O_2_ production in the cell-free reaction coming from the catabolism of buffer components by enzymes present in the extract. With H_2_O_2_ being an unstable molecule, a pre-incubation step could help reduce this noise. By pre-incubating the cell-free extract with only the buffer at 37°C before adding the plasmid DNA and the inducer, we observed a strong drop in total fluorescence but an increase in the response fold change at 100 μM H_2_O_2_, from 1.1- to more than 4-fold (**Figure 2C**). The optimal signal-to-noise ratio was observed after a 1 hour preincubation using the OxyR-expressing plasmid combined with the pAhpC reporter plasmid. All subsequent experiments include this preincubation step.

We then screened multiple promoters described in the literature: pOxyS, pKatG and pAhpC coming from the *in vivo* sensor design^21^ and the promoters pZinT and pYjjZ activated by the OxyR *in vivo.* All the promoters produced GFP in the cell-free mix but only the first three demonstrated a noticeable response to 100μM of H_2_O_2_ (**Figure 2D**).

pAhpC was identified as the optimal candidate with which to build a biosensor due to its large fold-change and high expression level upon activation. We then evaluated the best combination of expression levels for reporter and TF by measuring the fold-change of the biosensor in the presence of concentration gradients of the two plasmids (**Figure 2E**). An optimum was found at [pAhpC-GFP DNA] = 12nM and [J23101-OxyR DNA] = 24 nM. In these conditions, the fold-change in response to 100 μM H_2_O_2_ was increased to more than 7-fold. Using these calibrated parameters, the biosensor was capable of detecting H_2_O_2_ over several orders of magnitude, from micromolar to millimolar concentrations, with a fold-change up to 10.8, and an EC50 of 75 μM, a relatively low leakage and a high swing (**Figure 2F, Table 1**).

**Table 1:** Performance of the H_2_O_2_ sensor. Metrics of the hydrogen peroxide sensor were calculated on the data presented on figure 2B. The max fold change is the fluorescence in non induced and induced state, the leakage is the fluorescence in the non induced state, the swing is the difference of fluorescence between the non induced and induced state. EC50 is the half-maximal effective concentration. MEF (Mean Equivalent Fluorescence) quantifies the fluorescence measured by the plate reader as equal to the one generated by a certain amount of FITC.

| Metric | $\text{H}_2\text{O}_2$ sensor |
| --- | --- |
| Max fold change | $11 \pm 2$ |
| Leakage MEF (FITC) | $0.03 \pm 0$ |
| Max swing MEF (FITC) | $0.30 \pm 0.05$ |
| EC50 ($\mu\text{M}$) | 75 |

### Optimizing enzymatic conversion of custom metabolites into hydrogen peroxide

In order to demonstrate the PeroxiHUB concept and its potential for future applications, we used the BioSensor workflow to identify candidate enzymatic transducers for three central metabolites: sarcosine, choline, and lactate. These molecules are all central metabolites, identified as disease biomarkers but also of potential interest in other fields. As an example, they all have been described being of potential interest for diagnostic or prognostic of prostate cancer^23–25^

Pathways and enzymes producing H_2_O_2_ directly from sarcosine and choline were identified using the Galaxy workflow. For lactate processing, the workflow suggested several different enzymes, but for the biosensor implementation we opted for one previously validated from the literature^26^. Consistent with our modularity objective, the transducers were implemented by supplementing the optimal, two-plasmid H_2_O_2_ transcription biosensor with an additional plasmid expressing the specific enzyme under the control of a constitutive promoter (**Figure 3A**).

**Figure 3:**
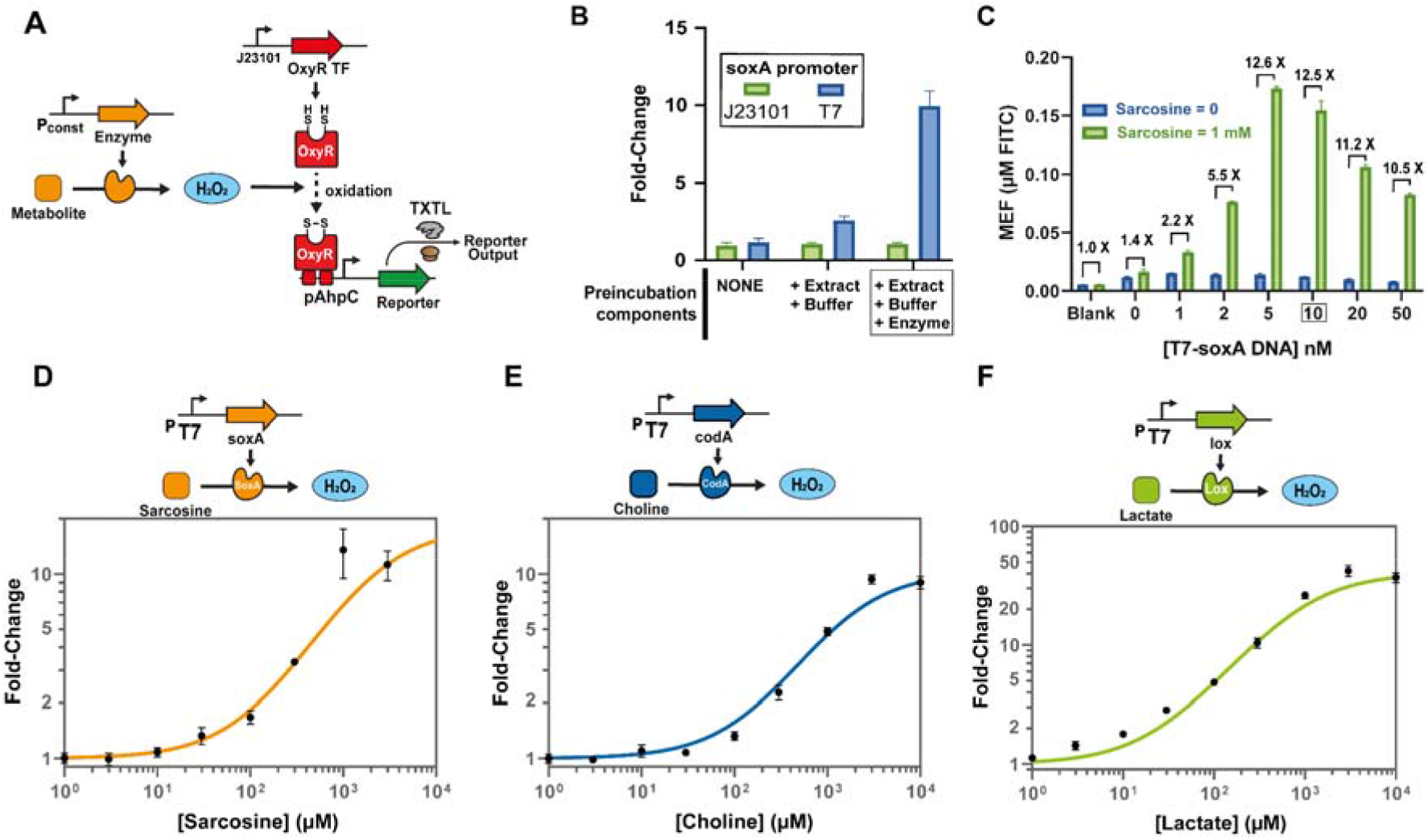
Computer predicted enzymatic transducers with optimized expression conditions enable custom metabolite sensing. **(A)** Implementation of enzyme-mediated biosensors: a plasmid expressing the enzyme predicted to generate H_2_O_2_ from the target is added to the optimized H_2_O_2_ sensor. **(B)** optimization of enzyme expression conditions for sarcosine transducer: expression of the enzyme under a T7 promoter added before the preincubation step maximizes sarcosine sensing. Fold-change is calculated as the ratio of fluorescence produced between 1mM of sarcosine and no sarcosine. **(C)** Fine tuning of enzyme expression using a DNA concentration gradient is necessary to identify the best condition for the sarcosine transducer. Data for the other transducers can be found in Supplementary Figure S4. **(D)** Dose response curve of optimized SoxA-mediated sarcosine biosensor. **(E)** Dose response curve of optimized CodA-mediated choline biosensor. **(F)** Dose response curve of optimized lox-mediated lactate biosensor. Error bars represent the standard deviation calculated from 3 replicates.

Initial assays in which the transducer SoxA was cloned into the same backbone as the plasmid used to express OxyR with the strong constitutive bacterial promoter J23101 were unsuccessful, with no detectable response to sarcosine even at high plasmid concentrations in the cell-free mix (**Figure 3B**). We reasoned that expression of the enzyme in this configuration was too low for sensor function, potentially because of insufficient promoter strength and resource limitations. Using sarcosine oxidase (SoxA) as a model, we thus tested if switching from J23101 to the strong T7 promoter, which relies on a different polymerase pool than the other expressed components, could help solve our issue and limit resource competition. Finally, we also increased the total pool of available transducers at the beginning of the detection reaction by expressing enzymes during the 1h preincubation.

For SoxA, these optimizations drastically increased the fold-change of the biosensor to ~10 at 1 mM of sarcosine (**Figure 3B**). Fine-tuning the level of expression of the enzyme by varying the concentration of the DNA template also had a major impact on the response of the sensor. The optimal transducer plasmid concentration was variable between different transducers (10 nM for *soxA*, 24 nM for *codA*, and 1 nM for *lox*) (**Figure 3C, Supplementary Figure S4**) highlighting the need for DNA concentration gradient screening for each new transducer developed. Final sensors for sarcosine, lactate and choline were characterized over a gradient of inducer concentrations, showing good response over several orders of magnitude (**Figure 3D–F, Table 2, Supplementary Figure S5**).

**Table 2:** Performance of the sarcosine, choline and lactate sensor. Metrics of the Sarcosine, Choline and Lactate sensor were calculated on the data presented on figure 3D 3E & 3F. The max fold change is the ratio of fluorescence between the uninduced and the induced states, the leakage is the fluorescence measured in the uninduced state, the swing is the difference of fluorescence between the uninduced and induced states. EC50 is the half-maximal effective concentration. MEF (Mean Equivalent Fluorescence) quantifies the fluorescence measured by the plate reader as equal to the one generated by a certain amount of FITC.

| Metric | Sarcosine sensor | Choline sensor | Lactate sensor |
| --- | --- | --- | --- |
| Max fold change | 13.5 ± 4.0 | 9.3 ± 0.5 | 42 ± 4.5 |
| Leakage MEF (FITC) | 0.00 ± 0 | 0.01 ± 0 | 0.00 ± 0 |
| Max swing MEF (FITC) | 0.16 ± 0.05 | 0.18 ± 0.01 | 0.27 ± 0.04 |
| EC50 (μM) | 1933 | 1535 | 934 |

To simplify the future use of such biosensors, we evaluated the possibility to flash freeze preincubated batches of cell-free mix in liquid nitrogen, enabling their storage at −80°C and immediate later use without any additional preincubation steps. The experimental results showed few differences in response between frozen and unfrozen preincubated extracts, opening the way for broad use of these sensors without an increase in detection time from the incubation step (**Supplementary Figure S6**).

### Expanding the range of detectable reporter outputs

One advantage of cell-free biosensors producing a transcriptional response is that their output can be easily changed according to the final application needs (such as read-out modality, timing, or signal processing). To expand the potential of the PeroxiHUB sensing platform, we connected the H_2_O_2_ biosensor to different reporter genes. Colorimetry and luminescence were chosen as they represent classical signals used in sensing devices, with the potential for naked-eye detection and faster measurements (**Figure 4A**). The colorimetric output was implemented using lacZ as a reporter gene in an extract made from a *ΔlacZ* BL21 strain. The cell-free mix was then supplemented with CPRG (Chlorophenol red-β-D-galactopyranoside), which is converted from yellow to the purple-colored CPR (Chlorophenol red) by LacZ (**Figure 4B**). The resulting output can be either identified visually or quantified by monitoring absorbance of the reaction at 574 nm. Using an internal ladder for quantification, previous work has demonstrated the feasibility of robust cell-free biosensors in low-resource conditions using this output^27^. We explored various CPRG and reporter DNA concentrations to obtain the best differentiation of colorimetric output inside the H_2_O_2_ sensing range. Time progression of the absorbance at 574 nm followed a sigmoidal function with a maximum principally dependent upon the initial CPRG concentration and a kinetic component governed by the reporter DNA concentration. The ideal conditions were determined to be [CPRG] = 100 μM and [pAhpC-LacZ DNA] = 6 nM (**Supplemental Figures S7 & S8**). These conditions brought the direct sensing of H_2_O_2_ with the colorimetric output to a lower limit of detection than when using GFP, with detectable concentrations at the micromolar level (**Figure 4C**). Sarcosine sensing was also demonstrated to be possible over a wide range of concentrations, although without the increase in sensitivity observed for the H_2_O_2_ sensor (**Figure 4D**).

**Figure 4:**
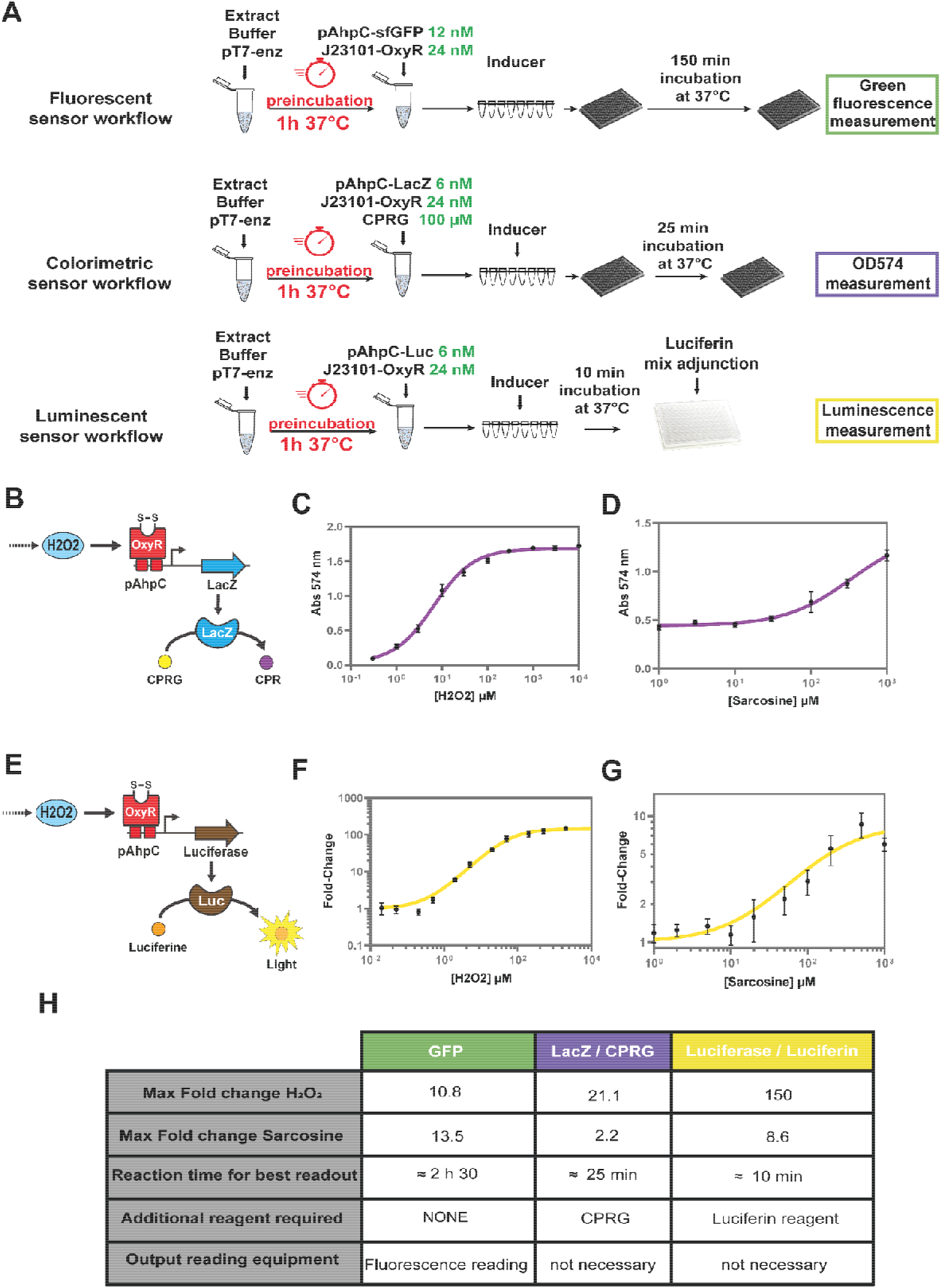
Luminescent and colorimetric modular reporters for PeroxiHUB platform. **(A)** Experimental workflows followed for each reporter. **(B)** Design and implementation of the colorimetric reporter. **(C)** Colorimetric hydrogen peroxide sensor dose response curve. **(D)** Colorimetric sarcosine sensor dose response curve. **(E)** Design and implementation of the luminescent reporter. **(F)** Luminescent hydrogen peroxide sensor dose response curve. **(G)** Luminescent sarcosine sensor dose response curve. **(H)** Comparison of potentialities and limits of each reporting system developed. All error bars represent the standard deviation calculated from 3 replicates.

Similar to the development of the colorimetric reporter, the luciferase-based output was adapted from a previous design implemented in cell-free^12^. The luciferase enzyme produced in response to H_2_O_2_ reacts with the luciferin reagent to generate a detectable light output **(Figure 4E)**. After the incubation step, the 20 μL reaction mix is supplemented with the luciferin-containing reagent and luminescence intensity was measured (see **Figure 4A**). Unlike the fluorescent and colorimetric outputs, this system showed no background in the absence of the reporter plasmid, confirming that the cell-free mix has no endogenous luminescence **(Supplementary Figure S9).** We tested different durations for the detection step to maximize the sensor’s response to H_2_O_2_ and identified a ten minute incubation time as optimal before measuring luminescence. Indeed, ten minutes were sufficient for the sensor to generate a significant signal, while longer incubation resulted in increased noise, thus decreasing the signal-to-noise ratio **(Supplementary Figure S10)**. Using these parameters, the H_2_O_2_ response of the sensor was much higher than what was observed with the other reporters, with response going up to 150-fold-change between non-induced and induced states **(Figure 4F)**. When testing this reporter for enzyme-mediated detections with the example of sarcosine, the output observed was generally in the same range of fold-change as what was observed with the GFP reporter (**Figure 4G**). However, even if the highest observed fold-change was slightly lower with the luciferase output (8.6-fold) than the one observed with GFP (13.5-fold), the luminescent output was found to be better at low concentration (1.7-fold vs 3.0-fold between GFP and Luciferase at a concentration of 100 μM) which supports its use for low-concentration inducer detection. Due to its much faster reaction time, the luciferase output presents a convenient redout for the PeroxiHUB platform **(Figure 4H)**.

## DISCUSSION

Here we developed a cell-free, modular sensing hub that generates a transcriptional output using H_2_O_2_ as a common signaling currency. The PeroxiHUB platform enables the detection of different molecules via the use of metabolic transducers producing H_2_O_2_ as a by-product of their enzymatic reaction. Because of the large number of enzymatic reactions producing H_2_O_2_, PeroxiHUB is highly-modular and allows detection of new molecules of interest by simply switching enzymatic transducers. We used PeroxiHUB to detect three different metabolites and found that only a one-step enzymatic transducer plasmid concentration tuning was necessary, while all the other reaction parameters could be kept constant. The core of the method includes the H_2_O_2_ sensor with invariant optimal conditions, the T7-containing backbone for the transducer enzyme cloning and the preincubation conditions.

Among the parameters shown to impact the response of the H_2_O_2_ sensor, the most important one is the preincubation step, which is necessary for both the sensing of the H_2_O_2_ and for the proper expression of enzymatic transducers. These two effects seem independent as a preincubation step with extract and buffer only is sufficient to improve the behavior of the H_2_O_2_ sensor to work, but using that preincubation step to also produce the enzyme has a major impact on the performance of transducer-mediated sensors. The improvement in H_2_O_2_ sensor performance is in part due to the overall reduction of the reporter expression after preincubation, reducing background noise. One hypothesis to explain the high background of non-preincubated reactions is the presence of endogenous H_2_O_2_ produced by enzymatic reactions originating from the extract. Pre-incubating the cell-free reaction then allows endogenous catalase activities to degrade that initial pool of H_2_O_2_, reducing the noise and increasing the fold-change of the biosensor.

The other beneficial impact of preincubation, an increase in enzymatic transducer activity, is most likely expression-related. Preincubation likely allows the transducer to be synthesized at high level and available before the start of the detection reaction. The higher limit of detection observed for enzyme mediated sensors compared to the H_2_O_2_ sensor suggests that the enzymatic activity is the key bottleneck limiting the efficiency of transducer-based sensors. The importance of enzyme concentration is also evidenced by the effects of changes in the promoter driving enzyme expression and in DNA template concentration.

Enzyme expression likely has two opposite effects: on the one hand, their expression by the cell-free system leads to a reduction in the overall expression levels for other proteins by resource competition. On the other hand, they increase the H_2_O_2_ pool and the biosensor’s response. These effects, vary between the tested transducer, and affect both sensor signal level of signal and background noise in non-induced condition **(Supplementary Figure S5).** Consequently, the various transducers have differential apparent efficiencies, the Lox-based sensor producing a 3- to 5-times higher max fold-change compared to the SoxA- and CodA-based ones **(Table 2)**.

We have not investigated at this stage the source of variations in transducers behaviors. Many parameters could be involved, including enzyme kinetic, expression levels, folding, stability, or the presence of potential inhibitors within the extract. Future studies might improve a particular transducer by testing different homologs^28^, or using directed evolution to reach higher enzymatic activity.

With efficient enzymes and optimized expression conditions, it should also be possible to extend the range of detectable molecules by using multiple, successive enzymatic conversions leading to H_2_O_2_. These multi-step enzymatic conversions can be identified using the BioSensor Galaxy workflow. Indeed, as demonstrated by the metrics coming from the HMDB panel (**Figure 1B**), the already high number of potentially detectables molecules through 1-step enzymatic conversions can be greatly expanded by plugging in an additional reaction step.

We also demonstrated that the platform is amenable to the use of different reporter systems, expanding the range of application contexts. Together, the various reporting possibilities expand the range of applications for which PeroxiHub can be used. They all present some advantages and drawbacks that promote or discourage their use for a specific application in different contexts. GFP is the simplest and cheapest reporting system as it doesn’t require any additional chemicals and shows a relatively good response. The LacZ/CPRG-mediated colorimetric output is faster and does not require equipment for qualitative measurement, which makes it a good reporting system for portable and low-resource detection problems. Finally, the luciferase output is the fastest and the most sensitive to low inducer concentrations **(Figure 4H),** and could still be associated with portable readout systems such as smartphone based platforms^29^. The platform could also be expanded to additional outputs by expanding its connectivity with existing or newly-developed detection and monitoring systems. For example, the recently implemented glucose/glucometer cell-free reporting system^30^ could be modularly adapted to the platform to provide a cheap digital readout of measured concentrations.

Finally, another promising application of the PeroxiHUB platform is its use as a signal integrator for cell-free computational devices, such as analog computing systems. We previously built a 4-input metabolic perceptron classifying samples in a binary manner on the basis of their concentration of four different metabolites. To do so, metabolites underwent enzyme-mediated conversion into a single detectable molecule, in this case benzoate^11^. The key bottleneck in the generalization of such computing devices is the identification of central “hub” molecules detectable in cell-free systems and in which several metabolites of interest could be converted. The PeroxiHub platform appears as an attractive candidate for such a task.

## MATERIALS and METHODS

### BioSensor Galaxy workflow development and node description

The new Biosensor workflow was developed within the Galaxy SynBioCAD portal following the general methodology described in the original paper establishing the platform^15^. It is also released on the Galaxy ToolShed^31^, which enables its installation on any other Galaxy server. Existing nodes present within the SynBioCAD environment were connected to the custom, newly developed rp2biosensor node. Below is the description of the main nodes composing the Biosensor workflow.

**RetroPath2.0** is an open-source tool designed to build a retrosynthesis network linking a compound of interest to one or more other compounds^18^. The compound of interest is provided by its structure, and chemical transformations formalized by reaction rules are applied, which predict newly formed products of the simulated reactions. For two or more steps of exploration, new products of the previous step are used as substrates and reaction rules are applied again. This operation is performed until the number of steps is reached or no new products are found. RetroPath2.0 is available at myExperiment.org (https://www.myexperiment.org/workflows/4987.html), as a conda package on anaconda.org (https://anaconda.org/conda-forge/retropath2_wrapper), as well as a Galaxy node on the Galaxy ToolShed (https://toolshed.g2.bx.psu.edu/view/tduigou/retropath2/9c8ac9980bd6). RetroPath2.0 release r20220104 was used.

**rp2biosensor** is an open-source Python software that extracts from the retrosynthetic network generated by RetroPath2.0 the subnetwork of interest, linking the biosensor to the compound to be detected, and produce an interactive web page showing the transducing reactions. Briefly, rp2biosensor parses the retrosynthesis network outputted by RetroPath2.0, completes the predicted reactions by putting back co-substrates and co-products omitted during the retrosynthesis using the rxn_rebuild (https://github.com/brsynth/rxn_rebuild) Python package, enumerates the shortest path linking the compound of interest, i.e. the biosensor, to the compound to be detected, e.g. lactate, and finally outputs the resulting subnetwork as an interactive web page to let the user browse the results. rp2biosensor source code is available on GitHub (https://github.com/brsynth/rp2biosensor), is released as a conda package on anaconda.org (https://anaconda.org/conda-forge/rp2biosensor), and as a Galaxy node on the Galaxy ToolShed (https://toolshed.g2.bx.psu.edu/view/tduigou/rp2biosensor/b0efd4b2ffba). rp2biosensor version 3.0.0 was used.

### BioSensor Galaxy workflow executions

The typical use case requires the user to input the chemical structure of a compound to be detected and eventually the chemical structure of a TF effector (prefilled with the structure of H_2_O_2_ by default). Structures should be provided using the InChI standard format. The output is an interactive web page that can be opened within the Galaxy environment. Thanks to the Galaxy workflow system, all intermediate and final outputs can be easily downloaded for later usages.

For the prediction of reactions enabling the detection of (*S*)-lactate, choline and sarcosine, their standard InChIs have been used as input for the “Molecule to be detected parameter” (see **Supplementary Table S1**). The BioSensor workflow was launched for one step of exploration, using reaction rules precompiled for both “reverse” and “forward” usage, with diameters ranging from 2 to 12, the default values of the workflow.

For the efficiency assessment of the developed tool, the HMDB database version 5.0 was used to explore detectable compounds from H_2_O_2_. Only compounds fulfilling the following criteria have been kept: compounds should be associated with at least one disease, have a valid InChI structure, contain at least one carbon, and have a molecular weight of at most 1 kDa. RetroPath2.0 was set for a 2-step exploration using both forward and retro reaction rules with diameters ranging from 2 to 12. If both 1- and 2-step pathways exist for a given compound, only shortest paths are reported by rp2biosensor.

### Plasmid construction and purification

Plasmids used in this study were constructed using Gibson assembly method with pBEAST as a vector backbone^12^ and inserts either amplified from the *E. coli* genome (for the OxyR gene, the AhpC, OxyS and KatG promoters) or from existing plasmids (for Luciferase gene amplified from pBen-Luc used in Voyvodic et al.^12^) or synthesized as gene fragments (IDT, for the soxA, codA, lox and genes and the ZinT and YjjZ promoters, Twist Bioscience for LacZ gene). All genes synthesized were codon-optimized for *E. coli.*

### Bacterial strains and growth conditions

Clonings and plasmid amplifications were made using the classical *E. coli* cloning strain DH5α or the commercially available NEB® Turbo strain. Liquid cultures were made at 37°C using LB media with 100 μg mL^−1^ ampicillin for the maintenance of the pBEAST derived plasmids. For solid cultures 1.5 % agar w/v was added.

For cell-free extract preparation, the strain BL21 (DE3) Gold dLacZ (a gift from Jeff Hasty (Addgene plasmid # 99247)) was grown in 2-YTP media supplemented with 50 μg mL^−1^ tetracycline.

### Cell-free reaction mix preparation

Cell-free TX-TL extract was prepared following a protocol adapted from Sun et al.^32^ previously used in other work from our lab^12^. Cultures were grown to an OD600 of 2.0 and centrifuged at 5000xg for 12 min at 4°C. The pellets were washed several times by resuspension/centrifugation cycles before being weighed and stored overnight in 50 mL tubes at −80°C. The pellets were then resuspended in 1 mL S30A buffer (14 mM Mg-glutamate, 60 mM K-glutamate, 50 mM Tris pH 7.7) per gram of pellet, thawed, and lysed by a single pass through an Avestin EmulsiFlex-C3 homogeniser at 15000-20000 psi. The resultant lysate was centrifuged at 12000xg for 30 min at 4°C, then incubated 1 h at 37°C before being centrifuged again with the same settings. Finally, the supernatant was dialysed overnight inside a 12-14 kDa molecular weight cut-off (MWCO) dialysis tubing inS30B buffer (14 mM Mg-glutamate, 60 mM K-glutamate, ~5 mM Tris pH 8.2) before being centrifuged one final time at 12000xg for 30 min, aliquoted in 1.5 ml tubes, flash frozen in liquid nitrogen and stored at −80°C until use.

One aliquot was used for buffer calibration in order to determine the best concentrations of Mg-glutamate, K-Glutamate and DTT to maximize protein production. Consecutive cell-free experiments were run expressing constitutive GFP in the presence of gradients of these three components, following the methodology described in Sun et al.^32^. After the ideal conditions were determined, a batch of buffer was prepared in a single Falcon tube to ensure homogeneity, before being aliquoted in 2 mL tubes, flash frozen in liquid nitrogen and stored at −80 °C until use.

### Cell-free reaction preparation

For cell-free reactions, buffer and extract aliquots were thawed on ice. Each reaction was prepared in individual PCR tubes containing 22 μL total mix: 7.33 μL extract, 9.17 μL buffer and 5.5 μL of other components (DNA, inducer, water and any additional chemicals). Once all the components added to the PCR tubes the mixes were briefly vortexed and spun down, 20 μL were pipetted into a 384-round-well non-binding plate for reporter gene expression measurement. In some cases, a master mix containing all the components present at the same level in all the conditions tested (e.g. extract, buffer, DNA) was prepared prior to pipetting into the individual PCR tubes. All the experiments were run in triplicate.

### Reporter signal measurements

To measure GFP fluorescence, 8 hour kinetics were performed at 37°C with either a Cytation 3 a Synergy HTX plate reader (Biotek Instruments) using excitation/emission settings of 485 nm and 528 nm, respectively.

Collected data were normalized by FITC Mean Equivalent Fluorescence (MEF) through conversion factors that were established for each plate reader using fluorescein standards with the same plates and machine settings as the ones used in the experiments, as per Jung et al.^33^ & Batista et al.^28^. For CPRG reporter measurement, OD574 was measured. Data were normalized by subtracting a blank sample containing everything but reporter DNA.

For the luciferase reporter, after the 37°C incubation step, 20 μL of the final cell-free reactions mix were added to a white 96-well plate. 50 μL of Luciferin reagent mix (Promega, Luciferase Assay Reagent) were then added to each well, mixed by pipetting up and down, and the plate was immediately inserted inside the plate reader to capture luminescence.

## Author Contributions

PS, MK, JB, & JLF conceived the study. TD & KBK developed and published the bioinformatic tool. PS, PV & AZ designed the experiments. PS performed the experiments. PS & AZ analyzed the results. JLF & JB acquired the funding. PS and JB wrote the manuscript with contributions from all authors. All authors read and approved the final manuscript. The authors declare no competing interest.

## Supporting Information

The Supplementary Information file (PDF) contains Supplementary Figures S1-S9 and Supplementary Tables S1-S5.

**Supplementary Figures:** (S1) Detail of the SynbioCAD based Galaxy Biosensor workflow, (S2) Example of result graph output from the Biosensor workflow for Sarcosine, (S3) Unoptimized H2O2 sensor response in cell-free system, (S4) Fine tuning of enzyme expression using DNA gradient, (S5) Final sensors fluorescent dose response, (S6) Liquid Nitrogen Flash-Freezing of preincubated mix,(S7) CPRG concentration optimization for colorimetric H2O2 biosensor, (S8) [pAhpC-LacZ DNA] concentration optimization for colorimetric H2O2 biosensor, (S9) Luminescent sensor early evaluation, (S10) Luminescent sensor incubation time optimization.

**Supplementary Tables:** (S1) Chemicals identifiers used in the study, (S2) Characteristics of enzymes used in the study, (S3) Plasmids used in this study, (S4) DNA sequences for constructs used in this study

## Acknowledgements

PS, PV, JLF and JB acknowledge funding support by the ANR SynBioDiag grant (ANR-18-CE33-0015). JLF acknowledges funding support by the ANR grants iCFree (BiopNSE grant number 13001533) and Aladin (EQUIPEX+ grant number 15000882). JB acknowledges support from the ERC starting “COMPUCELL” (grant number 657579), INSERM and the Bettencourt Schuller foundation. The Centre de Biologie Structurale acknowledges support from the French Infrastructure for Integrated Structural Biology (FRISBI) (ANR-10-INSB-05-01). MK acknowledges funding support from INRAe’s MICA department, Université Paris-Saclay, Ile de France (IdF) region’s DIM-RFSI, and ANR DREAMY (ANR-21-CE48-003).

## REFERENCES

(1) Spratlin, J. L.; Serkova, N. J.; Eckhardt, S. G. Clinical Applications of Metabolomics in Oncology: A Review. Clin. Cancer Res. 2009, 15 (2), 431–440. https://doi.org/10.1158/1078-0432.CCR-08-1059.

(2) Siddiqui, M. R.; AlOthman, Z. A.; Rahman, N. Analytical Techniques in Pharmaceutical Analysis: A Review. Arab. J. Chem. 2017, 10, S1409–S1421. https://doi.org/10.1016/j.arabjc.2013.04.016.

(3) Beale, D. J.; Pinu, F. R.; Kouremenos, K. A.; Poojary, M. M.; Narayana, V. K.; Boughton, B. A.; Kanojia, K.; Dayalan, S.; Jones, O. A. H.; Dias, D. A. Review of Recent Developments in GC–MS Approaches to Metabolomics-Based Research. Metabolomics 2018, 14 (11), 152. https://doi.org/10.1007/s11306-018-1449-2.

(4) Park, M.; Tsai, S.-L.; Chen, W. Microbial Biosensors: Engineered Microorganisms as the Sensing Machinery. Sensors 2013, 13 (5), 5777–5795. https://doi.org/10.3390/s130505777.

(5) Nirenberg, M. W.; Matthaei, J. H. The Dependence of Cell-Free Protein Synthesis in E. Coli upon Naturally Occurring or Synthetic Polyribonucleotides. Proc. Natl. Acad. Sci. 1961, 47 (10), 1588–1602. https://doi.org/10.1073/pnas.47.10.1588.

(6) Hodgman, C. E.; Jewett, M. C. Cell-Free Synthetic Biology: Thinking Outside the Cell. Metab. Eng. 2012, 14 (3), 261–269. https://doi.org/10.1016/j.ymben.2011.09.002.

(7) Tabuchi, T.; Yokobayashi, Y. Cell-Free Riboswitches. RSC Chem. Biol. 2021, 2 (5), 1430–1440. https://doi.org/10.1039/D1CB00138H.

(8) Koch, M.; Pandi, A.; Borkowski, O.; Batista, A. C.; Faulon, J.-L. Custom-Made Transcriptional Biosensors for Metabolic Engineering. Curr. Opin. Biotechnol. 2019, 59, 78–84. https://doi.org/10.1016/j.copbio.2019.02.016.

(9) Pardee, K.; Green, A. A.; Takahashi, M. K.; Braff, D.; Lambert, G.; Lee, J. W.; Ferrante, T.; Ma, D.; Donghia, N.; Fan, M.; Daringer, N. M.; Bosch, I.; Dudley, D. M.; O’Connor, D. H.; Gehrke, L.; Collins, J. J. Rapid, Low-Cost Detection of Zika Virus Using Programmable Biomolecular Components. Cell 2016, 165 (5), 1255–1266. https://doi.org/10.1016/j.cell.2016.04.059.

(10) Zhang, L.; Guo, W.; Lu, Y. Advances in Cell-Free Biosensors: Principle, Mechanism, and Applications. Biotechnol. J. 2020, 15 (9), 2000187. https://doi.org/10.1002/biot.202000187.

(11) Pandi, A.; Koch, M.; Voyvodic, P. L.; Soudier, P.; Bonnet, J.; Kushwaha, M.; Faulon, J.-L. Metabolic Perceptrons for Neural Computing in Biological Systems. Nat. Commun. 2019, 10 (1), 3880. https://doi.org/10.1038/s41467-019-11889-0.

(12) Voyvodic, P. L.; Pandi, A.; Koch, M.; Conejero, I.; Valjent, E.; Courtet, P.; Renard, E.; Faulon, J.-L.; Bonnet, J. Plug-and-Play Metabolic Transducers Expand the Chemical Detection Space of Cell-Free Biosensors. Nat. Commun. 2019, 10 (1), 1697. https://doi.org/10.1038/s41467-019-09722-9.

(13) Libis, V.; Delépine, B.; Faulon, J.-L. Expanding Biosensing Abilities through Computer-Aided Design of Metabolic Pathways. ACS Synth. Biol. 2016, 5 (10), 1076–1085. https://doi.org/10.1021/acssynbio.5b00225.

(14) Alcántara, R.; Axelsen, K. B.; Morgat, A.; Belda, E.; Coudert, E.; Bridge, A.; Cao, H.; de Matos, P.; Ennis, M.; Turner, S.; Owen, G.; Bougueleret, L.; Xenarios, I.; Steinbeck, C. Rhea—a Manually Curated Resource of Biochemical Reactions. Nucleic Acids Res. 2012, 40 (Database issue), D754–D760. https://doi.org/10.1093/nar/gkr1126.

(15) Lac, M. du; Duigou, T.; Hérisson, J.; Carbonell, P.; Swainston, N.; Zulkower, V.; Shah, F.; Faure, L.; Mahdy, M.; Soudier, P.; Faulon, J.-L. Galaxy-SynBioCAD: Synthetic Biology Design Automation Tools in Galaxy Workflows. bioRxiv June 15, 2020, p 2020.06.14.145730. https://doi.org/10.1101/2020.06.14.145730.

(16) Moretti, S.; Tran, V. D. T.; Mehl, F.; Ibberson, M.; Pagni, M. MetaNetX/MNXref: Unified Namespace for Metabolites and Biochemical Reactions in the Context of Metabolic Models. Nucleic Acids Res. 2021, 49 (D1), D570–D574. https://doi.org/10.1093/nar/gkaa992.

(17) Delépine, B.; Libis, V.; Carbonell, P.; Faulon, J.-L. SensiPath: Computer-Aided Design of Sensing-Enabling Metabolic Pathways. Nucleic Acids Res. 2016, 44 (W1), W226–W231. https://doi.org/10.1093/nar/gkw305.

(18) Delépine, B.; Duigou, T.; Carbonell, P.; Faulon, J.-L. RetroPath2.0: A Retrosynthesis Workflow for Metabolic Engineers. Metab. Eng. 2018, 45, 158–170. https://doi.org/10.1016/j.ymben.2017.12.002.

(19) Duigou, T.; du Lac, M.; Carbonell, P.; Faulon, J.-L. RetroRules: A Database of Reaction Rules for Engineering Biology. Nucleic Acids Res. 2019, 47 (D1), D1229–D1235. https://doi.org/10.1093/nar/gky940.

(20) Wishart, D. S.; Tzur, D.; Knox, C.; Eisner, R.; Guo, A. C.; Young, N.; Cheng, D.; Jewell, K.; Arndt, D.; Sawhney, S.; Fung, C.; Nikolai, L.; Lewis, M.; Coutouly, M.-A.; Forsythe, I.; Tang, P.; Shrivastava, S.; Jeroncic, K.; Stothard, P.; Amegbey, G.; Block, D.; Hau, David. D.; Wagner, J.; Miniaci, J.; Clements, M.; Gebremedhin, M.; Guo, N.; Zhang, Y.; Duggan, G. E.; MacInnis, G. D.; Weljie, A. M.; Dowlatabadi, R.; Bamforth, F.; Clive, D.; Greiner, R.; Li, L.; Marrie, T.; Sykes, B. D.; Vogel, H. J.; Querengesser, L. HMDB: The Human Metabolome Database. Nucleic Acids Res. 2007, 35 (suppl_1), D521–D526. https://doi.org/10.1093/nar/gkl923.

(21) Rubens, J. R.; Selvaggio, G.; Lu, T. K. Synthetic Mixed-Signal Computation in Living Cells. Nat. Commun. 2016, 7 (1), 11658. https://doi.org/10.1038/ncomms11658.

(22) Zheng, M.; Åslund, F.; Storz, G. Activation of the OxyR Transcription Factor by Reversible Disulfide Bond Formation. Science 1998, 279 (5357), 1718–1722. https://doi.org/10.1126/science.279.5357.1718.

(23) Sreekumar, A.; Poisson, L. M.; Rajendiran, T. M.; Khan, A. P.; Cao, Q.; Yu, J.; Laxman, B.; Mehra, R.; Lonigro, R. J.; Li, Y.; Nyati, M. K.; Ahsan, A.; Kalyana-Sundaram, S.; Han, B.; Cao, X.; Byun, J.; Omenn, G. S.; Ghosh, D.; Pennathur, S.; Alexander, D. C.; Berger, A.; Shuster, J. R.; Wei, J. T.; Varambally, S.; Beecher, C.; Chinnaiyan, A. M. Metabolomic Profiles Delineate Potential Role for Sarcosine in Prostate Cancer Progression. Nature 2009, 457 (7231), 910–914. https://doi.org/10.1038/nature07762.

(24) Awwad, H. M.; Geisel, J.; Obeid, R. The Role of Choline in Prostate Cancer. Clin. Biochem. 2012, 45 (18), 1548–1553. https://doi.org/10.1016/j.clinbiochem.2012.08.012.

(25) Tessem, M.-B.; Swanson, M. G.; Keshari, K. R.; Albers, M. J.; Joun, D.; Tabatabai, Z. L.; Simko, J. P.; Shinohara, K.; Nelson, S. J.; Vigneron, D. B.; Gribbestad, I. S.; Kurhanewicz, J. Evaluation of Lactate and Alanine as Metabolic Biomarkers of Prostate Cancer Using 1H HR-MAS Spectroscopy of Biopsy Tissues. Magn. Reson. Med. 2008, 60 (3), 510–516. https://doi.org/10.1002/mrm.21694.

(26) Taurino, I.; Reiss, R.; Richter, M.; Fairhead, M.; Thöny-Meyer, L.; De Micheli, G.; Carrara, S. Comparative Study of Three Lactate Oxidases from Aerococcus Viridans for Biosensing Applications. Electrochimica Acta 2013, 93, 72–79. https://doi.org/10.1016/j.electacta.2013.01.080.

(27) McNerney, M. P.; Zhang, Y.; Steppe, P.; Silverman, A. D.; Jewett, M. C.; Styczynski, M. P. Point-of-Care Biomarker Quantification Enabled by Sample-Specific Calibration. Sci. Adv. 5 (9), eaax4473. https://doi.org/10.1126/sciadv.aax4473.

(28) Batista, A. C.; Levrier, A.; Soudier, P.; Voyvodic, P. L.; Achmedov, T.; Reif-Trauttmansdorff, T.; DeVisch, A.; Cohen-Gonsaud, M.; Faulon, J.-L.; Beisel, C. L.; Bonnet, J.; Kushwaha, M. Differentially Optimized Cell-Free Buffer Enables Robust Expression from Unprotected Linear DNA in Exonuclease-Deficient Extracts. ACS Synth. Biol. 2022, 11 (2), 732–746. https://doi.org/10.1021/acssynbio.1c00448.

(29) Zhang, D.; Liu, Q. Biosensors and Bioelectronics on Smartphone for Portable Biochemical Detection. Biosens. Bioelectron. 2016, 75, 273–284. https://doi.org/10.1016/j.bios.2015.08.037.

(30) Amalfitano, E.; Karlikow, M.; Norouzi, M.; Jaenes, K.; Cicek, S.; Masum, F.; Sadat Mousavi, P.; Guo, Y.; Tang, L.; Sydor, A.; Ma, D.; Pearson, J. D.; Trcka, D.; Pinette, M.; Ambagala, A.; Babiuk, S.; Pickering, B.; Wrana, J.; Bremner, R.; Mazzulli, T.; Sinton, D.; Brumell, J. H.; Green, A. A.; Pardee, K. A Glucose Meter Interface for Point-of-Care Gene Circuit-Based Diagnostics. Nat. Commun. 2021, 12 (1), 724. https://doi.org/10.1038/s41467-020-20639-6.

(31) Blankenberg, D.; Von Kuster, G.; Bouvier, E.; Baker, D.; Afgan, E.; Stoler, N.; Taylor, J.; Nekrutenko, A.; Galaxy Team. Dissemination of Scientific Software with Galaxy ToolShed. Genome Biol. 2014, 15 (2), 403. https://doi.org/10.1186/gb4161.

(32) Sun, Z. Z.; Hayes, C. A.; Shin, J.; Caschera, F.; Murray, R. M.; Noireaux, V. Protocols for Implementing an Escherichia Coli Based TX-TL Cell-Free Expression System for Synthetic Biology. JoVE J. Vis. Exp. 2013, No. 79, e50762. https://doi.org/10.3791/50762.

(33) Jung, J. K.; Alam, K. K.; Verosloff, M. S.; Capdevila, D. A.; Desmau, M.; Clauer, P. R.; Lee, J. W.; Nguyen, P. Q.; Pastén, P. A.; Matiasek, S. J.; Gaillard, J.-F.; Giedroc, D. P.; Collins, J. J.; Lucks, J. B. Cell-Free Biosensors for Rapid Detection of Water Contaminants. Nat. Biotechnol. 2020, 38 (12), 1451–1459. https://doi.org/10.1038/s41587-020-0571-7.

