## Supplementary Information for "PeroxiHUB: a modular cell-free biosensing platform using H_2_O_2_ as signal integrator"

### Table of Contents


| | | Page # |
|---|---|---|
| **Supplementary Figures** | | |
| S1 | Detail of the SynbioCAD based Galaxy Biosensor workflow | S3 |
| S2 | Example of result graph output from the Biosensor workflow for Sarcosine | S4 |
| S3 | Unoptimized H2O2 sensor response in cell-free system | S5 |
| S4 | Fine tuning of enzyme expression using DNA gradient | S6 |
| S5 | Final sensors fluorescent dose response | S7 |
| S6 | Liquid Nitrogen Flash-Freezing of preincubated mix | S8 |
| S7 | CPRG concentration optimization for colorimetric H2O2 biosensor | S9 |
| S8 | [pAhpC-LacZ DNA] concentration optimization for colorimetric H2O2 biosensor | S10 |
| S9 | Luminescent sensor early evaluation | S11 |
| S10 | Luminescent sensor incubation time optimisation | S12 |
| **Supplementary Tables** | | |
| S1 | Chemicals identifiers used in the study | S13 |
| S2 | Characteristics of enzymes used in the study | S13 |
| S3 | Plasmids used in this study | S14 |
| S4 | DNA sequences for constructs used in the study | S15 - S33 |

**Fig S1:**

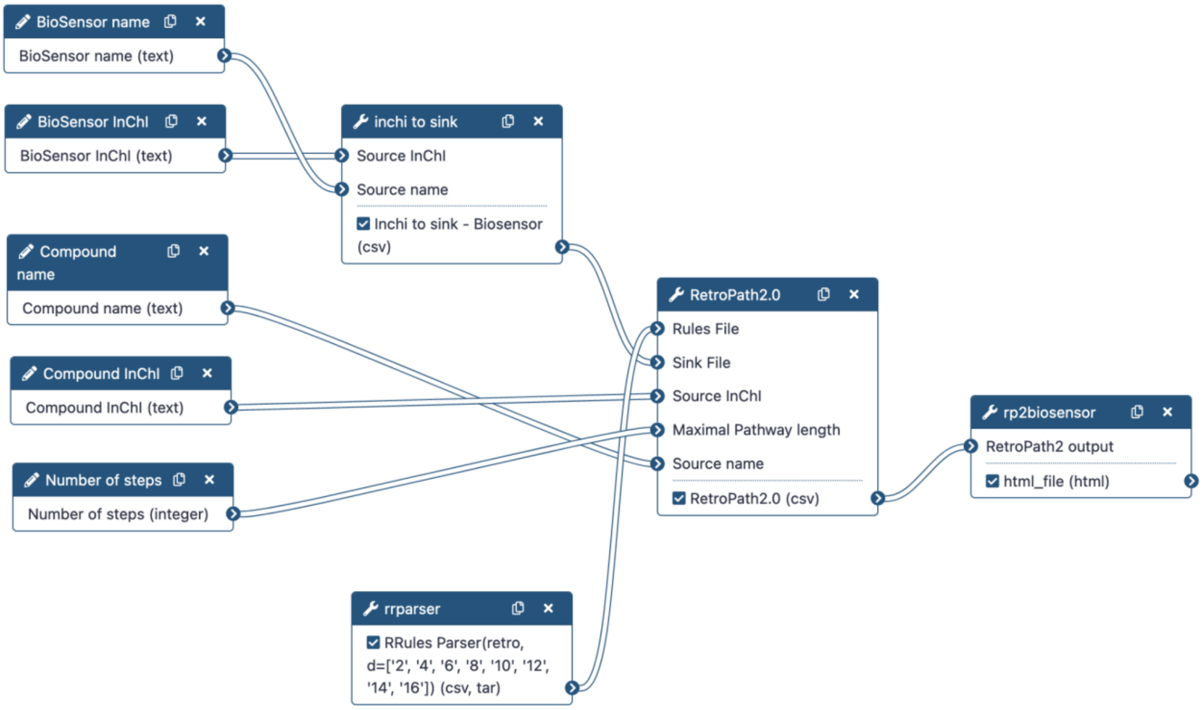


**Supplementary Figure S1: Complete view of the BioSensor Galaxy workflow automatizing biosensor predictions.** Characteristics of the main nodes are described in the method section.

**Fig S2:**

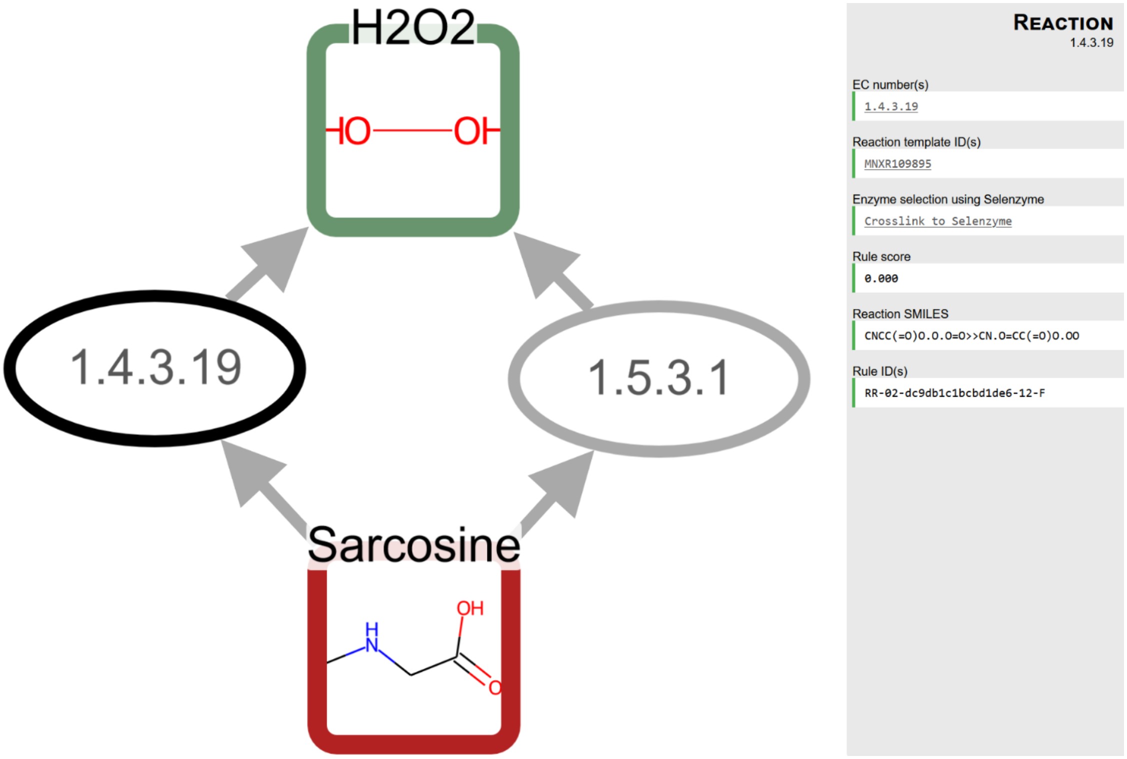


**Supplementary Figure S2: Example of result web page output from the Biosensor workflow for sarcosine.**

**Fig S3:**

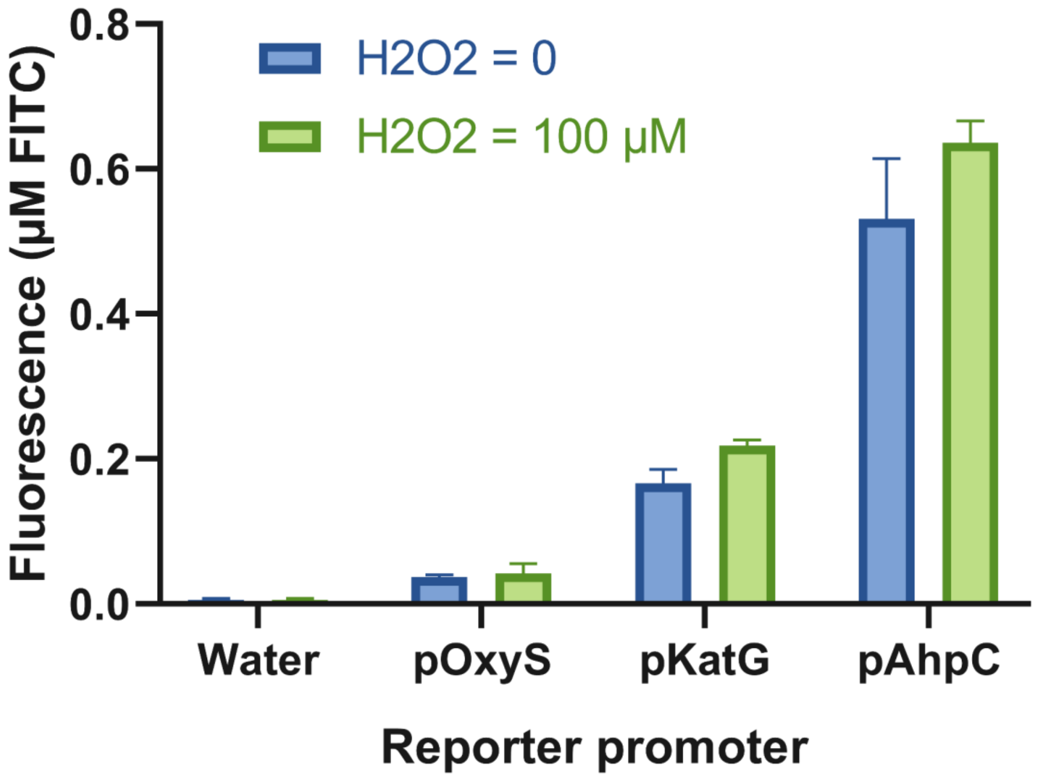


**Supplementary Figure S3: Unoptimized $H_2O_2$ sensor response in cell-free system.** Direct cell-free implementation of biosensor candidates derived from *in vivo* design did not show significant response to $H_2O_2$. 16.5µL of Cell-free mix was supplemented with 1.1µL of TF and Reporter plasmid each, 1.1µL water and 2.2µL of either water or $H_2O_2$. The final concentrations were of 10 nM for each plasmid added and 100µM for the $H_2O_2$ inducer. Fluorescence values are MEF measured at an endpoint of 8h

**Fig S4:**

## A

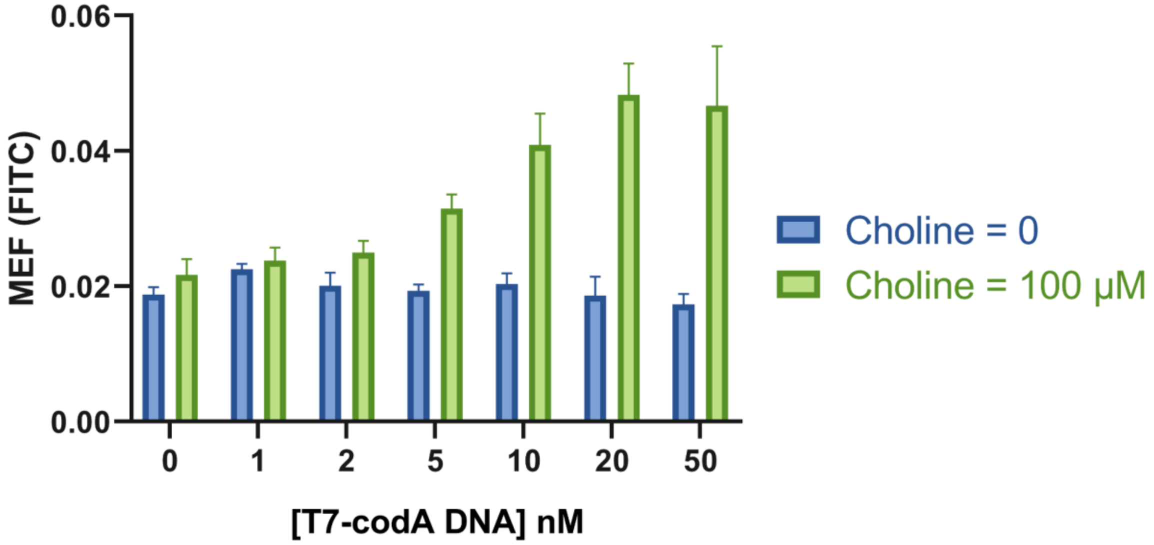


## B

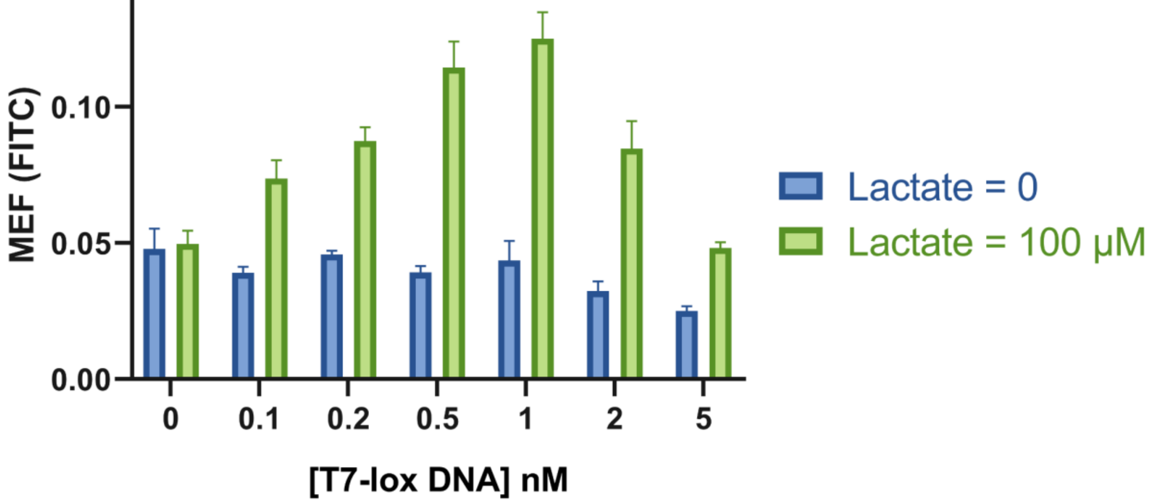


**Supplementary Figure S4: Fine tuning of enzyme expression using DNA gradient (A)** [T7-codA DNA] optimisation for choline sensing **(B)** [T7-lox DNA] optimisation for lactate sensing

Fig S5:

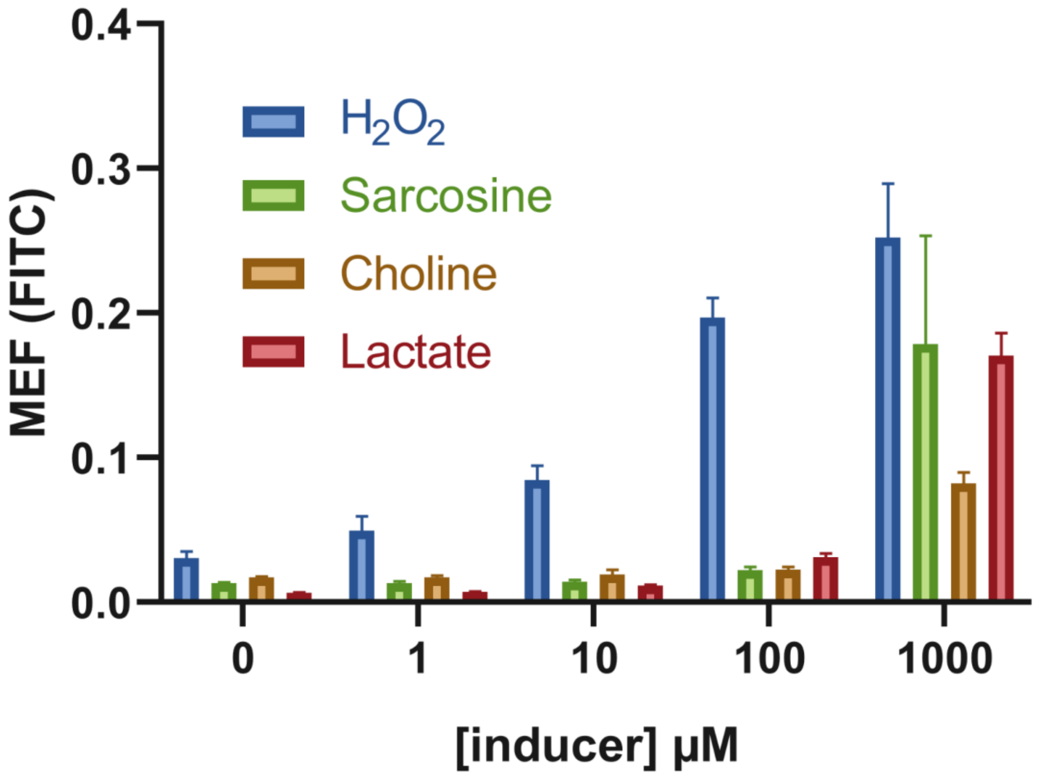


**Supplementary Figure S5: Final sensors fluorescent dose response**

Fig S6:

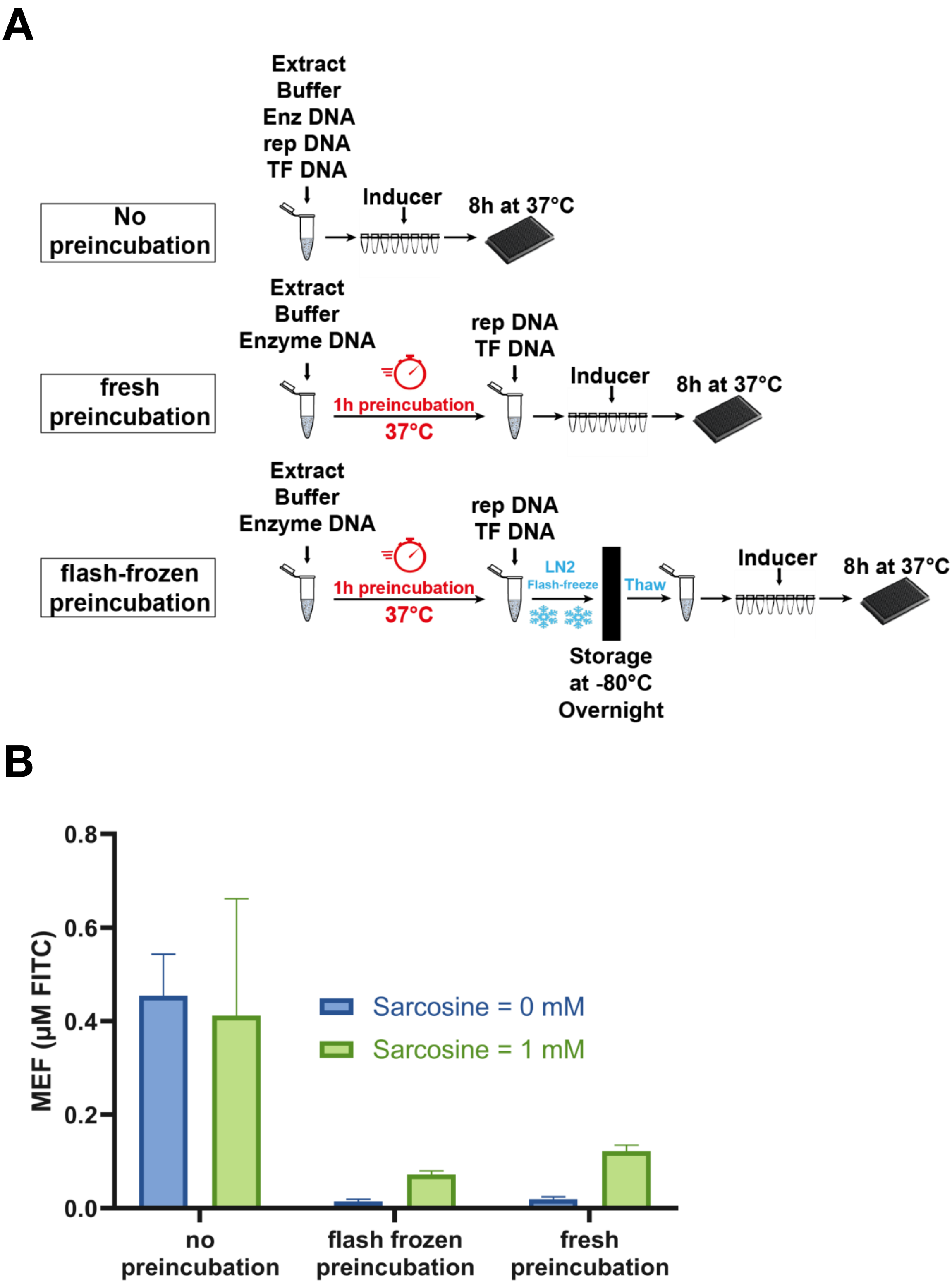


**Supplementary Figure S6: Liquid Nitrogen Flash-Freezing of preincubated mix (A)** Experimental workflows detailing the no preincubation, the fresh preincubation and the flash frozen preincubation pipelines used for biosensors evaluation **(B)** 2h30 endpoint fluorescence taken in presence and absence of inducer for each pipeline tested reveals few differences of responses between the fresh and the flash frozen preincubation workflows.

**Fig S7:**

**A**

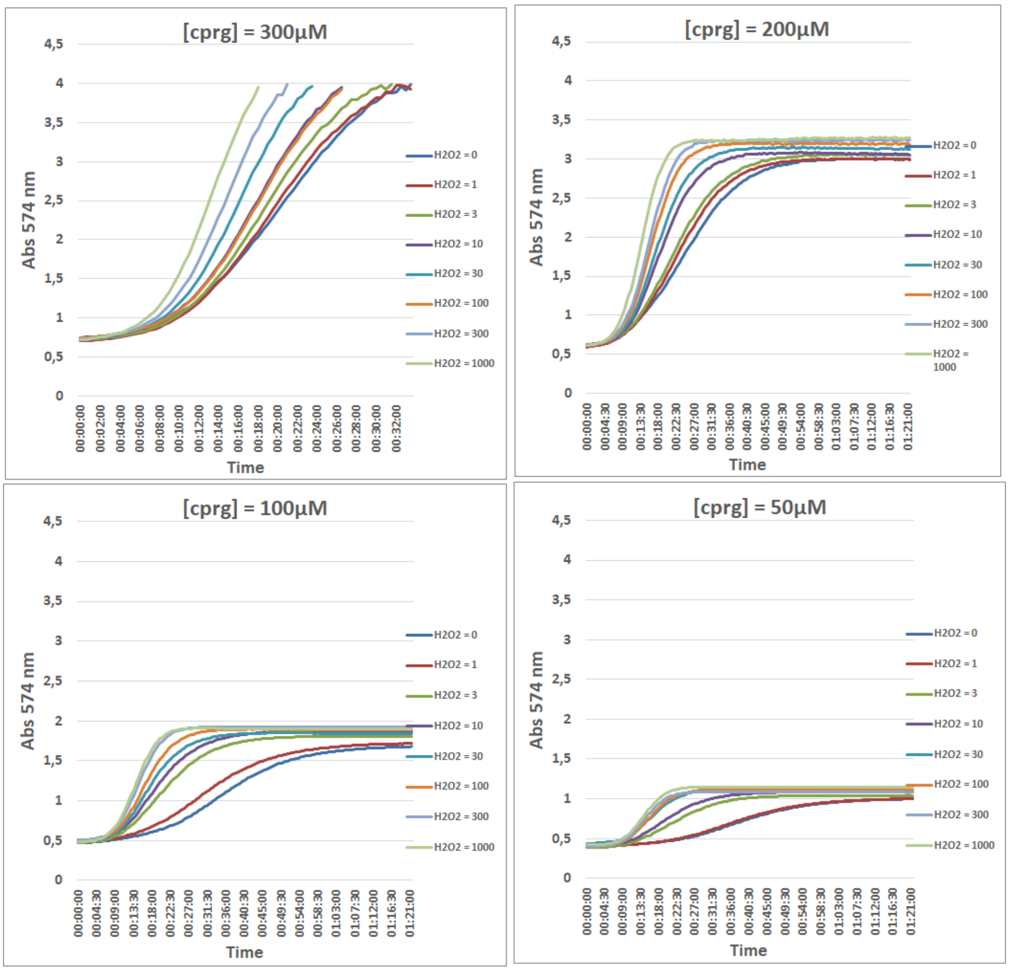


**B**

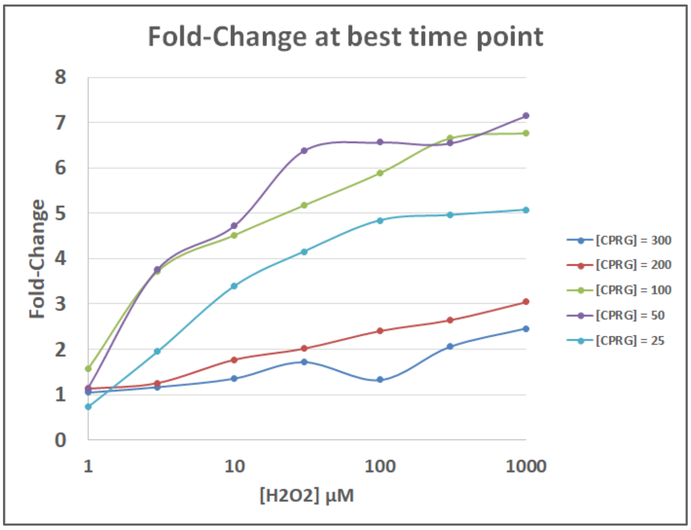


**Supplementary Figure S7: CPRG concentration optimization for colorimetric $H_2O_2$ biosensor (A)** 574 nm absorbance time course kinetic of colorimetric $H_2O_2$ biosensor with various initial concentration of CPRG **(B)** Blank subtracted $H_2O_2$ dose response curve of colorimetric biosensor with various initial concentration of CPRG

**Fig S8:**

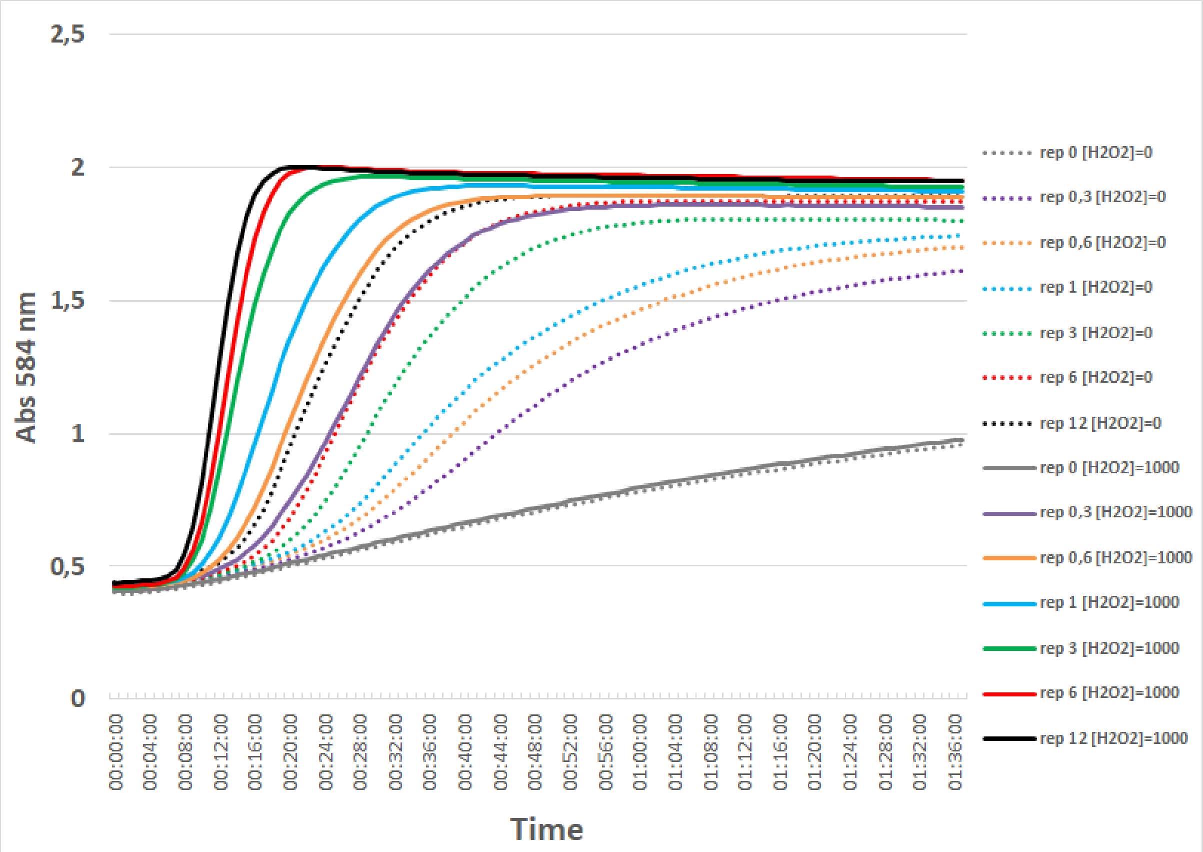


**Supplementary Figure S8: [pAhpC-LacZ DNA] concentration optimization for colorimetric $H_2O_2$ biosensor**

**Fig S9:**

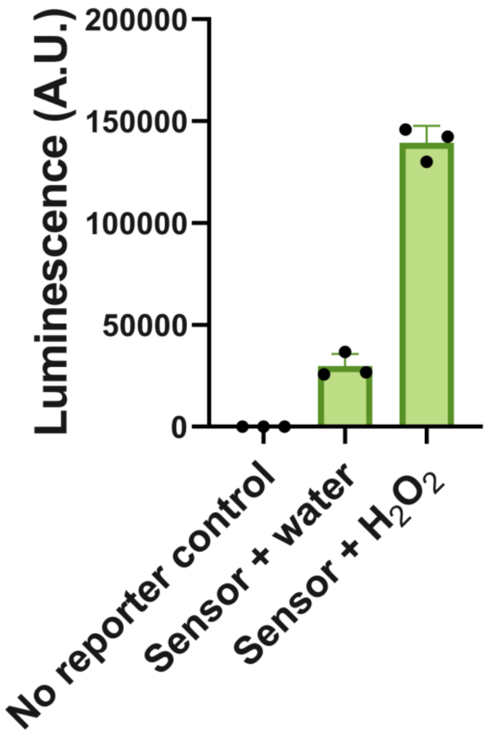


**Supplementary Figure S9: Luminescent sensor early evaluation.** [pAhpC-Luc DNA] was added at 0 or 6nM and [j23101-OxyR DNA] at 24 nM to 1h preincubated cell-free mix together with either water or 1mM $H_2O_2$. The reaction was incubated for 30 min at 37°C and 20µL of final reaction was added to 50 µL of Luciferin reagent mix before taking a luminescence point measurement. Dots represent individual values obtained.Error bars represent the standard deviation calculated from the 3 replicates.

**Fig S10:**

**A**

**B**

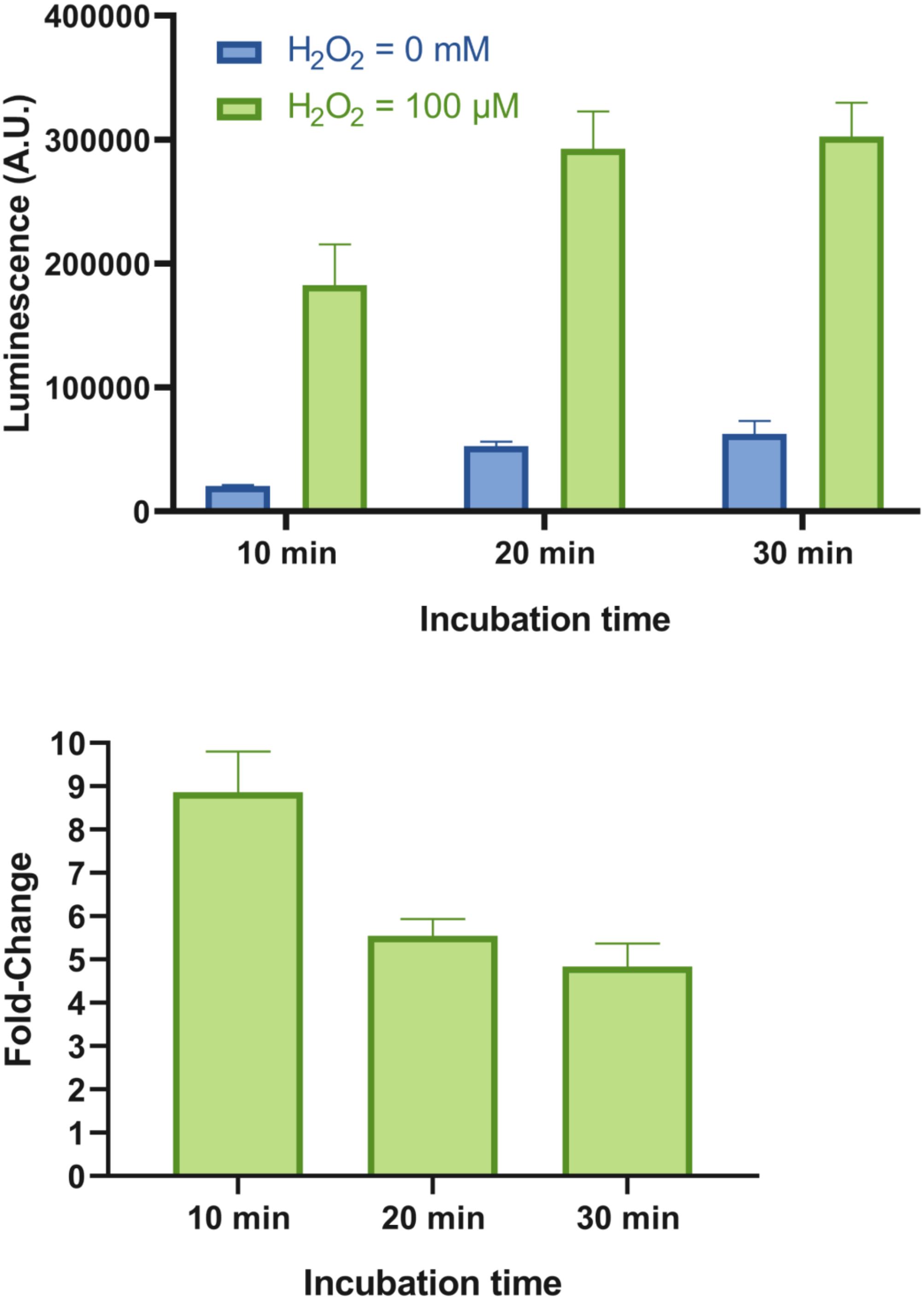


**Supplementary Figure S10: Luminescent sensor incubation time optimisation (A)** Luminescence absolute values measured in arbitrary units **(B)** Luminescence Fold-Change between induced and non induced condition

**Supplementary Table S1**

Chemicals identifiers used in the study

| **Compound** | **InChI** |
|---|---|
| S-lactate | InChI=1S/C3H6O3/c1-2(4)3(5)6/h2,4H,1H3,(H,5,6)/p-1/t2-/m0/s1 |
| choline | InChI=1S/C5H14NO/c1-6(2,3)4-5-7/h7H,4-5H2,1-3H3/q+1 |
| sarcosine | InChI=1S/C3H7NO2/c1-4-2-3(5)6/h4H,2H2,1H3,(H,5,6) |

**Supplementary Table S2**
Characteristics of enzymes used in the study

| **substrate** | **name** | **uniprot ID** | **organism** | **kinetic** | | **catalytic efficiency** |
|---|---|---|---|---|---|---|
| | | | | Km | Kcat | Kcat/Km |
| sarcosine | SoxA | P40859 | *Bacillus subtilis* | 141,6 mM source | 0.204 s-1 source | 1.441 s-1.M-1 source |
| choline | CodA | Q7X2H8 | *Arthrobacter globiformis* | 0,6 mM source | 13.4 s-1 source | 22000 s-1.M-1 source |
| lactate | lox | D4YFM2 | *Aerococcus viridans* | 0,5 mM source | 88 s-1 source | 176000 s-1.M-1 source |

**Supplementary Table S3**
Plasmids used in this study:

| Plasmid | Antibiotic Selection | Accession/ Catalog reference | Source |
|---|---|---|---|
| pBeast-J23101-RBS-OxyR | $Amp^R$ | | This work |
| pBeast-pAhpC-RBS-sfGFP | $Amp^R$ | | This work |
| pBeast-pOxyS-RBS-sfGFP | $Amp^R$ | | This work |
| pBeast-pKatG-RBS-sfGFP | $Amp^R$ | | This work |
| pBeast-pYjjZ-RBS-sfGFP | $Amp^R$ | | This work |
| pBeast-pZint-sfGFP | $Amp^R$ | | This work |
| pBeast-J23101-RBS-soxA | $Amp^R$ | | This work |
| pBeast-pT7-soxA | $Amp^R$ | | This work |
| pBeast-pT7-codA | $Amp^R$ | | This work |
| pBeast-pT7-lox | $Amp^R$ | | This work |
| pBeast-pAhpC-RBS-LacZ | $Amp^R$ | | This work |
| pBeast-pAhpC-RBS-Luc | $Amp^R$ | | This work |

**Supplementary Table S4**

DNA sequences for constructs used in this study:

| Plasmid/ DNA construct | Sequence (5'->3') |
|---|---|
| pBeast-J23101-RBS-OxyR | ATACTAGAGGATGACCCCATCTGTTTACAGCTAGCTCAGTCCTAG<br>GTATTATGCTAGCTAGTAGAGTCACACAGGAAAGTAGTAGATGAAT<br>ATTCGTGATCTTGAGTACCTGGTGGCATTGGCTGAACACCGCCAT<br>TTTCGGCGTGCGGCAGATTCCTGCCACGTTAGCCAGCCGACGCTT<br>AGCGGGCAAATTCGTAAGCTGGAAGATGAGCTGGGCGTGATGTT<br>GCTGGAGCGGACCAGCCGTAAAGTGTTGTTCACCCAGGCGGGAA<br>TGCTGCTGGTGGATCAGGCGCGTACCGTGCTGCGTGAGGTGAAA<br>GTCCTTAAAGAGATGGCAAGCCAGCAGGGCGAAACCATGTCCGG<br>ACCGCTGCACATTGGTTTGATTCCCACAGTTGGACCGTACCTGCT<br>ACCGCATATTATCCCTATGCTGCACCAGACCTTTCCAAAGCTGGAA<br>ATGTATCTGCATGAAGCACAGACCCACCAGTTACTGGCGCAACTG<br>GACAGCGGCAAACTCGATTGCGTGATCCTCGCGCTGGTGAAAGA<br>GAGCGAAGCATTCATTGAAGTGCCGTTGTTTGATGAGCCAATGTT<br>GCTGGCTATCTATGAAGATCACCCGTGGGCGAACCGCGAATGCGT<br>ACCGATGGCCGATCTGGCAGGGGAAAAACTGCTGATGCTGGAAG<br>ATGGTCACTGTTTGCGCGATCAGGCAATGGGTTTCTGTTTTGAAG<br>CCGGGGCGGATGAAGATACACACTTCCGCGCGACCAGCCTGGAA<br>ACTCTTCGCAACATGGTGGCGGCAGGTAGCGGGATCACTTTACTG<br>CCAGCGCTGGCTGTGCCGCCGGAGCGCAAACGCGATGGGGTTGT<br>TTATCTGCCGTGCATTAAGCCGGAACCACGCCGCACTATTGGCCT<br>GGTTTATCGTCCTGGCTCACCGCTGCGCAGCCGCTATGAGCAGCT<br>GGCGGAAGCCATCCGCGCAAGAATGGATGGCCATTTCGATAAAGT<br>TTTAAAACAGGCGGTTTAAactttatctgagaatagtcaatcttcggaaatcccaggtg<br>gcatgctaaaagtctcgtaaagcgttctatcaataacccgttggtgccaggcatcaaataaaacga<br>aaggctcagtcgaaagactgggcctttcgttttatctgttgtttgtcggtgaacgctctctactagagtc<br>acactggctcaccttcgggtgggcctttctgcgtttataccgtctcagaatcggccgtgaacaataaa<br>atagtttcggtattattgaccacttccgagtagaatcgtgcttcagtaagagtcgaccgatgcccttga<br>gagccttcaacccagtcagctccttccggtgggcgcggggcaTGACTATCGTCGCCG<br>CACTTATGACTGTCTTCTTTATCATGCAACTCGTAGGACAGGTGCC<br>GGCAGCGCTCTTCCGCTTCCTCGCTCACTGACTCGCTGCGCTCG<br>GTCGTTCGGCTGCGGCGAGCGGTATCAGCTCACTCAAAGGCGGT<br>AATACGGTTATCCACAGAATCAGGGGATAACGCAGGAAAGAACAT<br>GTGAGCAAAAGGCCAGCAAAAGGCCAGGAACCGTAAAAAGGCCG<br>CGTTGCTGGCGTTTTTCCATAGGCTCCGCCCCCCTGACGAGCATC<br>ACAAAAATCGACGCTCAAGTCAGAGGTGGCGAAACCCGACAGGA<br>CTATAAAGATACCAGGCGTTTCCCCCTGGAAGCTCCCTCGTGCGC<br>TCTCCTGTTCCGACCCTGCCGCTTACCGGATACCTGTCCGCCTTT<br>CTCCCTTCGGGAAGCGTGGCGCTTTCTCAATGCTCACGCTGTAGG<br>TATCTCAGTTCGGTGTAGGTCGTTCGCTCCAAGCTGGGCTGTGTG<br>CACGAACCCCCCGTTCAGCCCGACCGCTGCGCCTTATCCGGTAA<br>CTATCGTCTTGAGTCCAACCCGGTAAGACACGACTTATCGCCACT<br>GGCAGCAGCCACTGGTAACAGGATTAGCAGAGCGAGGTATGTAG<br>GCGGTGCTACAGAGTTCTTGAAGTGGTGGCCTAACTACGGCTACA<br>CTAGAAGGACAGTATTTGGTATCTGCGCTCTGCTGAAGCCAGTTA<br>CCTTCGGAAAAAGAGTTGGTAGCTCTTGATCCGGCAAACAAACCA<br>CCGCTGGTAGCGGTGGTTTTTTTGTTTGCAAGCAGCAGATTACGC |

| | |
|---|---|
| | GCAGAAAAAAAGGATCTCAAGAAGATCCTTTGATCTTTTCTACGGG<br>GTCTGACGCTCAGTGGAACGAAAACTCACGTTAAGGGATTTTGGT<br>CATGAGATTATCAAAAAGGATCTTCACCTAGATCCTTTTAAATTAAA<br>AATGAAGTTTTAAATCAATCTAAAGTATATATGAGTAAACTTGGTCT<br>GACAGTTACCAATGCTTAATCAGTGAGGCACCTATCTCAGCGATCT<br>GTCTATTTCGTTCATCCATAGTTGCCTGACTCCCCGTCGTGTAGAT<br>AACTACGATACGGGAGGGCTTACCATCTGGCCCCAGTGCTGCAAT<br>GATACCGCGAGACCCACGCTCACCGGCTCCAGATTTATCAGCAAT<br>AAACCAGCCAGCCGGAAGGGCCGAGCGCAGAAGTGGTCCTGCAA<br>CTTTATCCGCCTCCATCCAGTCTATTAATTGTTGCCGGGAAGCTAG<br>AGTAAGTAGTTCGCCAGTTAATAGTTTGCGCAACGTTGTTGCCATT<br>GCTACAGGCATCGTGGTGTCACGCTCGTCGTTTGGTATGGCTTCA<br>TTCAGCTCCGGTTCCCAACGATCAAGGCGAGTTACATGATCCCCC<br>ATGTTGTGCAAAAAAGCGGTTAGCTCCTTCGGTCCTCCGATCGTT<br>GTCAGAAGTAAGTTGGCCGCAGTGTTATCACTCATGGTTATGGCA<br>GCACTGCATAATTCTCTTACTGTCATGCCATCCGTAAGATGCTTTT<br>CTGTGACTGGTGAGTACTCAACCAAGTCATTCTGAGAATAGTGTAT<br>GCGGCGACCGAGTTGCTCTTGCCCGGCGTCAATACGGGATAATA<br>CCGCGCCACATAGCAGAACTTTAAAAGTGCTCATCATTGGAAAAC<br>GTTCTTCGGGGCGAAAACTCTCAAGGATCTTACCGCTGTTGAGAT<br>CCAGTTCGATGTAACCCACTCGTGCACCCAACTGATCTTCAGCAT<br>CTTTTACTTTCACCAGCGTTTCTGGGTGAGCAAAAACAGGAAGGC<br>AAAATGCCGCAAAAAAGGGAATAAGGGCGACACGGAAATGTTGAA<br>TACTCATACTCTTCCTTTTTCAATATTATTGAAGCATTTATCAGGGT<br>TATTGTCTCATGAGCGGATACATATTTGAATGTATTTAGAAAAATAA<br>ACAAATAGGGGTTCCGCGCACATTTCCCCGAAAAGTGCCACCTGA<br>CGTCTAAGAAACCATTATTATCATGACATTAACCTATAAAAATAGGC<br>GTATCACGAGGCCCTTTCGTCTTCAAGAATTCTGGCGAATCCTCTG<br>ACCAGCCAGAAAACGACCTTTCTGTGGTGAAACCGGATGCTGCAA<br>TTCAGAGCGGCAGCAAGTGGGGGACAGCAGAAGACCTGACCGCC<br>GCAGAGTGGATGTTTGACATGGTGAAGACTATCGCACCATCAGCC<br>AGAAAACCGAATTTTGCTGGGTGGGCTAACGATATCcgcctgatgcgtga<br>acgtgacggacgtaaccaccgcgacatgtgtgtgctgttccgctggctcggatacccttactctgttg<br>aaaacgaatagataggttaaggaacggttatttctgcgtagatctatcttacacagcatcacactgg<br>ctcaccttcgggtgggcctttctgcgtttatatactagagagagaatataaaaagccagattattaat<br>ccggctttttttattatttaggcaactgaaacgattcggatcctgtattactattctta |
| pBeast-pAhpC-<br>RBS-sfGFP | GCTTAGATCAGGTGATTGCCCTTTGTTTATGAGGGTGTTGTAATCC<br>ATGTCGTTGTTGCATTTGTAAGGGCAACACCTCAGCCTGCAGGCA<br>GGCACTGAAGATACCAAAGGGTAGTTCAGATTACACGGTCACCTG<br>GAAAGGGGGCCATTTTACTTTTTATCGCCGCTGGCGGTGCAAAGT<br>TCACAAAGTTGTCTTACGAAGGTTGTAAGGTAAAACTTATCGATTT<br>GATAATGGAAACGCATTAGCCGAATCGGCAAAAATTGGTTACCTTA<br>CATCTCATCGAAAACACGGAGGAAGTATAGATGTCTAGAGAAAGA<br>GGAGAAATACTAGatgcgtaaaggcgaagagctgttcactggtgtcgtccctattctggtg<br>gaactggatggtgatgtcaacggtcataagttttccgtgcgtggcgagggtgaaggtgacgcaact<br>aatggtaaactgacgctgaagttcatctgtactactggtaaactgccggtaccttggccgactctggt<br>aacgacgctgacttatggtgttcagtgctttgctcgttatccggaccatatgaagcagcatgacttctt<br>caagtccgccatgccggaaggctatgtgcaggaacgcacgatttcctttaaggatgacggcacgt<br>acaaaacgcgtgcggaagtgaaatttgaaggcgataccctggtaaaccgcattgagctgaaag<br>gcattgactttaaagaagacggcaatatcctgggccataagctggaatacaattttaacagccaca<br>atgtttacatcaccgccgataaacaaaaaaatggcattaaagcgaattttaaaattcgccacaacg<br>tggaggatggcagcgtgcagctggctgatcactaccagcaaaacactccaatcggtgatggtcct<br>gttctgctgccagacaatcactatctgagcacgcaaagcgttctgtctaaagatccgaacgagaa |

| | |
|---|---|
| | acgcgatcatatggttctgctggagttcgtaaccgcagcgggcatcacgcatggtatggatgaact<br>gtacaaatgatgaactttatctgagaatagtcaatcttcggaaatcccaggtggcatgctaaaagtc<br>tcgtaaagcgttctatcaataacccgttggtgccaggcatcaaataaaacgaaaggctcagtcga<br>aagactgggcctttcgttttatctgttgtttgtcggtgaacgctctctactagagtcacactggctcacct<br>tcgggtgggcctttctgcgtttataccgtctcagaatcggccgtgaacaataaaatagtttcggtattat<br>tgaccacttccgagtagaatcgtgcttcagtaagagtcgaccgatgcccttgagagccttcaaccc<br>agtcagctccttccggtgggcgcggggcaTGACTATCGTCGCCGCACTTATGAC<br>TGTCTTCTTTATCATGCAACTCGTAGGACAGGTGCCGGCAGCGCT<br>CTTCCGCTTCCTCGCTCACTGACTCGCTGCGCTCGGTCGTTCGGC<br>TGCGGCGAGCGGTATCAGCTCACTCAAAGGCGGTAATACGGTTAT<br>CCACAGAATCAGGGGATAACGCAGGAAAGAACATGTGAGCAAAAG<br>GCCAGCAAAAGGCCAGGAACCGTAAAAAGGCCGCGTTGCTGGCG<br>TTTTTCCATAGGCTCCGCCCCCCTGACGAGCATCACAAAAATCGA<br>CGCTCAAGTCAGAGGTGGCGAAACCCGACAGGACTATAAAGATAC<br>CAGGCGTTTCCCCCTGGAAGCTCCCTCGTGCGCTCTCCTGTTCCG<br>ACCCTGCCGCTTACCGGATACCTGTCCGCCTTTCTCCCTTCGGGA<br>AGCGTGGCGCTTTCTCAATGCTCACGCTGTAGGTATCTCAGTTCG<br>GTGTAGGTCGTTCGCTCCAAGCTGGGCTGTGTGCACGAACCCCC<br>CGTTCAGCCCGACCGCTGCGCCTTATCCGGTAACTATCGTCTTGA<br>GTCCAACCCGGTAAGACACGACTTATCGCCACTGGCAGCAGCCAC<br>TGGTAACAGGATTAGCAGAGCGAGGTATGTAGGCGGTGCTACAGA<br>GTTCTTGAAGTGGTGGCCTAACTACGGCTACACTAGAAGGACAGT<br>ATTTGGTATCTGCGCTCTGCTGAAGCCAGTTACCTTCGGAAAAAGA<br>GTTGGTAGCTCTTGATCCGGCAAACAAACCACCGCTGGTAGCGGT<br>GGTTTTTTTGTTTGCAAGCAGCAGATTACGCGCAGAAAAAAAGGAT<br>CTCAAGAAGATCCTTTGATCTTTTCTACGGGGTCTGACGCTCAGTG<br>GAACGAAAACTCACGTTAAGGGATTTTGGTCATGAGATTATCAAAA<br>AGGATCTTCACCTAGATCCTTTTAAATTAAAAATGAAGTTTTAAATC<br>AATCTAAAGTATATATGAGTAAACTTGGTCTGACAGTTACCAATGC<br>TTAATCAGTGAGGCACCTATCTCAGCGATCTGTCTATTTCGTTCAT<br>CCATAGTTGCCTGACTCCCCGTCGTGTAGATAACTACGATACGGG<br>AGGGCTTACCATCTGGCCCCAGTGCTGCAATGATACCGCGAGACC<br>CACGCTCACCGGCTCCAGATTTATCAGCAATAAACCAGCCAGCCG<br>GAAGGGCCGAGCGCAGAAGTGGTCCTGCAACTTTATCCGCCTCC<br>ATCCAGTCTATTAATTGTTGCCGGGAAGCTAGAGTAAGTAGTTCGC<br>CAGTTAATAGTTTGCGCAACGTTGTTGCCATTGCTACAGGCATCGT<br>GGTGTCACGCTCGTCGTTTGGTATGGCTTCATTCAGCTCCGGTTC<br>CCAACGATCAAGGCGAGTTACATGATCCCCCATGTTGTGCAAAAA<br>AGCGGTTAGCTCCTTCGGTCCTCCGATCGTTGTCAGAAGTAAGTT<br>GGCCGCAGTGTTATCACTCATGGTTATGGCAGCACTGCATAATTCT<br>CTTACTGTCATGCCATCCGTAAGATGCTTTTCTGTGACTGGTGAGT<br>ACTCAACCAAGTCATTCTGAGAATAGTGTATGCGGCGACCGAGTT<br>GCTCTTGCCCGGCGTCAATACGGGATAATACCGCGCCACATAGCA<br>GAACTTTAAAAGTGCTCATCATTGGAAAACGTTCTTCGGGGCGAAA<br>ACTCTCAAGGATCTTACCGCTGTTGAGATCCAGTTCGATGTAACCC<br>ACTCGTGCACCCAACTGATCTTCAGCATCTTTTACTTTCACCAGCG<br>TTTCTGGGTGAGCAAAAACAGGAAGGCAAAATGCCGCAAAAAAGG<br>GAATAAGGGCGACACGGAAATGTTGAATACTCATACTCTTCCTTTT<br>TCAATATTATTGAAGCATTTATCAGGGTTATTGTCTCATGAGCGGA<br>TACATATTTGAATGTATTTAGAAAAATAAACAAATAGGGGTTCCGC<br>GCACATTTCCCCGAAAAGTGCCACCTGACGTCTAAGAAACCATTAT<br>TATCATGACATTAACCTATAAAAATAGGCGTATCACGAGGCCCTTT<br>CGTCTTCAAGAATTCTGGCGAATCCTCTGACCAGCCAGAAAACGA<br>CCTTTCTGTGGTGAAACCGGATGCTGCAATTCAGAGCGGCAGCAA |

| | |
|---|---|
| | GTGGGGGACAGCAGAAGACCTGACCGCCGCAGAGTGGATGTTTG<br>ACATGGTGAAGACTATCGCACCATCAGCCAGAAAACCGAATTTTG<br>CTGGGTGGGCTAACGATATCcgcctgatgcgtgaacgtgacggacgtaaccaccg<br>cgacatgtgtgtgctgttccgctggctcggatacccttactctgttgaaaacgaatagataggttaag<br>gaacggttatttctgcgtagatctatcttacacagcatcacactggctcaccttcgggtgggcctttctg<br>cgtttatatactagagagagaatataaaaagccagattattaatccggctttttttattatttaggcaact<br>gaaacgattcggatcctgtattactattctta |
| pBeast-pOxyS-<br>RBS-sfGFP | TTCATTATCCATCCTCCATCGCCACGATAGTTCATGGCGATAGGTA<br>GAATAGCAATGAACGATTATCCCTATCAAGCATTCTGACTGAGCAT<br>TGCTCACATCTAGAGAAAGAGGAGAAATACTAGatgcgtaaaggcgaaga<br>gctgttcactggtgtcgtccctattctggtggaactggatggtgatgtcaacggtcataagttttccgtg<br>cgtggcgagggtgaaggtgacgcaactaatggtaaactgacgctgaagttcatctgtactactggt<br>aaactgccggtaccttggccgactctggtaacgacgctgacttatggtgttcagtgctttgctcgttat<br>ccggaccatatgaagcagcatgacttcttcaagtccgccatgccggaaggctatgtgcaggaac<br>gcacgatttcctttaaggatgacggcacgtacaaaacgcgtgcggaagtgaaatttgaaggcgat<br>accctggtaaaccgcattgagctgaaaggcattgactttaaagaagacggcaatatcctgggcca<br>taagctggaatacaattttaacagccacaatgtttacatcaccgccgataaacaaaaaaatggcat<br>taaagcgaattttaaaattcgccacaacgtggaggatggcagcgtgcagctggctgatcactacc<br>agcaaaacactccaatcggtgatggtcctgttctgctgccagacaatcactatctgagcacgcaaa<br>gcgttctgtctaaagatccgaacgagaaacgcgatcatatggttctgctggagttcgtaaccgcag<br>cgggcatcacgcatggtatggatgaactgtacaaatgatgaactttatctgagaatagtcaatcttc<br>ggaaatcccaggtggcatgctaaaagtctcgtaaagcgttctatcaataacccgttggtgccaggc<br>atcaaataaaacgaaaggctcagtcgaaagactgggcctttcgttttatctgttgtttgtcggtgaac<br>gctctctactagagtcacactggctcaccttcgggtgggcctttctgcgtttataccgtctcagaatcg<br>gccgtgaacaataaaatagtttcggtattattgaccacttccgagtagaatcgtgcttcagtaagagt<br>cgaccgatgcccttgagagccttcaacccagtcagctccttccggtgggcgcggggcaTGACT<br>ATCGTCGCCGCACTTATGACTGTCTTCTTTATCATGCAACTCGTAG<br>GACAGGTGCCGGCAGCGCTCTTCCGCTTCCTCGCTCACTGACTC<br>GCTGCGCTCGGTCGTTCGGCTGCGGCGAGCGGTATCAGCTCACT<br>CAAAGGCGGTAATACGGTTATCCACAGAATCAGGGGATAACGCAG<br>GAAAGAACATGTGAGCAAAAGGCCAGCAAAAGGCCAGGAACCGT<br>AAAAAGGCCGCGTTGCTGGCGTTTTTCCATAGGCTCCGCCCCCCT<br>GACGAGCATCACAAAAATCGACGCTCAAGTCAGAGGTGGCGAAAC<br>CCGACAGGACTATAAAGATACCAGGCGTTTCCCCCTGGAAGCTCC<br>CTCGTGCGCTCTCCTGTTCCGACCCTGCCGCTTACCGGATACCTG<br>TCCGCCTTTCTCCCTTCGGGAAGCGTGGCGCTTTCTCAATGCTCA<br>CGCTGTAGGTATCTCAGTTCGGTGTAGGTCGTTCGCTCCAAGCTG<br>GGCTGTGTGCACGAACCCCCCGTTCAGCCCGACCGCTGCGCCTT<br>ATCCGGTAACTATCGTCTTGAGTCCAACCCGGTAAGACACGACTT<br>ATCGCCACTGGCAGCAGCCACTGGTAACAGGATTAGCAGAGCGA<br>GGTATGTAGGCGGTGCTACAGAGTTCTTGAAGTGGTGGCCTAACT<br>ACGGCTACACTAGAAGGACAGTATTTGGTATCTGCGCTCTGCTGA<br>AGCCAGTTACCTTCGGAAAAAGAGTTGGTAGCTCTTGATCCGGCA<br>AACAAACCACCGCTGGTAGCGGTGGTTTTTTTGTTTGCAAGCAGC<br>AGATTACGCGCAGAAAAAAAGGATCTCAAGAAGATCCTTTGATCTT<br>TTCTACGGGGTCTGACGCTCAGTGGAACGAAAACTCACGTTAAGG<br>GATTTTGGTCATGAGATTATCAAAAAGGATCTTCACCTAGATCCTTT<br>TAAATTAAAAATGAAGTTTTAAATCAATCTAAAGTATATATGAGTAA<br>ACTTGGTCTGACAGTTACCAATGCTTAATCAGTGAGGCACCTATCT<br>CAGCGATCTGTCTATTTCGTTCATCCATAGTTGCCTGACTCCCCGT<br>CGTGTAGATAACTACGATACGGGAGGGCTTACCATCTGGCCCCAG<br>TGCTGCAATGATACCGCGAGACCCACGCTCACCGGCTCCAGATTT<br>ATCAGCAATAAACCAGCCAGCCGGAAGGGCCGAGCGCAGAAGTG |

| | |
|---|---|
| | GTCCTGCAACTTTATCCGCCTCCATCCAGTCTATTAATTGTTGCCG<br>GGAAGCTAGAGTAAGTAGTTCGCCAGTTAATAGTTTGCGCAACGT<br>TGTTGCCATTGCTACAGGCATCGTGGTGTCACGCTCGTCGTTTGG<br>TATGGCTTCATTCAGCTCCGGTTCCCAACGATCAAGGCGAGTTAC<br>ATGATCCCCCATGTTGTGCAAAAAAGCGGTTAGCTCCTTCGGTCC<br>TCCGATCGTTGTCAGAAGTAAGTTGGCCGCAGTGTTATCACTCAT<br>GGTTATGGCAGCACTGCATAATTCTCTTACTGTCATGCCATCCGTA<br>AGATGCTTTTCTGTGACTGGTGAGTACTCAACCAAGTCATTCTGAG<br>AATAGTGTATGCGGCGACCGAGTTGCTCTTGCCCGGCGTCAATAC<br>GGGATAATACCGCGCCACATAGCAGAACTTTAAAAGTGCTCATCAT<br>TGGAAAACGTTCTTCGGGGCGAAAACTCTCAAGGATCTTACCGCT<br>GTTGAGATCCAGTTCGATGTAACCCACTCGTGCACCCAACTGATC<br>TTCAGCATCTTTTACTTTCACCAGCGTTTCTGGGTGAGCAAAAACA<br>GGAAGGCAAAATGCCGCAAAAAAGGGAATAAGGGCGACACGGAA<br>ATGTTGAATACTCATACTCTTCCTTTTTCAATATTATTGAAGCATTTA<br>TCAGGGTTATTGTCTCATGAGCGGATACATATTTGAATGTATTTAG<br>AAAAATAAACAAATAGGGGTTCCGCGCACATTTCCCCGAAAAGTG<br>CCACCTGACGTCTAAGAAACCATTATTATCATGACATTAACCTATAA<br>AAATAGGCGTATCACGAGGCCCTTTCGTCTTCAAGAATTCTGGCG<br>AATCCTCTGACCAGCCAGAAAACGACCTTTCTGTGGTGAAACCGG<br>ATGCTGCAATTCAGAGCGGCAGCAAGTGGGGGACAGCAGAAGAC<br>CTGACCGCCGCAGAGTGGATGTTTGACATGGTGAAGACTATCGCA<br>CCATCAGCCAGAAAACCGAATTTTGCTGGGTGGGCTAACGATATC<br>cgcctgatgcgtgaacgtgacggacgtaaccaccgcgacatgtgtgtgctgttccgctggctcgga<br>taccottactctgttgaaaacgaatagataggttaaggaacggttatttctgcgtagatctatcttaca<br>cagcatcacactggctcaccttcgggtgggcctttctgcgtttatatactagagagagaatataaaa<br>agccagattattaatccggctttttattatttaggcaactgaaacgattcggatcctgtattactattctt<br>a |
| pBeast-pKatG-RBS-sfGFP | TGTGGCTTTTATGAAAATCACACAGTGATCACAAATTTTAAACAGA<br>GCACAAAATGCTGCCTCGAAATGAGGGCGGGAAAATAAGGTTATC<br>AGCCTTGTTTTCTCCCTCATTACTTGAAGGATATGAAGCTAAAACC<br>CTTTTTTATAAAGCATTTGTCCGAATTCGGACATAATCAAAAAAGCT<br>TAATTAAGATCAATTTGATCTACATCTCTTTAACCAACAATATGTAA<br>GATCTCAACTATCGCATCCGTGGATTAATTCAATTATAACTTCTCTC<br>TAACGCTGTGTATCGTAACGGTAACACTGTAGAGGGGAGCACATT<br>GATGTCTAGAGAAAGAGGAGAAATACTAGatgcgtaaaggcgaagagctgtt<br>cactggtgtcgtccctattctggtggaactggatggtgatgtcaacggtcataagttttccgtgcgtgg<br>cgagggtgaaggtgacgcaactaatggtaaactgacgctgaagttcatctgtactactggtaaact<br>gccggtaccttggccgactctggtaacgacgctgacttatggtgttcagtgctttgctcgttatccgga<br>ccatatgaagcagcatgacttcttcaagtccgccatgccggaaggctatgtgcaggaacgcacg<br>atttcctttaaggatgacggcacgtacaaaacgcgtgcggaagtgaaatttgaaggcgataccct<br>ggtaaaccgcattgagctgaaaggcattgactttaaagaagacggcaatatcctgggccataag<br>ctggaatacaattttaacagccacaatgtttacatcaccgccgataaacaaaaaaatggcattaaa<br>gcgaattttaaaattcgccacaacgtggaggatggcagcgtgcagctggctgatcactaccagca<br>aaacactccaatcggtgatggtcctgttctgctgccagacaatcactatctgagcacgcaaagcgtt<br>ctgtctaaagatccgaacgagaaacgcgatcatatggttctgctggagttcgtaaccgcagcggg<br>catcacgcatggtatggatgaactgtacaaatgatgaactttatctgagaatagtcaatcttcggaa<br>atcccaggtggcatgctaaaagtctcgtaaagcgttctatcaataacccgttggtgccaggcatca<br>aataaaacgaaaggctcagtcgaaagactgggcctttcgttttatctgttgtttgtcggtgaacgctct<br>ctactagagtcacactggctcaccttcgggtgggcctttctgcgtttataccgtctcagaatcggccgt<br>gaacaataaaatagtttcggtattattgaccacttccgagtagaatcgtgcttcagtaagagtcgac<br>cgatgcccttgagagccttcaacccagtcagctccttccggtgggcgcggggcaTGACTATC<br>GTCGCCGCACTTATGACTGTCTTCTTTATCATGCAACTCGTAGGAC |

| | |
|---|---|
| | AGGTGCCGGCAGCGCTCTTCCGCTTCCTCGCTCACTGACTCGCTG<br>CGCTCGGTCGTTCGGCTGCGGCGAGCGGTATCAGCTCACTCAAA<br>GGCGGTAATACGGTTATCCACAGAATCAGGGGATAACGCAGGAAA<br>GAACATGTGAGCAAAAGGCCAGCAAAAGGCCAGGAACCGTAAAAA<br>GGCCGCGTTGCTGGCGTTTTTCCATAGGCTCCGCCCCCCTGACG<br>AGCATCACAAAAATCGACGCTCAAGTCAGAGGTGGCGAAACCCGA<br>CAGGACTATAAAGATACCAGGCGTTTCCCCCTGGAAGCTCCCTCG<br>TGCGCTCTCCTGTTCCGACCCTGCCGCTTACCGGATACCTGTCCG<br>CCTTTCTCCCTTCGGGAAGCGTGGCGCTTTCTCAATGCTCACGCT<br>GTAGGTATCTCAGTTCGGTGTAGGTCGTTCGCTCCAAGCTGGGCT<br>GTGTGCACGAACCCCCCGTTCAGCCCGACCGCTGCGCCTTATCC<br>GGTAACTATCGTCTTGAGTCCAACCCGGTAAGACACGACTTATCG<br>CCACTGGCAGCAGCCACTGGTAACAGGATTAGCAGAGCGAGGTA<br>TGTAGGCGGTGCTACAGAGTTCTTGAAGTGGTGGCCTAACTACGG<br>CTACACTAGAAGGACAGTATTTGGTATCTGCGCTCTGCTGAAGCC<br>AGTTACCTTCGGAAAAAGAGTTGGTAGCTCTTGATCCGGCAAACA<br>AACCACCGCTGGTAGCGGTGGTTTTTTTGTTTGCAAGCAGCAGAT<br>TACGCGCAGAAAAAAAGGATCTCAAGAAGATCCTTTGATCTTTTCT<br>ACGGGGTCTGACGCTCAGTGGAACGAAAACTCACGTTAAGGGATT<br>TTGGTCATGAGATTATCAAAAAGGATCTTCACCTAGATCCTTTTAAA<br>TTAAAAATGAAGTTTTAAATCAATCTAAAGTATATATGAGTAAACTT<br>GGTCTGACAGTTACCAATGCTTAATCAGTGAGGCACCTATCTCAG<br>CGATCTGTCTATTTCGTTCATCCATAGTTGCCTGACTCCCCGTCGT<br>GTAGATAACTACGATACGGGAGGGCTTACCATCTGGCCCCAGTGC<br>TGCAATGATACCGCGAGACCCACGCTCACCGGCTCCAGATTTATC<br>AGCAATAAACCAGCCAGCCGGAAGGGCCGAGCGCAGAAGTGGTC<br>CTGCAACTTTATCCGCCTCCATCCAGTCTATTAATTGTTGCCGGGA<br>AGCTAGAGTAAGTAGTTCGCCAGTTAATAGTTTGCGCAACGTTGTT<br>GCCATTGCTACAGGCATCGTGGTGTCACGCTCGTCGTTTGGTATG<br>GCTTCATTCAGCTCCGGTTCCCAACGATCAAGGCGAGTTACATGA<br>TCCCCCATGTTGTGCAAAAAAGCGGTTAGCTCCTTCGGTCCTCCG<br>ATCGTTGTCAGAAGTAAGTTGGCCGCAGTGTTATCACTCATGGTTA<br>TGGCAGCACTGCATAATTCTCTTACTGTCATGCCATCCGTAAGATG<br>CTTTTCTGTGACTGGTGAGTACTCAACCAAGTCATTCTGAGAATAG<br>TGTATGCGGCGACCGAGTTGCTCTTGCCCGGCGTCAATACGGGAT<br>AATACCGCGCCACATAGCAGAACTTTAAAAGTGCTCATCATTGGAA<br>AACGTTCTTCGGGGCGAAAACTCTCAAGGATCTTACCGCTGTTGA<br>GATCCAGTTCGATGTAACCCACTCGTGCACCCAACTGATCTTCAG<br>CATCTTTTACTTTCACCAGCGTTTCTGGGTGAGCAAAAACAGGAAG<br>GCAAAATGCCGCAAAAAAGGGAATAAGGGCGACACGGAAATGTTG<br>AATACTCATACTCTTCCTTTTTCAATATTATTGAAGCATTTATCAGG<br>GTTATTGTCTCATGAGCGGATACATATTTGAATGTATTTAGAAAAAT<br>AAACAAATAGGGGTTCCGCGCACATTTCCCCGAAAAGTGCCACCT<br>GACGTCTAAGAAACCATTATTATCATGACATTAACCTATAAAAATAG<br>GCGTATCACGAGGCCCTTTCGTCTTCAAGAATTCTGGCGAATCCT<br>CTGACCAGCCAGAAAACGACCTTTCTGTGGTGAAACCGGATGCTG<br>CAATTCAGAGCGGCAGCAAGTGGGGGACAGCAGAAGACCTGACC<br>GCCGCAGAGTGGATGTTTGACATGGTGAAGACTATCGCACCATCA<br>GCCAGAAAACCGAATTTTGCTGGGTGGGCTAACGATATCcgcctgatg<br>cgtgaacgtgacggacgtaaccaccgcgacatgtgtgtgctgttccgctggctcggataccccttact<br>ctgttgaaaacgaatagataggttaaggaacggttatttctgcgtagatctatcttacacagcatcac<br>actggctcaccttcgggtgggcctttctgcgtttatatactagagagagagaatataaaaagccagatt<br>attaatccggctttttttattatttaggcaactgaaacgattcggatcctgtattactattctta |

| | |
|---|---|
| pBeast-pYjjZ-RBS-sfGFP | catagcgataggtttacccgatagcaagggatttatctggcttgcaaatgataaaaattatcatatga<br>tattggttatcattatcaatgaaagagatgaaatcatgcgtaaaggcgaagagctgttcactggtgtc<br>gtccctattctggtggaactggatggtgatgtcaacggtcataagttttccgtgcgtggcgagggtga<br>aggtgacgcaactaatggtaaactgacgctgaagttcatctgtactactggtaaactgccggtacct<br>tggccgactctggtaacgacgctgacttatggtgttcagtgctttgctcgttatccggaccatatgaag<br>cagcatgacttcttcaagtccgccatgccggaaggctatgtgcaggaacgcacgatttcctttaag<br>gatgacggcacgtacaaaacgcgtgcggaagtgaaatttgaaggcgataccctggtaaaccgc<br>attgagctgaaaggcattgactttaaagaagacggcaatatcctgggccataagctggaatacaa<br>ttttaacagccacaatgtttacatcaccgccgataaacaaaaaaatggcattaaagcgaattttaa<br>aattcgccacaacgtggaggatggcagcgtgcagctggctgatcactaccagcaaaacactcca<br>atcggtgatggtcctgttctgctgccagacaatcactatctgagcacgcaaagcgttctgtctaaag<br>atccgaacgagaaacgcgatcatatggttctgctggagttcgtaaccgcagcgggcatcacgcat<br>ggtatggatgaactgtacaaatgatgaactttatctgagaatagtcaatcttcggaaatcccaggtg<br>gcatgctaaaagtctcgtaaagcgttctatcaataacccgttggtgccaggcatcaaataaaacga<br>aaggctcagtcgaaagactgggcctttcgttttatctgttgtttgtcggtgaacgctctctactagagtc<br>acactggctcaccttcgggtgggcctttctgcgtttataccgtctcagaatcggccgtgaacaataaa<br>atagtttcggtattattgaccacttccgagtagaatcgtgcttcagtaagagtcgaccgatgcccttga<br>gagccttcaacccagtcagctccttccggtgggcgcggggcaTGACTATCGTCGCCG<br>CACTTATGACTGTCTTCTTTATCATGCAACTCGTAGGACAGGTGCC<br>GGCAGCGCTCTTCCGCTTCCTCGCTCACTGACTCGCTGCGCTCG<br>GTCGTTCGGCTGCGGCGAGCGGTATCAGCTCACTCAAAGGCGGT<br>AATACGGTTATCCACAGAATCAGGGGATAACGCAGGAAAGAACAT<br>GTGAGCAAAAGGCCAGCAAAAGGCCAGGAACCGTAAAAAGGCCG<br>CGTTGCTGGCGTTTTTCCATAGGCTCCGCCCCCCTGACGAGCATC<br>ACAAAAATCGACGCTCAAGTCAGAGGTGGCGAAACCCGACAGGA<br>CTATAAAGATACCAGGCGTTTCCCCCTGGAAGCTCCCTCGTGCGC<br>TCTCCTGTTCCGACCCTGCCGCTTACCGGATACCTGTCCGCCTTT<br>CTCCCTTCGGGAAGCGTGGCGCTTTCTCAATGCTCACGCTGTAGG<br>TATCTCAGTTCGGTGTAGGTCGTTCGCTCCAAGCTGGGCTGTGTG<br>CACGAACCCCCCGTTCAGCCCGACCGCTGCGCCTTATCCGGTAA<br>CTATCGTCTTGAGTCCAACCCGGTAAGACACGACTTATCGCCACT<br>GGCAGCAGCCACTGGTAACAGGATTAGCAGAGCGAGGTATGTAG<br>GCGGTGCTACAGAGTTCTTGAAGTGGTGGCCTAACTACGGCTACA<br>CTAGAAGGACAGTATTTGGTATCTGCGCTCTGCTGAAGCCAGTTA<br>CCTTCGGAAAAAGAGTTGGTAGCTCTTGATCCGGCAAACAAACCA<br>CCGCTGGTAGCGGTGGTTTTTTTGTTTGCAAGCAGCAGATTACGC<br>GCAGAAAAAAAGGATCTCAAGAAGATCCTTTGATCTTTTCTACGGG<br>GTCTGACGCTCAGTGGAACGAAAACTCACGTTAAGGGATTTTGGT<br>CATGAGATTATCAAAAAGGATCTTCACCTAGATCCTTTTAAATTAAA<br>AATGAAGTTTTAAATCAATCTAAAGTATATATGAGTAAACTTGGTCT<br>GACAGTTACCAATGCTTAATCAGTGAGGCACCTATCTCAGCGATCT<br>GTCTATTTCGTTCATCCATAGTTGCCTGACTCCCCGTCGTGTAGAT<br>AACTACGATACGGGAGGGCTTACCATCTGGCCCCAGTGCTGCAAT<br>GATACCGCGAGACCCACGCTCACCGGCTCCAGATTTATCAGCAAT<br>AAACCAGCCAGCCGGAAGGGCCGAGCGCAGAAGTGGTCCTGCAA<br>CTTTATCCGCCTCCATCCAGTCTATTAATTGTTGCCGGGAAGCTAG<br>AGTAAGTAGTTCGCCAGTTAATAGTTTGCGCAACGTTGTTGCCATT<br>GCTACAGGCATCGTGGTGTCACGCTCGTCGTTTGGTATGGCTTCA<br>TTCAGCTCCGGTTCCCAACGATCAAGGCGAGTTACATGATCCCCC<br>ATGTTGTGCAAAAAAGCGGTTAGCTCCTTCGGTCCTCCGATCGTT<br>GTCAGAAGTAAGTTGGCCGCAGTGTTATCACTCATGGTTATGGCA<br>GCACTGCATAATTCTCTTACTGTCATGCCATCCGTAAGATGCTTTT<br>CTGTGACTGGTGAGTACTCAACCAAGTCATTCTGAGAATAGTGTAT<br>GCGGCGACCGAGTTGCTCTTGCCCGGCGTCAATACGGGATAATA |

| | |
|---|---|
| | CCGCGCCACATAGCAGAACTTTAAAAGTGCTCATCATTGGAAAAC<br>GTTCTTCGGGGCGAAAACTCTCAAGGATCTTACCGCTGTTGAGAT<br>CCAGTTCGATGTAACCCACTCGTGCACCCAACTGATCTTCAGCAT<br>CTTTTACTTTCACCAGCGTTTCTGGGTGAGCAAAAACAGGAAGGC<br>AAAATGCCGCAAAAAAGGGAATAAGGGCGACACGGAAATGTTGAA<br>TACTCATACTCTTCCTTTTTCAATATTATTGAAGCATTTATCAGGGT<br>TATTGTCTCATGAGCGGATACATATTTGAATGTATTTAGAAAAATAA<br>ACAAATAGGGGTTCCGCGCACATTTCCCCGAAAAGTGCCACCTGA<br>CGTCTAAGAAACCATTATTATCATGACATTAACCTATAAAAATAGGC<br>GTATCACGAGGCCCTTTCGTCTTCAAGAATTCTGGCGAATCCTCTG<br>ACCAGCCAGAAAACGACCTTTCTGTGGTGAAACCGGATGCTGCAA<br>TTCAGAGCGGCAGCAAGTGGGGGACAGCAGAAGACCTGACCGCC<br>GCAGAGTGGATGTTTGACATGGTGAAGACTATCGCACCATCAGCC<br>AGAAAACCGAATTTTGCTGGGTGGGCTAACGATATCcgcctgatgcgtga<br>acgtgacggacgtaaccaccgcgacatgtgtgtgctgttccgctggctcggataccсttactctgttg<br>aaaacgaatagataggttaaggaacggttatttctgcgtagatctatcttacacagcatcacactgg<br>ctcaccttcgggtgggcctttctgcgtttatatactagagagagaatataaaaagccagattattaat<br>ccggctttttttattatttaggcaactgaaacgattcggatcctgtattactattctta |
| pBeast-pZint-sfGFP | tgagaaagccatgctctcgtttcctaagagttgttgcattttgctatatgttacaatataacattacacat<br>catatacattaactctggaggaaactgttatgcgtaaaggcgaagagctgttcactggtgtcgtccct<br>attctggtggaactggatggtgatgtcaacggtcataagttttccgtgcgtggcgagggtgaaggtg<br>acgcaactaatggtaaactgacgctgaagttcatctgtactactggtaaactgccggtaccttggcc<br>gactctggtaacgacgctgacttatggtgttcagtgctttgctcgttatccggaccatatgaagcagc<br>atgacttcttcaagtccgccatgccggaaggctatgtgcaggaacgcacgatttcctttaaggatga<br>cggcacgtacaaaacgcgtgcggaagtgaaatttgaaggcgataccctggtaaaccgcattgag<br>ctgaaaggcattgactttaaagaagacggcaatatcctgggccataagctggaatacaattttaac<br>agccacaatgtttacatcaccgccgataaacaaaaaaatggcattaaagcgaattttaaaattcgc<br>cacaacgtggaggatggcagcgtgcagctggctgatcactaccagcaaaacactccaatcggt<br>gatggtcctgttctgctgccagacaatcactatctgagcacgcaaagcgttctgtctaaagatccga<br>acgagaaacgcgatcatatggttctgctggagttcgtaaccgcagcgggcatcacgcatggtatg<br>gatgaactgtacaaatgatgaactttatctgagaatagtcaatcttcggaaatcccaggtggcatgc<br>taaaagtctcgtaaagcgttctatcaataacccgttggtgccaggcatcaaataaaacgaaaggct<br>cagtcgaaagactgggcctttcgttttatctgttgtttgtcggtgaacgctctctactagagtcacactg<br>gctcaccttcgggtgggcctttctgcgtttataccgtctcagaatcggccgtgaacaataaaatagttt<br>cggtattattgaccacttccgagtagaatcgtgcttcagtaagagtcgaccgatgcccttgagagcc<br>ttcaacccagtcagctccttccggtgggcgcggggcaTGACTATCGTCGCCGCACTT<br>ATGACTGTCTTCTTTATCATGCAACTCGTAGGACAGGTGCCGGCA<br>GCGCTCTTCCGCTTCCTCGCTCACTGACTCGCTGCGCTCGGTCGT<br>TCGGCTGCGGCGAGCGGTATCAGCTCACTCAAAGGCGGTAATAC<br>GGTTATCCACAGAATCAGGGGATAACGCAGGAAAGAACATGTGAG<br>CAAAAGGCCAGCAAAAGGCCAGGAACCGTAAAAAGGCCGCGTTG<br>CTGGCGTTTTTCCATAGGCTCCGCCCCCCTGACGAGCATCACAAA<br>AATCGACGCTCAAGTCAGAGGTGGCGAAACCCGACAGGACTATAA<br>AGATACCAGGCGTTTCCCCCTGGAAGCTCCCTCGTGCGCTCTCCT<br>GTTCCGACCCTGCCGCTTACCGGATACCTGTCCGCCTTTCTCCCT<br>TCGGGAAGCGTGGCGCTTTCTCAATGCTCACGCTGTAGGTATCTC<br>AGTTCGGTGTAGGTCGTTCGCTCCAAGCTGGGCTGTGTGCACGAA<br>CCCCCCGTTCAGCCCGACCGCTGCGCCTTATCCGGTAACTATCGT<br>CTTGAGTCCAACCCGGTAAGACACGACTTATCGCCACTGGCAGCA<br>GCCACTGGTAACAGGATTAGCAGAGCGAGGTATGTAGGCGGTGC<br>TACAGAGTTCTTGAAGTGGTGGCCTAACTACGGCTACACTAGAAG<br>GACAGTATTTGGTATCTGCGCTCTGCTGAAGCCAGTTACCTTCGG<br>AAAAAGAGTTGGTAGCTCTTGATCCGGCAAACAAACCACCGCTGG |

| | |
|---|---|
| | TAGCGGTGGTTTTTTTGTTTGCAAGCAGCAGATTACGCGCAGAAAA<br>AAAGGATCTCAAGAAGATCCTTTGATCTTTTCTACGGGGTCTGACG<br>CTCAGTGGAACGAAAACTCACGTTAAGGGATTTTGGTCATGAGATT<br>ATCAAAAAGGATCTTCACCTAGATCCTTTTAAATTAAAAATGAAGTT<br>TTAAATCAATCTAAAGTATATATGAGTAAACTTGGTCTGACAGTTAC<br>CAATGCTTAATCAGTGAGGCACCTATCTCAGCGATCTGTCTATTTC<br>GTTCATCCATAGTTGCCTGACTCCCCGTCGTGTAGATAACTACGAT<br>ACGGGAGGGCTTACCATCTGGCCCCAGTGCTGCAATGATACCGC<br>GAGACCCACGCTCACCGGCTCCAGATTTATCAGCAATAAACCAGC<br>CAGCCGGAAGGGCCGAGCGCAGAAGTGGTCCTGCAACTTTATCC<br>GCCTCCATCCAGTCTATTAATTGTTGCCGGGAAGCTAGAGTAAGTA<br>GTTCGCCAGTTAATAGTTTGCGCAACGTTGTTGCCATTGCTACAGG<br>CATCGTGGTGTCACGCTCGTCGTTTGGTATGGCTTCATTCAGCTC<br>CGGTTCCCAACGATCAAGGCGAGTTACATGATCCCCCATGTTGTG<br>CAAAAAAGCGGTTAGCTCCTTCGGTCCTCCGATCGTTGTCAGAAG<br>TAAGTTGGCCGCAGTGTTATCACTCATGGTTATGGCAGCACTGCA<br>TAATTCTCTTACTGTCATGCCATCCGTAAGATGCTTTTCTGTGACT<br>GGTGAGTACTCAACCAAGTCATTCTGAGAATAGTGTATGCGGCGA<br>CCGAGTTGCTCTTGCCCGGCGTCAATACGGGATAATACCGCGCCA<br>CATAGCAGAACTTTAAAAGTGCTCATCATTGGAAAACGTTCTTCGG<br>GGCGAAAACTCTCAAGGATCTTACCGCTGTTGAGATCCAGTTCGA<br>TGTAACCCACTCGTGCACCCAACTGATCTTCAGCATCTTTTACTTT<br>CACCAGCGTTTCTGGGTGAGCAAAAACAGGAAGGCAAAATGCCGC<br>AAAAAAGGGAATAAGGGCGACACGGAAATGTTGAATACTCATACT<br>CTTCCTTTTTCAATATTATTGAAGCATTTATCAGGGTTATTGTCTCA<br>TGAGCGGATACATATTTGAATGTATTTAGAAAAATAAACAAATAGG<br>GGTTCCGCGCACATTTCCCCGAAAAGTGCCACCTGACGTCTAAGA<br>AACCATTATTATCATGACATTAACCTATAAAAATAGGCGTATCACGA<br>GGCCCTTTCGTCTTCAAGAATTCTGGCGAATCCTCTGACCAGCCA<br>GAAAACGACCTTTCTGTGGTGAAACCGGATGCTGCAATTCAGAGC<br>GGCAGCAAGTGGGGGACAGCAGAAGACCTGACCGCCGCAGAGT<br>GGATGTTTGACATGGTGAAGACTATCGCACCATCAGCCAGAAAAC<br>CGAATTTTGCTGGGTGGGCTAACGATATCcgcctgatgcgtgaacgtgacgg<br>acgtaaccaccgcgacatgtgtgtgctgttccgctggctcggatacccttactctgttgaaaacgaat<br>agataggttaaggaacggttatttctgcgtagatctatcttacacagcatcacactggctcaccttcg<br>ggtgggcctttctgcgtttatatactagagagagaatataaaaagccagattattaatccggctttttta<br>ttatttaggcaactgaaacgattcggatcctgtattactattctta |
| pBeast-J23101-RBS-soxA | AGGAATACTAGAGGATGACCCCATCTGTTTACAGCTAGCTCAGTC<br>CTAGGTATTATGCTAGCTAGTAGAGTCACACAGGAAAGTAGTAGAT<br>GTCTACCCACTTCGACGTTATCGTTGTTGGTGCTGGTTCTATGGGT<br>ATGGCTGCTGGTTACCAGCTGGCTAAACAGGGTGTTAAAACCCTG<br>CTGGTTGACGCTTTCGACCCGCCGCACACCAACGGTTCTCACCAC<br>GGTGACACCCGTATCATCCGTCACGCTTACGGTGAAGGTCGTGAA<br>TACGTTCCGCTGGcTCTGCGTTCTCAGGAACTGTGGTACGAACTG<br>GAAAAAGAAACCCACCACAAAATCTTCACCAAGACCGGTGTTCTG<br>GTCTTCGGGCCGAAAGGTGAATCTGCTTTCGTTGCTGAAACCATG<br>GAAGCTGCTAAAGAACACTCTCTGACCGTTGACCTGCTGGAAGGT<br>GACGAAATCAACAAACGTTGGCCGGGTATCACCGTTCCGGAAAAC<br>TACAACGCTATCTTCGAACCGAACAGCGGTGTTCTGTTCTCTGAAA<br>ACTGCATCCGTGCTTACCGTGAACTGGCTGAAGCTCGTGGTGCTA<br>AAGTTCTGACCCACACCCGTGTTGAAGACTTCGACATCTCTCCGG<br>ACTCTGTTAAAATCGAAACCGCTAACGGTTCTTACACCGCTGACAA<br>ACTGATCGTTTCTATGGGTGCTTGGAACTCTAAACTGCTGTCTAAA |

| | |
|---|---|
| | CTGAACCTGGACATCCCGCTGCAGCCGTACCGTCAGGTTGTTGGT<br>TTCTTCGAATCTGACGAATCTAAATACTCTAACGACATCGACTTCC<br>CGGGTTTCATGGTTGAAGTTCCGAACGGTATCTACTACGGTTTCC<br>CGTCTTTCGGTGGTTGCGGTCTGAAACTGGGTTACCACACCTTCG<br>GTCAGAAAATCGACCCGGACACCATCAACCGTGAATTCGGTGTTT<br>ACCCGGAAGACGAATCTAACCTGCGTGCTTTCCTGGAAGAATACA<br>TGCCGGGTGCTAACGGTGAACTGAAACGTGGTGCTGTTTGCATGT<br>ACACCAAAACCCTGGACGAACACTTCATCATCGACCTGCACCCGG<br>AACACTCTAACGTTGTTATCGCTGCTGGTTTCAGCGGTCACGGTTT<br>CAAATTCTCTTCTGGTGTTGGTGAAGTTCTGTCTCAGCTGGCTCTG<br>ACCGGTAAAACCGAACACGACATCTCTATCTTCTCTATCAACCGTC<br>CGGCTCTGAAAGAAAGCCTGCAGAAAACCACCATCTAATAATGAac<br>tttatctgagaatagtcaatcttcggaaatcccaggtggcatgctaaaagtctcgtaaagcgttctatc<br>aataacccgttggtgccaggcatcaaataaaacgaaaggctcagtcgaaagactgggcctttcgt<br>tttatctgttgtttgtcggtgaacgctctctactagagtcacactggctcaccttcgggtgggcctttctgc<br>gtttataccgtctcagaatcggccgtgaacaataaaatagtttcggtattattgaccacttccgagtag<br>aatcgtgcttcagtaagagtcgaccgatgcccttgagagccttcaacccagtcagctccttccggtg<br>ggcgcggggcaTGACTATCGTCGCCGCACTTATGACTGTCTTCTTTATC<br>ATGCAACTCGTAGGACAGGTGCCGGCAGCGCTCTTCCGCTTCCTC<br>GCTCACTGACTCGCTGCGCTCGGTCGTTCGGCTGCGGCGAGCGG<br>TATCAGCTCACTCAAAGGCGGTAATACGGTTATCCACAGAATCAG<br>GGGATAACGCAGGAAAGAACATGTGAGCAAAAGGCCAGCAAAAG<br>GCCAGGAACCGTAAAAAGGCCGCGTTGCTGGCGTTTTTCCATAGG<br>CTCCGCCCCCCTGACGAGCATCACAAAAATCGACGCTCAAGTCAG<br>AGGTGGCGAAACCCGACAGGACTATAAAGATACCAGGCGTTTCCC<br>CCTGGAAGCTCCCTCGTGCGCTCTCCTGTTCCGACCCTGCCGCTT<br>ACCGGATACCTGTCCGCCTTTCTCCCTTCGGGAAGCGTGGCGCTT<br>TCTCAATGCTCACGCTGTAGGTATCTCAGTTCGGTGTAGGTCGTTC<br>GCTCCAAGCTGGGCTGTGTGCACGAACCCCCCGTTCAGCCCGAC<br>CGCTGCGCCTTATCCGGTAACTATCGTCTTGAGTCCAACCCGGTA<br>AGACACGACTTATCGCCACTGGCAGCAGCCACTGGTAACAGGATT<br>AGCAGAGCGAGGTATGTAGGCGGTGCTACAGAGTTCTTGAAGTG<br>GTGGCCTAACTACGGCTACACTAGAAGGACAGTATTTGGTATCTG<br>CGCTCTGCTGAAGCCAGTTACCTTCGGAAAAAGAGTTGGTAGCTC<br>TTGATCCGGCAAACAAACCACCGCTGGTAGCGGTGGTTTTTTTGTT<br>TGCAAGCAGCAGATTACGCGCAGAAAAAAAGGATCTCAAGAAGAT<br>CCTTTGATCTTTTCTACGGGGTCTGACGCTCAGTGGAACGAAAACT<br>CACGTTAAGGGATTTTGGTCATGAGATTATCAAAAAGGATCTTCAC<br>CTAGATCCTTTTAAATTAAAAATGAAGTTTTAAATCAATCTAAAGTAT<br>ATATGAGTAAACTTGGTCTGACAGTTACCAATGCTTAATCAGTGAG<br>GCACCTATCTCAGCGATCTGTCTATTTCGTTCATCCATAGTTGCCT<br>GACTCCCCGTCGTGTAGATAACTACGATACGGGAGGGCTTACCAT<br>CTGGCCCCAGTGCTGCAATGATACCGCGAGACCCACGCTCACCG<br>GCTCCAGATTTATCAGCAATAAACCAGCCAGCCGGAAGGGCCGAG<br>CGCAGAAGTGGTCCTGCAACTTTATCCGCCTCCATCCAGTCTATTA<br>ATTGTTGCCGGGAAGCTAGAGTAAGTAGTTCGCCAGTTAATAGTTT<br>GCGCAACGTTGTTGCCATTGCTACAGGCATCGTGGTGTCACGCTC<br>GTCGTTTGGTATGGCTTCATTCAGCTCCGGTTCCCAACGATCAAG<br>GCGAGTTACATGATCCCCCATGTTGTGCAAAAAAGCGGTTAGCTC<br>CTTCGGTCCTCCGATCGTTGTCAGAAGTAAGTTGGCCGCAGTGTT<br>ATCACTCATGGTTATGGCAGCACTGCATAATTCTCTTACTGTCATG<br>CCATCCGTAAGATGCTTTTCTGTGACTGGTGAGTACTCAACCAAGT<br>CATTCTGAGAATAGTGTATGCGGCGACCGAGTTGCTCTTGCCCGG<br>CGTCAATACGGGATAATACCGCGCCACATAGCAGAACTTTAAAAG |

| | |
|---|---|
| | TGCTCATCATTGGAAAACGTTCTTCGGGGCGAAAACTCTCAAGGA<br>TCTTACCGCTGTTGAGATCCAGTTCGATGTAACCCACTCGTGCAC<br>CCAACTGATCTTCAGCATCTTTTACTTTCACCAGCGTTTCTGGGTG<br>AGCAAAAACAGGAAGGCAAAATGCCGCAAAAAAGGGAATAAGGGC<br>GACACGGAAATGTTGAATACTCATACTCTTCCTTTTTCAATATTATT<br>GAAGCATTTATCAGGGTTATTGTCTCATGAGCGGATACATATTTGA<br>ATGTATTTAGAAAAATAAACAAATAGGGGTTCCGCGCACATTTCCC<br>CGAAAAGTGCCACCTGACGTCTAAGAAACCATTATTATCATGACAT<br>TAACCTATAAAAATAGGCGTATCACGAGGCCCTTTCGTCTTCAAGA<br>ATTCTGGCGAATCCTCTGACCAGCCAGAAAACGACCTTTCTGTGG<br>TGAAACCGGATGCTGCAATTCAGAGCGGCAGCAAGTGGGGGACA<br>GCAGAAGACCTGACCGCCGCAGAGTGGATGTTTGACATGGTGAA<br>GACTATCGCACCATCAGCCAGAAAACCGAATTTTGCTGGGTGGGC<br>TAACGATATCcgcctgatgcgtgaacgtgacggacgtaaccaccgcgacatgtgtgtgctg<br>ttccgctggctcggataccccttactctgttgaaaacgaatagataggttaaggaacggttatttctgcg<br>tagatctatcttacacagcatcacactggctcaccttcgggtgggcctttctgcgtttatatactagag<br>agagaatataaaaagccagattattaatccggctttttttattatttaggcaactgaaacgattcggatc<br>ctgtattactattctta |
| pBeast-pT7-soxA | taatacgactcactatagggagagctagcaataattttgtttaactttaagaaggagatataATGT<br>CAACACATTTTGATGTGATCGTTGTCGGGGCAGGTTCGATGGGAA<br>TGGCTGCTGGATATCAGTTGGCAAAACAAGGAGTAAAGACACTGT<br>TGGTAGACGCTTTTGACCCCCCGCACACAAATGGGTCTCATCATG<br>GCGACACACGTATTATTCGCCACGCATATGGGGAAGGACGTGAGT<br>ATGTACCATTGGCCTTGCGTTCACAGGAGTTATGGTACGAGTTGG<br>AGAAGGAGACCCACCACAAGATCTTCACAAAAACAGGCGTTTTAG<br>TTTTTGGACCGAAAGGGGAAAGCGCTTTCGTTGCCGAGACAATGG<br>AAGCCGCGAAAGAGCATTCTTTAACCGTAGACCTGCTGGAGGGG<br>GACGAGATCAATAAGCGCTGGCCCGGTATCACGGTCCCCGAGAA<br>CTATAACGCTATCTTTGAACCAAATTCTGGTGTCCTTTTTTCGGAG<br>AATTGCATTCGTGCTTACCGCGAGTTAGCAGAAGCCCGTGGTGCC<br>AAAGTATTGACACACACGCGCGTGGAAGACTTCGACATTTCACCC<br>GATTCTGTCAAAATTGAAACCGCTAATGGTTCCTATACGGCGGATA<br>AGCTTATCGTTAGCATGGGCGCATGGAACTCGAAGCTGCTGTCCA<br>AACTGAACTTGGATATCCCATTACAGCCGTATCGCCAAGTCGTCG<br>GCTTCTTTGAAAGCGATGAATCCAAATATAGCAACGATATTGACTT<br>TCCGGGGTTTATGGTCGAAGTACCAAATGGGATTTACTATGGCTTC<br>CCTTCCTTCGGTGGTTGTGGGCTTAAGCTGGGTTATCATACGTTTG<br>GACAGAAAATCGACCCGGACACGATTAACCGTGAATTCGGCGTAT<br>ACCCAGAAGATGAGTCAAACCTTCGCGCCTTTCTTGAAGAATATAT<br>GCCTGGTGCCAACGGGGAATTAAAACGTGGAGCCGTATGCATGTA<br>TACGAAGACATTAGACGAGCATTTCATCATTGACCTTCATCCCGAA<br>CATTCGAATGTGGTTATCGCCGCTGGGTTTAGCGGCCACGGCTTC<br>AAATTCAGCAGCGGGGTGGGAGAGGTACTGTCGCAATTAGCTTTA<br>ACTGGCAAAACGGAACATGATATTTCCATTTTCTCCATCAACCGCC<br>CAGCCTTGAAGGAATCCTTGCAGAAGACCACAATTactttatctgagaata<br>gtcaatcttcggaaatcccaggtggcatgctaaaagtctcgtaaagcgttctatcaataacccgttg<br>gtgCAAAGCCCGCCGAAAGGCGGGCTTTTCTGTccgtctcagaatcggccgt<br>gaacaataaaatagtttcggtattattgaccacttccgagtagaatcgtgcttcagtaagagtcgac<br>cgatgcccttgagagccttcaacccagtcagctccttccggtgggcgcggggcaTGACTATC<br>GTCGCCGCACTTATGACTGTCTTCTTTATCATGCAACTCGTAGGAC<br>AGGTGCCGGCAGCGCTCTTCCGCTTCCTCGCTCACTGACTCGCTG<br>CGCTCGGTCGTTCGGCTGCGGCGAGCGGTATCAGCTCACTCAAA<br>GGCGGTAATACGGTTATCCACAGAATCAGGGGATAACGCAGGAAA |

| | |
|---|---|
| | GAACATGTGAGCAAAAGGCCAGCAAAAGGCCAGGAACCGTAAAAA<br>GGCCGCGTTGCTGGCGTTTTTCCATAGGCTCCGCCCCCCTGACG<br>AGCATCACAAAAATCGACGCTCAAGTCAGAGGTGGCGAAACCCGA<br>CAGGACTATAAAGATACCAGGCGTTTCCCCCTGGAAGCTCCCTCG<br>TGCGCTCTCCTGTTCCGACCCTGCCGCTTACCGGATACCTGTCCG<br>CCTTTCTCCCTTCGGGAAGCGTGGCGCTTTCTCAATGCTCACGCT<br>GTAGGTATCTCAGTTCGGTGTAGGTCGTTCGCTCCAAGCTGGGCT<br>GTGTGCACGAACCCCCCGTTCAGCCCGACCGCTGCGCCTTATCC<br>GGTAACTATCGTCTTGAGTCCAACCCGGTAAGACACGACTTATCG<br>CCACTGGCAGCAGCCACTGGTAACAGGATTAGCAGAGCGAGGTA<br>TGTAGGCGGTGCTACAGAGTTCTTGAAGTGGTGGCCTAACTACGG<br>CTACACTAGAAGGACAGTATTTGGTATCTGCGCTCTGCTGAAGCC<br>AGTTACCTTCGGAAAAAGAGTTGGTAGCTCTTGATCCGGCAAACA<br>AACCACCGCTGGTAGCGGTGGTTTTTTTGTTTGCAAGCAGCAGAT<br>TACGCGCAGAAAAAAAGGATCTCAAGAAGATCCTTTGATCTTTTCT<br>ACGGGGTCTGACGCTCAGTGGAACGAAAACTCACGTTAAGGGATT<br>TTGGTCATGAGATTATCAAAAAGGATCTTCACCTAGATCCTTTTAAA<br>TTAAAAATGAAGTTTTAAATCAATCTAAAGTATATATGAGTAAACTT<br>GGTCTGACAGTTACCAATGCTTAATCAGTGAGGCACCTATCTCAG<br>CGATCTGTCTATTTCGTTCATCCATAGTTGCCTGACTCCCCGTCGT<br>GTAGATAACTACGATACGGGAGGGCTTACCATCTGGCCCCAGTGC<br>TGCAATGATACCGCGAGACCCACGCTCACCGGCTCCAGATTTATC<br>AGCAATAAACCAGCCAGCCGGAAGGGCCGAGCGCAGAAGTGGTC<br>CTGCAACTTTATCCGCCTCCATCCAGTCTATTAATTGTTGCCGGGA<br>AGCTAGAGTAAGTAGTTCGCCAGTTAATAGTTTGCGCAACGTTGTT<br>GCCATTGCTACAGGCATCGTGGTGTCACGCTCGTCGTTTGGTATG<br>GCTTCATTCAGCTCCGGTTCCCAACGATCAAGGCGAGTTACATGA<br>TCCCCCATGTTGTGCAAAAAAGCGGTTAGCTCCTTCGGTCCTCCG<br>ATCGTTGTCAGAAGTAAGTTGGCCGCAGTGTTATCACTCATGGTTA<br>TGGCAGCACTGCATAATTCTCTTACTGTCATGCCATCCGTAAGATG<br>CTTTTCTGTGACTGGTGAGTACTCAACCAAGTCATTCTGAGAATAG<br>TGTATGCGGCGACCGAGTTGCTCTTGCCCGGCGTCAATACGGGAT<br>AATACCGCGCCACATAGCAGAACTTTAAAAGTGCTCATCATTGGAA<br>AACGTTCTTCGGGGCGAAAACTCTCAAGGATCTTACCGCTGTTGA<br>GATCCAGTTCGATGTAACCCACTCGTGCACCCAACTGATCTTCAG<br>CATCTTTTACTTTCACCAGCGTTTCTGGGTGAGCAAAAACAGGAAG<br>GCAAAATGCCGCAAAAAAGGGAATAAGGGCGACACGGAAATGTTG<br>AATACTCATACTCTTCCTTTTTCAATATTATTGAAGCATTTATCAGG<br>GTTATTGTCTCATGAGCGGATACATATTTGAATGTATTTAGAAAAAT<br>AAACAAATAGGGGTTCCGCGCACATTTCCCCGAAAAGTGCCACCT<br>GACGTCTAAGAAACCATTATTATCATGACATTAACCTATAAAAATAG<br>GCGTATCACGAGGCCCTTTCGTCTTCAAGAATTCTGGCGAATCCT<br>CTGACCAGCCAGAAAACGACCTTTCTGTGGTGAAACCGGATGCTG<br>CAATTCAGAGCGGCAGCAAGTGGGGGACAGCAGAAGACCTGACC<br>GCCGCAGAGTGGATGTTTGACATGGTGAAGACTATCGCACCATCA<br>GCCAGAAAACCGAATTTTGCTGGGTGGGCTAACGATATCcgcctgatg<br>cgtgaacgtgacggacgtaaccaccgcgacatgtgtgctgttccgctggctcggataccccttact<br>ctgttgaaaacgaatagataggttaaggaacggttatttctgcgtagatctatcttacacagcatcac<br>actggctcaccttcgggtgggcctttctgcgtttatatactagagagagaatataaaaagccagatt<br>attaatccggctttttttattatttaggcaactgaaacgattcggatcctgtattactattctta |
| pBeast-pT7-codA | taatacgactcactatagggagagctagcaataattttgtttaactttaagaaggagatataATGC<br>ACATCGACAACATCGAAAACCTGTCTGACCGTGAATTCGACTACAT<br>CGTTGTTGGTGGTGGTTCTGCTGGTGCTGCTGTTGCTGCTCGTCT |

| | |
|---|---|
| | GTCTGAAGACCCGGCTGTTTCTGTTGCTCTGGTTGAAGCTGGTCC<br>GGACGACCGTGGTGTTCCGGAAGTTCTGCAGCTGGACCGTTGGA<br>TGGAACTGCTGGAATCTGGTTACGACTGGGACTACCCGATCGAAC<br>CGCAGGAAAACGGTAACTCTTTCATGCGTCACGCTCGTGCTAAAG<br>TTATGGGTGGTTGCTCTTCTCACAACTCTTGCATCGCTTTCTGGGC<br>TCCGCGTGAAGACCTGGACGAATGGGAAGCTAAATACGGTGCTAC<br>CGGTTGGAACGCTGAAGCTGCTTGGCCGCTGTACAAACGTCTGGA<br>AACCAACGAAGACGCTGGTCCGGACGCTCCGCACCACGGTGACT<br>CTGGTCCGGTTCACCTGATGAACGTTCCGCCGAAAGACCCGACC<br>GGTGTTGCTCTGCTGGACGCTTGCGAACAGGCTGGTATCCCGCG<br>TGCTAAATTCAACACCGGTACCACCGTTGTTAACGGTGCTAACTTC<br>TTCCAGATCAACCGTCGTGCTGACGGTACCCGTTCTTCTTCTTCTG<br>TTTCTTACATCCACCCGATCGTTGAACAGGAAAACTTCACCCTGCT<br>GACCGGTCTGCGTGCTCGTCAGCTGGTTTTCGACGCTGACCGTC<br>GTTGCACCGGTGTTGACATCGTTGACTCTGCTTTCGGTCACACCC<br>ACCGTCTGACCGCTCGTAACGAAGTTGTTCTGTCTACCGGTGCTA<br>TCGACACCCCGAAACTGCTGATGCTGTCTGGTATCGGTCCGGCTG<br>CTCACCTGGCTGAACACGGTATCGAAGTTCTGGTTGACTCTCCGG<br>GTGTTGGTGAACACCTGCAGGACCACCCGGAAGGTGTTGTTCAGT<br>TCGAAGCTAAACAGCCGATGGTTGCTGAATCTACCCAGTGGTGG<br>AAATCGGTATCTTCACCCCGACCGAAGACGGTCTGGACCGTCCGG<br>ACCTGATGATGCACTACGGTTCTGTTCCGTTCGACATGAACACCCT<br>GCGTCACGGTTACCCGACCACCGAAAACGGTTTCTCTCTGACCCC<br>GAACGTTACCCACGCTCGTTCTCGTGGTACCGTTCGTCTGCGTTC<br>TCGTGACTTCCGTGACAAACCGATGGTTGACCCGCGTTACTTCAC<br>CGACCCGGAAGGTCACGACATGCGTGTTATGGTTGCTGGTATCCG<br>TAAAGCTCGTGAAATCGCTGCTCAGCCGGCTATGGCTGAATGGAC<br>CGGTCGTGAACTGTCTCCGGGTGTTGAAGCTCAGACCGACGAAG<br>AACTGCAGGACTACATCCGTAAAACCCACAACACCGTTTACCACC<br>CGGTTGGTACCGTTCGTATGGGTGCTGTTGAAGACGAAATGTCTC<br>CGCTGGACCCGGAACTGCGTGTTAAAGGTGTTACCGGTCTGCGT<br>GTTGCTGACGCTTCTGTTATGCCGGAACACGTTACCGTTAACCCG<br>AACATCACCGTTATGATGATCGGTGAACGTTGCGCTGACCTGATC<br>CGTTCTGCTCGTGCTGGTGAAACCACCACCGCTGACGCTGAACTG<br>TCTGCTGCTCTGGCTactttatctgagaatagtcaatcttcggaaatcccaggtggcatg<br>ctaaaagtctcgtaaagcgttctatcaataacccgttggtgCAAAGCCCGCCGAAAGG<br>CGGGCTTTTCTGTccgtctcagaatcggccgtgaacaataaaatagtttcggtattattgac<br>cacttccgagtagaatcgtgcttcagtaagagtcgaccgatgcccttgagagccttcaacccagtc<br>agctccttccggtgggcgcggggcaTGACTATCGTCGCCGCACTTATGACTGT<br>CTTCTTTATCATGCAACTCGTAGGACAGGTGCCGGCAGCGCTCTT<br>CCGCTTCCTCGCTCACTGACTCGCTGCGCTCGGTCGTTCGGCTGC<br>GGCGAGCGGTATCAGCTCACTCAAAGGCGGTAATACGGTTATCCA<br>CAGAATCAGGGGATAACGCAGGAAAGAACATGTGAGCAAAAGGC<br>CAGCAAAAGGCCAGGAACCGTAAAAAGGCCGCGTTGCTGGCGTT<br>TTTCCATAGGCTCCGCCCCCCTGACGAGCATCACAAAAATCGACG<br>CTCAAGTCAGAGGTGGCGAAACCCGACAGGACTATAAAGATACCA<br>GGCGTTTCCCCCTGGAAGCTCCCTCGTGCGCTCTCCTGTTCCGAC<br>CCTGCCGCTTACCGGATACCTGTCCGCCTTTCTCCCTTCGGGAAG<br>CGTGGCGCTTTCTCAATGCTCACGCTGTAGGTATCTCAGTTCGGT<br>GTAGGTCGTTCGCTCCAAGCTGGGCTGTGTGCACGAACCCCCCG<br>TTCAGCCCGACCGCTGCGCCTTATCCGGTAACTATCGTCTTGAGT<br>CCAACCCGGTAAGACACGACTTATCGCCACTGGCAGCAGCCACTG<br>GTAACAGGATTAGCAGAGCGAGGTATGTAGGCGGTGCTACAGAGT<br>TCTTGAAGTGGTGGCCTAACTACGGCTACACTAGAAGGACAGTAT |

| | |
|---|---|
| | TTGGTATCTGCGCTCTGCTGAAGCCAGTTACCTTCGGAAAAAGAG<br>TTGGTAGCTCTTGATCCGGCAAACAAACCACCGCTGGTAGCGGTG<br>GTTTTTTTGTTTGCAAGCAGCAGATTACGCGCAGAAAAAAAGGATC<br>TCAAGAAGATCCTTTGATCTTTTCTACGGGGTCTGACGCTCAGTGG<br>AACGAAAACTCACGTTAAGGGATTTTGGTCATGAGATTATCAAAAA<br>GGATCTTCACCTAGATCCTTTTAAATTAAAAATGAAGTTTTAAATCA<br>ATCTAAAGTATATATGAGTAAACTTGGTCTGACAGTTACCAATGCTT<br>AATCAGTGAGGCACCTATCTCAGCGATCTGTCTATTTCGTTCATCC<br>ATAGTTGCCTGACTCCCCGTCGTGTAGATAACTACGATACGGGAG<br>GGCTTACCATCTGGCCCCAGTGCTGCAATGATACCGCGAGACCCA<br>CGCTCACCGGCTCCAGATTTATCAGCAATAAACCAGCCAGCCGGA<br>AGGGCCGAGCGCAGAAGTGGTCCTGCAACTTTATCCGCCTCCATC<br>CAGTCTATTAATTGTTGCCGGGAAGCTAGAGTAAGTAGTTCGCCA<br>GTTAATAGTTTGCGCAACGTTGTTGCCATTGCTACAGGCATCGTG<br>GTGTCACGCTCGTCGTTTGGTATGGCTTCATTCAGCTCCGGTTCC<br>CAACGATCAAGGCGAGTTACATGATCCCCCATGTTGTGCAAAAAA<br>GCGGTTAGCTCCTTCGGTCCTCCGATCGTTGTCAGAAGTAAGTTG<br>GCCGCAGTGTTATCACTCATGGTTATGGCAGCACTGCATAATTCTC<br>TTACTGTCATGCCATCCGTAAGATGCTTTTCTGTGACTGGTGAGTA<br>CTCAACCAAGTCATTCTGAGAATAGTGTATGCGGCGACCGAGTTG<br>CTCTTGCCCGGCGTCAATACGGGATAATACCGCGCCACATAGCAG<br>AACTTTAAAAGTGCTCATCATTGGAAAACGTTCTTCGGGGCGAAAA<br>CTCTCAAGGATCTTACCGCTGTTGAGATCCAGTTCGATGTAACCCA<br>CTCGTGCACCCAACTGATCTTCAGCATCTTTTACTTTCACCAGCGT<br>TTCTGGGTGAGCAAAAACAGGAAGGCAAAATGCCGCAAAAAAGGG<br>AATAAGGGCGACACGGAAATGTTGAATACTCATACTCTTCCTTTTT<br>CAATATTATTGAAGCATTTATCAGGGTTATTGTCTCATGAGCGGAT<br>ACATATTTGAATGTATTTAGAAAAATAAACAAATAGGGGTTCCGCG<br>CACATTTCCCCGAAAAGTGCCACCTGACGTCTAAGAAACCATTATT<br>ATCATGACATTAACCTATAAAAATAGGCGTATCACGAGGCCCTTTC<br>GTCTTCAAGAATTCTGGCGAATCCTCTGACCAGCCAGAAAACGAC<br>CTTTCTGTGGTGAAACCGGATGCTGCAATTCAGAGCGGCAGCAAG<br>TGGGGGACAGCAGAAGACCTGACCGCCGCAGAGTGGATGTTTGA<br>CATGGTGAAGACTATCGCACCATCAGCCAGAAAACCGAATTTTGC<br>TGGGTGGGCTAACGATATCcgcctgatgcgtgaacgtgacggacgtaaccaccgc<br>gacatgtgtgtgctgttccgctggctcggatacccttactctgttgaaaacgaatagataggttaagg<br>aacggttatttctgcgtagatctatcttacacagcatcacactggctcaccttcgggtgggcctttctgc<br>gtttatatactagagagagaatataaaaagccagattattaatccggctttttttattatttaggcaactg<br>aaacgattcggatcctgtattactattctta |
| pBeast-pT7-lox | taatacgactcactatagggagagctagcaataattttgtttaactttaagaaggagatataATGA<br>ATAACAATGACATTGAATATAATGCACCTAGTGAAATCAAGTACATT<br>GATGTTGTCAATACTTACGACTTAGAAGAAGAAGCAAGTAAAGTGG<br>TACCACATGGTGGTTTTAACTATATTGCCGGTGCATCTGGTGATGA<br>GTGGACTAAACGCGCTAATGACCGTGCTTGGAAACATAAATTACTA<br>TACCCACGTCTAGCGCAAGATGTTGAAGCGCCCGATACAAGTACT<br>GAAATTTTAGGTCATAAAATTAAAGCCCCATTCATCATGGCACCAA<br>TTGCTGCACATGGTTTAGCCCACACTACTAAAGAAGCTGGTACTG<br>CACGTGCAGTTTCAGAATTTGGTACAATTATGTCCATCTCAGCTTA<br>TTCTGGTGCAACATTTGAAGAAATTTCTGAAGGCTTAAATGGCGGA<br>CCCCGTTGGTTCCAAATCTATATGGCTAAAGATGACCAACAAAACC<br>GTGATATCTTAGACGAAGCTAAATCTGATGGTGCAACTGCTATCAT<br>CCTTACAGCTGACTCAACTGTTTCTGGAAACCGTGACCGTGATGT<br>GAAGAATAAATTCGTTTACCCATTTGGTATGCCAATTGTTCAACGTT |

| | |
|---|---|
| | ACTTACGTGGTACAGCAGAAGGTATGTCATTAAACAATATCTACGG<br>TGCTTCAAAACAAAAAATCTCACCAAGAGATATTGAGGAAATCGCC<br>GCTCATTCTGGATTACCAGTATTCGTTAAAGGTATTCAACACCCAG<br>AAGATGCAGATATGGCAATCAAAGCTGGTGCATCAGGTATCTGGG<br>TATCTAACCACGGTGCTCGTCAACTATATGAAGCTCCAGGTTCATT<br>TGACACCCTTCCAGCTATTGCTGAACGTGTAAACAAACGTGTACCA<br>ATCGTCTTTGATTCAGGTGTACGTCGTGGTGAACACGTTGCCAAA<br>GCGCTAGCTTCAGGGGCAGACGTTGTTGCTTTAGGACGCCCAGTC<br>TTATTTGGTTTAGCTTTAGGTGGCTGGCAAGGTGCTTACTCAGTAC<br>TTGACTACTTCCAAAAAGACTTAACACGCGTAATGCAATTAACAGG<br>TTCACAAAATGTGGAAGACTTGAAGGGTCTAGATTTATTCGATAAC<br>CCATACGGTTATGAATACTAGagcgttctatcaataacccgttggtgCAAAGCC<br>CGCCGAAAGGCGGGCTTTTCTGTccgtctcagaatcggccgtgaacaataaaat<br>agtttcggtattattgaccacttccgagtagaatcgtgcttcagtaagagtcgaccgatgcccttgag<br>agccttcaacccagtcagctccttccggtgggcgcggggcaTGACTATCGTCGCCGC<br>ACTTATGACTGTCTTCTTTATCATGCAACTCGTAGGACAGGTGCCG<br>GCAGCGCTCTTCCGCTTCCTCGCTCACTGACTCGCTGCGCTCGGT<br>CGTTCGGCTGCGGCGAGCGGTATCAGCTCACTCAAAGGCGGTAA<br>TACGGTTATCCACAGAATCAGGGGATAACGCAGGAAAGAACATGT<br>GAGCAAAAGGCCAGCAAAAGGCCAGGAACCGTAAAAAGGCCGCG<br>TTGCTGGCGTTTTTCCATAGGCTCCGCCCCCCTGACGAGCATCAC<br>AAAAATCGACGCTCAAGTCAGAGGTGGCGAAACCCGACAGGACTA<br>TAAAGATACCAGGCGTTTCCCCCTGGAAGCTCCCTCGTGCGCTCT<br>CCTGTTCCGACCCTGCCGCTTACCGGATACCTGTCCGCCTTTCTC<br>CCTTCGGGAAGCGTGGCGCTTTCTCAATGCTCACGCTGTAGGTAT<br>CTCAGTTCGGTGTAGGTCGTTCGCTCCAAGCTGGGCTGTGTGCAC<br>GAACCCCCCGTTCAGCCCGACCGCTGCGCCTTATCCGGTAACTAT<br>CGTCTTGAGTCCAACCCGGTAAGACACGACTTATCGCCACTGGCA<br>GCAGCCACTGGTAACAGGATTAGCAGAGCGAGGTATGTAGGCGG<br>TGCTACAGAGTTCTTGAAGTGGTGGCCTAACTACGGCTACACTAG<br>AAGGACAGTATTTGGTATCTGCGCTCTGCTGAAGCCAGTTACCTTC<br>GGAAAAAGAGTTGGTAGCTCTTGATCCGGCAAACAAACCACCGCT<br>GGTAGCGGTGGTTTTTTTGTTTGCAAGCAGCAGATTACGCGCAGA<br>AAAAAAGGATCTCAAGAAGATCCTTTGATCTTTTCTACGGGGTCTG<br>ACGCTCAGTGGAACGAAAACTCACGTTAAGGGATTTTGGTCATGA<br>GATTATCAAAAAGGATCTTCACCTAGATCCTTTTAAATTAAAAATGA<br>AGTTTTAAATCAATCTAAAGTATATATGAGTAAACTTGGTCTGACAG<br>TTACCAATGCTTAATCAGTGAGGCACCTATCTCAGCGATCTGTCTA<br>TTTCGTTCATCCATAGTTGCCTGACTCCCCGTCGTGTAGATAACTA<br>CGATACGGGAGGGCTTACCATCTGGCCCCAGTGCTGCAATGATAC<br>CGCGAGACCCACGCTCACCGGCTCCAGATTTATCAGCAATAAACC<br>AGCCAGCCGGAAGGGCCGAGCGCAGAAGTGGTCCTGCAACTTTA<br>TCCGCCTCCATCCAGTCTATTAATTGTTGCCGGGAAGCTAGAGTAA<br>GTAGTTCGCCAGTTAATAGTTTGCGCAACGTTGTTGCCATTGCTAC<br>AGGCATCGTGGTGTCACGCTCGTCGTTTGGTATGGCTTCATTCAG<br>CTCCGGTTCCCAACGATCAAGGCGAGTTACATGATCCCCCATGTT<br>GTGCAAAAAAGCGGTTAGCTCCTTCGGTCCTCCGATCGTTGTCAG<br>AAGTAAGTTGGCCGCAGTGTTATCACTCATGGTTATGGCAGCACT<br>GCATAATTCTCTTACTGTCATGCCATCCGTAAGATGCTTTTCTGTG<br>ACTGGTGAGTACTCAACCAAGTCATTCTGAGAATAGTGTATGCGG<br>CGACCGAGTTGCTCTTGCCCGGCGTCAATACGGGATAATACCGCG<br>CCACATAGCAGAACTTTAAAAGTGCTCATCATTGGAAAACGTTCTT<br>CGGGGCGAAAACTCTCAAGGATCTTACCGCTGTTGAGATCCAGTT<br>CGATGTAACCCACTCGTGCACCCAACTGATCTTCAGCATCTTTTAC |

| | |
|---|---|
| | TTTCACCAGCGTTTCTGGGTGAGCAAAAACAGGAAGGCAAAATGC<br>CGCAAAAAAGGGAATAAGGGCGACACGGAAATGTTGAATACTCAT<br>ACTCTTCCTTTTTCAATATTATTGAAGCATTTATCAGGGTTATTGTC<br>TCATGAGCGGATACATATTTGAATGTATTTAGAAAAATAAACAAATA<br>GGGGTTCCGCGCACATTTCCCCGAAAAGTGCCACCTGACGTCTAA<br>GAAACCATTATTATCATGACATTAACCTATAAAAATAGGCGTATCAC<br>GAGGCCCTTTCGTCTTCAAGAATTCTGGCGAATCCTCTGACCAGC<br>CAGAAAACGACCTTTCTGTGGTGAAACCGGATGCTGCAATTCAGA<br>GCGGCAGCAAGTGGGGGACAGCAGAAGACCTGACCGCCGCAGA<br>GTGGATGTTTGACATGGTGAAGACTATCGCACCATCAGCCAGAAA<br>ACCGAATTTTGCTGGGTGGGCTAACGATATCcgcctgatgcgtgaacgtgac<br>ggacgtaaccaccgcgacatgtgtgtgctgttccgctggctcggatacccttactctgttgaaaacg<br>aatagataggttaaggaacggttatttctgcgtagatctatcttacacagcatcacactggctcacctt<br>cgggtgggcctttctgcgtttatatactagagagagaatataaaaagccagattattaatccggctttt<br>ttattatttaggcaactgaaacgattcggatcctgtattactattctta |
| pBeast-pAhpC-RBS-LacZ | GCTTAGATCAGGTGATTGCCCTTTGTTTATGAGGGTGTTGTAATCC<br>ATGTCGTTGTTGCATTTGTAAGGGCAACACCTCAGCCTGCAGGCA<br>GGCACTGAAGATACCAAAGGGTAGTTCAGATTACACGGTCACCTG<br>GAAAGGGGGCCATTTTACTTTTTATCGCCGCTGGCGGTGCAAAGT<br>TCACAAAGTTGTCTTACGAAGGTTGTAAGGTAAAACTTATCGATTT<br>GATAATGGAAACGCATTAGCCGAATCGGCAAAAATTGGTTACCTTA<br>CATCTCATCGAAAACACGGAGGAAGTATAGATGTCTAGAGAAAGA<br>GGAGAAATACTAGATGaccatgattacggattcactggccgtcgttttacaacgtcgtga<br>ctgggaaaaccctggcgttacccaacttaatcgccttgcagcacatccccctttcgccagctggcgt<br>aatagcgaagaggcccgcaccgatcgcccttcccaacagttgcgcagcctgaatggcgaatgg<br>cgctttgcctggtttccggcaccagaagcggtgccggaaagctggctggagtgcgatcttcctgag<br>gccgatactgtcgtcgtcccctcaaactggcagatgcacggttacgatgcgcccatctacaccaac<br>gtgacctatcccattacggtcaatccgccgtttgttcccacggagaatccgacgggttgttactcgct<br>cacatttaatgttgatgaaagctggctacaggaaggccagacgcgaattatttttgatggcgttaact<br>cggcgtttcatctgtggtgcaacgggcgctgggtcggttacggccaggacagtcgtttgccgtctga<br>atttgacctgagcgcatttttacgcgccggagaaaaccgcctcgcggtgatggtgctgcgctggag<br>tgacggcagttatctggaagatcaggatatgtggcggatgagcggcattttccgtgacgtctcgttg<br>ctgcataaaccgactacacaaatcagcgatttccatgttgccactcgctttaatgatgatttcagccg<br>cgctgtactggaggctgaagttcagatgtgcggcgagttgcgtgactacctacgggtaacagtttctt<br>tatggcagggtgaaacgcaggtcgccagcggcaccgcgcctttcggcggtgaaattatcgatga<br>gcgtggtggttatgccgatcgcgtcacactacgtctgaacgtcgaaaacccgaaactgtggagcg<br>ccgaaatcccgaatctctatcgtgcggtggttgaactgcacaccgccgacggcacgctgattgaa<br>gcagaagcctgcgatgtcggtttccgcgaggtgcggattgaaaatggtctgctgctgctgaacggc<br>aagccgttgctgattcgaggcgttaaccgtcacgagcatcatcctctgcatggtcaggtcatggatg<br>agcagacgatggtgcaggatatcctgctgatgaagcagaacaactttaacgccgtgcgctgttcg<br>cattatccgaaccatccgctgtggtacacgctgtgcgaccgctacggcctgtatgtggtggatgaa<br>gccaatattgaaacccacggcatggtgccaatgaatcgtctgaccgatgatccgcgctggctacc<br>ggcgatgagcgaacgcgtaacgcgaatggtgcagcgcgatcgtaatcacccgagtgtgatcatc<br>tggtcgctggggaatgaatcaggccacggcgctaatcacgacgcgctgtatcgctggatcaaatc<br>tgtcgatccttcccgcccggtgcagtatgaaggcggcggagccgacaccacggccaccgatatt<br>atttgcccgatgtacgcgcgcgtggatgaagaccagcccttcccggctgtgccgaaatggtccatc<br>aaaaaatggctttcgctacctggagagacgcgcccgctgatcctttgcgaatacgcccacgcgat<br>gggtaacagtcttggcggtttcgctaaatactggcaggcgtttcgtcagtatccccgtttacagggcg<br>gcttcgtctgggactgggtggatcagtcgctgattaaatatgatgaaaacggcaacccgtggtcgg<br>cttacggcggtgattttggcgatacgccgaacgatcgccagttctgtatgaacggtctggtctttgcc<br>gaccgcacgccgcatccagcgctgacggaagcaaaacaccagcagcagtttttccagttccgttt<br>atccgggcaaaccatcgaagtgaccagcgaatacctgttccgtcatagcgataacgagctcctgc<br>actggatggtggcgctggatggtaagccgctggcaagcggtgaagtgcctctggatgtcgctcca |

| | |
|---|---|
| | caaggtaaacagttgattgaactgcctgaactaccgcagccggagagcgccgggcaactctgg<br>ctcacagtacgcgtagtgcaaccgaacgcgaccgcatggtcagaagccgggcacatcagcgc<br>ctggcagcagtggcgtctggcggaaaacctcagtgtgacgctccccgccgcgtcccacgccatc<br>ccgcatctgaccaccagcgaaatggatttttgcatcgagctgggtaataagcgttggcaatttaacc<br>gccagtcaggctttctttcacagatgtggattggcgataaaaaacaactgctgacgccgctgcgcg<br>atcagttcacccgtgcaccgctggataacgacattggcgtaagtgaagcgacccgcattgaccct<br>aacgcctgggtcgaacgctggaaggcggcgggccattaccaggccgaagcagcgttgttgcag<br>tgcacggcagatacacttgctgatgcggtgctgattacgaccgctcacgcgtggcagcatcaggg<br>gaaaaccttatttatcagccggaaaacctaccggattgatggtagtggtcaaatggcgattaccgtt<br>gatgttgaagtggcgagcgatacaccgcatccggcgcggattggcctgaactgccagctggcgc<br>aggtagcagagcgggtaaactggctcggattagggccgcaagaaaactatcccgaccgcctta<br>ctgccgcctgttttgaccgctgggatctgccattgtcagacatgtataccccgtacgtcttcccgagc<br>gaaaacggtctgcgctgcgggacgcgcgaattgaattatggcccacaccagtggcgcggcgac<br>ttccagttcaacatcagccgctacagtcaacagcaactgatggaaaccagccatcgccatctgct<br>gcacgcggaagaaggcacatggctgaatatcgacggtttccatatggggattggtggcgacgac<br>tcctggagcccgtcagtatcggcggaattccagctgagcgccggtcgctaccattaccagttggtct<br>ggtgtcaaaaataaactttatctgagaatagtcaatcttcggaaatcccaggtggcatgctaaaagt<br>ctcgtaaagcgttctatcaataacccgttggtgccaggcatcaaataaaacgaaaggctcagtcg<br>aaagactgggcctttcgttttatctgttgtttgtcggtgaacgctctctactagagtcacactggctcac<br>cttcgggtgggcctttctgcgtttataccgtctcagaatcggccgtgaacaataaaatagtttcggtatt<br>attgaccacttccgagtagaatcgtgcttcagtaagagtcgaccgatgcccttgagagccttcaacc<br>cagtcagctccttccggtgggcgcggggcaTGACTATCGTCGCCGCACTTATGA<br>CTGTCTTCTTTATCATGCAACTCGTAGGACAGGTGCCGGCAGCGC<br>TCTTCCGCTTCCTCGCTCACTGACTCGCTGCGCTCGGTCGTTCGG<br>CTGCGGCGAGCGGTATCAGCTCACTCAAAGGCGGTAATACGGTTA<br>TCCACAGAATCAGGGGATAACGCAGGAAAGAACATGTGAGCAAAA<br>GGCCAGCAAAAGGCCAGGAACCGTAAAAAGGCCGCGTTGCTGGC<br>GTTTTTCCATAGGCTCCGCCCCCCTGACGAGCATCACAAAAATCG<br>ACGCTCAAGTCAGAGGTGGCGAAACCCGACAGGACTATAAAGATA<br>CCAGGCGTTTCCCCCTGGAAGCTCCCTCGTGCGCTCTCCTGTTCC<br>GACCCTGCCGCTTACCGGATACCTGTCCGCCTTTCTCCCTTCGGG<br>AAGCGTGGCGCTTTCTCAATGCTCACGCTGTAGGTATCTCAGTTC<br>GGTGTAGGTCGTTCGCTCCAAGCTGGGCTGTGTGCACGAACCCC<br>CCGTTCAGCCCGACCGCTGCGCCTTATCCGGTAACTATCGTCTTG<br>AGTCCAACCCGGTAAGACACGACTTATCGCCACTGGCAGCAGCCA<br>CTGGTAACAGGATTAGCAGAGCGAGGTATGTAGGCGGTGCTACA<br>GAGTTCTTGAAGTGGTGGCCTAACTACGGCTACACTAGAAGGACA<br>GTATTTGGTATCTGCGCTCTGCTGAAGCCAGTTACCTTCGGAAAAA<br>GAGTTGGTAGCTCTTGATCCGGCAAACAAACCACCGCTGGTAGCG<br>GTGGTTTTTTTGTTTGCAAGCAGCAGATTACGCGCAGAAAAAAAGG<br>ATCTCAAGAAGATCCTTTGATCTTTTCTACGGGGTCTGACGCTCAG<br>TGGAACGAAAACTCACGTTAAGGGATTTTGGTCATGAGATTATCAA<br>AAAGGATCTTCACCTAGATCCTTTTAAATTAAAAATGAAGTTTTAAA<br>TCAATCTAAAGTATATATGAGTAAACTTGGTCTGACAGTTACCAAT<br>GCTTAATCAGTGAGGCACCTATCTCAGCGATCTGTCTATTTCGTTC<br>ATCCATAGTTGCCTGACTCCCCGTCGTGTAGATAACTACGATACG<br>GGAGGGCTTACCATCTGGCCCCAGTGCTGCAATGATACCGCGAG<br>ACCCACGCTCACCGGCTCCAGATTTATCAGCAATAAACCAGCCAG<br>CCGGAAGGGCCGAGCGCAGAAGTGGTCCTGCAACTTTATCCGCC<br>TCCATCCAGTCTATTAATTGTTGCCGGGAAGCTAGAGTAAGTAGTT<br>CGCCAGTTAATAGTTTGCGCAACGTTGTTGCCATTGCTACAGGCAT<br>CGTGGTGTCACGCTCGTCGTTTGGTATGGCTTCATTCAGCTCCGG<br>TTCCCAACGATCAAGGCGAGTTACATGATCCCCCATGTTGTGCAA<br>AAAAGCGGTTAGCTCCTTCGGTCCTCCGATCGTTGTCAGAAGTAA |

| | |
|---|---|
| | GTTGGCCGCAGTGTTATCACTCATGGTTATGGCAGCACTGCATAA<br>TTCTCTTACTGTCATGCCATCCGTAAGATGCTTTTCTGTGACTGGT<br>GAGTACTCAACCAAGTCATTCTGAGAATAGTGTATGCGGCGACCG<br>AGTTGCTCTTGCCCGGCGTCAATACGGGATAATACCGCGCCACAT<br>AGCAGAACTTTAAAAGTGCTCATCATTGGAAAACGTTCTTCGGGGC<br>GAAAACTCTCAAGGATCTTACCGCTGTTGAGATCCAGTTCGATGTA<br>ACCCACTCGTGCACCCAACTGATCTTCAGCATCTTTTACTTTCACC<br>AGCGTTTCTGGGTGAGCAAAAACAGGAAGGCAAAATGCCGCAAAA<br>AAGGGAATAAGGGCGACACGGAAATGTTGAATACTCATACTCTTC<br>CTTTTTCAATATTATTGAAGCATTTATCAGGGTTATTGTCTCATGAG<br>CGGATACATATTTGAATGTATTTAGAAAAATAAACAAATAGGGGTT<br>CCGCGCACATTTCCCCGAAAAGTGCCACCTGACGTCTAAGAAACC<br>ATTATTATCATGACATTAACCTATAAAAATAGGCGTATCACGAGGC<br>CCTTTCGTCTTCAAGAATTCTGGCGAATCCTCTGACCAGCCAGAAA<br>ACGACCTTTCTGTGGTGAAACCGGATGCTGCAATTCAGAGCGGCA<br>GCAAGTGGGGGACAGCAGAAGACCTGACCGCCGCAGAGTGGATG<br>TTTGACATGGTGAAGACTATCGCACCATCAGCCAGAAAACCGAATT<br>TTGCTGGGTGGGCTAACGATATCcgcctgatgcgtgaacgtgacggacgtaacc<br>accgcgacatgtgtgtgctgttccgctggctcggatacccttactctgttgaaaacgaatagataggt<br>taaggaacggttatttctgcgtagatctatcttacacagcatcacactggctcaccttcgggtgggcc<br>tttctgcgtttatatactagagagagaatataaaaagccagattattaatccggctttttattatttaggc<br>aactgaaacgattcggatcctgtattactattctta |
| pBeast-pAhpC-RBS-Luc | GCTTAGATCAGGTGATTGCCCTTTGTTTATGAGGGTGTTGTAATCC<br>ATGTCGTTGTTGCATTTGTAAGGGCAACACCTCAGCCTGCAGGCA<br>GGCACTGAAGATACCAAAGGGTAGTTCAGATTACACGGTCACCTG<br>GAAAGGGGGCCATTTTACTTTTTATCGCCGCTGGCGGTGCAAAGT<br>TCACAAAGTTGTCTTACGAAGGTTGTAAGGTAAAACTTATCGATTT<br>GATAATGGAAACGCATTAGCCGAATCGGCAAAAATTGGTTACCTTA<br>CATCTCATCGAAAACACGGAGGAAGTATAGATGTCTAGAGAAAGA<br>GGAGAAATACTAGatggaagacgccaaaaacataaagaaaggcccggcgccattcta<br>tccgctggaagatggaaccgctggagagcaactgcataaggctatgaagagatacgccctggtt<br>cctggaacaattgcttttacagatgcacatatcgaggtggacatcacttacgctgagtacttcgaaat<br>gtccgttcggttggcagaagctatgaaacgatatgggctgaatacaaatcacagaatcgtcgtatg<br>cagtgaaaactctcttcaattctttatgccggtgttgggcgcgttatttatcggagttgcagttgcgccc<br>gcgaacgacatttataatgaacgtgaattgctcaacagtatgggcatttcgcagcctaccgtggtgtt<br>cgtttccaaaaaggggttgcaaaaaattttgaacgtgcaaaaaaagctcccaatcatccaaaaa<br>attattatcatggattctaaaacggattaccagggatttcagtcgatgtacacgttcgtcacatctcatc<br>tacctcccggttttaatgaatacgattttgtgccagagtccttcgatagggacaagacaattgcactg<br>atcatgaactcctctggatctactggtctgcctaaaggtgtcgctctgcctcatagaactgcctgcgtg<br>agattctcgcatgccagagatcctatttttggcaatcaaatcattccggatactgcgattttaagtgttgt<br>tccattccatcacggttttggaatgtttactacactcggatatttgatatgtggatttcgagtcgtcttaat<br>gtatagatttgaagaagagctgtttctgaggagccttcaggattacaagattcaaagtgcgctgctg<br>gtgccaaccctattctccttcttcgccaaaagcactctgattgacaaatacgatttatctaatttacacg<br>aaattgcttctggtggcgctcccctctctaaggaagtcggggaagcggttgccaagaggttccatct<br>gccaggtatcaggcaaggatatgggctcactgagactacatcagctattctgattacacccgagg<br>gggatgataaaccgggcgcggtcggtaaagttgttccattttttgaagcgaaggttgtggatctggat<br>accgggaaaacgctgggcgttaatcaaagaggcgaactgtgtgtgagaggtcctatgattatgtc<br>cggttatgtaaacaatccggaagcgaccaacgccttgattgacaaggatggatggctacattctgg<br>agacatagcttactgggacgaagacgaacacttcttcatcgttgaccgcctgaagtctctgattaag<br>tacaaaggctatcaggtggctcccgctgaattggaatccatcttgctccaacaccccaacatcttcg<br>acgcaggtgtcgcaggtcttcccgacgatgacgccggtgaacttcccgccgccgttgttgttttgga<br>gcacggaaagacgatgacggaaaaagagatcgtggattacgtcgccagtcaagtaacaaccg<br>cgaaaaagttgcgcggaggagttgtgtttgtggacgaagtaccgaaaggtcttaccggaaaactc |

| | |
|---|---|
| | gacgcaagaaaaatcagagagatcctcataaaggccaagaagggcggaaagatcgccgtgt<br>aaactttatctgagaatagtcaatcttcggaaatcccaggtggcatgctaaaagtctcgtaaagcgt<br>tctatcaataacccgttggtgccaggcatcaaataaaacgaaaggctcagtcgaaagactgggc<br>ctttcgttttatctgttgtttgtcggtgaacgctctctactagagtcacactggctcaccttcgggtgggcc<br>tttctgcgtttataccgtctcagaatcggccgtgaacaataaaatagtttcggtattattgaccacttcc<br>gagtagaatcgtgcttcagtaagagtcgaccgatgcccttgagagccttcaacccagtcagctcctt<br>ccggtgggcgcggggcaTGACTATCGTCGCCGCACTTATGACTGTCTTCT<br>TTATCATGCAACTCGTAGGACAGGTGCCGGCAGCGCTCTTCCGCT<br>TCCTCGCTCACTGACTCGCTGCGCTCGGTCGTTCGGCTGCGGCG<br>AGCGGTATCAGCTCACTCAAAGGCGGTAATACGGTTATCCACAGA<br>ATCAGGGGATAACGCAGGAAAGAACATGTGAGCAAAAGGCCAGC<br>AAAAGGCCAGGAACCGTAAAAAGGCCGCGTTGCTGGCGTTTTTCC<br>ATAGGCTCCGCCCCCCTGACGAGCATCACAAAAATCGACGCTCAA<br>GTCAGAGGTGGCGAAACCCGACAGGACTATAAAGATACCAGGCG<br>TTTCCCCCTGGAAGCTCCCTCGTGCGCTCTCCTGTTCCGACCCTG<br>CCGCTTACCGGATACCTGTCCGCCTTTCTCCCTTCGGGAAGCGTG<br>GCGCTTTCTCAATGCTCACGCTGTAGGTATCTCAGTTCGGTGTAG<br>GTCGTTCGCTCCAAGCTGGGCTGTGTGCACGAACCCCCCGTTCA<br>GCCCGACCGCTGCGCCTTATCCGGTAACTATCGTCTTGAGTCCAA<br>CCCGGTAAGACACGACTTATCGCCACTGGCAGCAGCCACTGGTAA<br>CAGGATTAGCAGAGCGAGGTATGTAGGCGGTGCTACAGAGTTCTT<br>GAAGTGGTGGCCTAACTACGGCTACACTAGAAGGACAGTATTTGG<br>TATCTGCGCTCTGCTGAAGCCAGTTACCTTCGGAAAAAGAGTTGG<br>TAGCTCTTGATCCGGCAAACAAACCACCGCTGGTAGCGGTGGTTT<br>TTTTGTTTGCAAGCAGCAGATTACGCGCAGAAAAAAAGGATCTCAA<br>GAAGATCCTTTGATCTTTTCTACGGGGTCTGACGCTCAGTGGAAC<br>GAAAACTCACGTTAAGGGATTTTGGTCATGAGATTATCAAAAAGGA<br>TCTTCACCTAGATCCTTTTAAATTAAAAATGAAGTTTTAAATCAATCT<br>AAAGTATATATGAGTAAACTTGGTCTGACAGTTACCAATGCTTAAT<br>CAGTGAGGCACCTATCTCAGCGATCTGTCTATTTCGTTCATCCATA<br>GTTGCCTGACTCCCCGTCGTGTAGATAACTACGATACGGGAGGGC<br>TTACCATCTGGCCCCAGTGCTGCAATGATACCGCGAGACCCACGC<br>TCACCGGCTCCAGATTTATCAGCAATAAACCAGCCAGCCGGAAGG<br>GCCGAGCGCAGAAGTGGTCCTGCAACTTTATCCGCCTCCATCCAG<br>TCTATTAATTGTTGCCGGGAAGCTAGAGTAAGTAGTTCGCCAGTTA<br>ATAGTTTGCGCAACGTTGTTGCCATTGCTACAGGCATCGTGGTGT<br>CACGCTCGTCGTTTGGTATGGCTTCATTCAGCTCCGGTTCCCAAC<br>GATCAAGGCGAGTTACATGATCCCCCATGTTGTGCAAAAAAGCGG<br>TTAGCTCCTTCGGTCCTCCGATCGTTGTCAGAAGTAAGTTGGCCG<br>CAGTGTTATCACTCATGGTTATGGCAGCACTGCATAATTCTCTTAC<br>TGTCATGCCATCCGTAAGATGCTTTTCTGTGACTGGTGAGTACTCA<br>ACCAAGTCATTCTGAGAATAGTGTATGCGGCGACCGAGTTGCTCT<br>TGCCCGGCGTCAATACGGGATAATACCGCGCCACATAGCAGAACT<br>TTAAAAGTGCTCATCATTGGAAAACGTTCTTCGGGGCGAAAACTCT<br>CAAGGATCTTACCGCTGTTGAGATCCAGTTCGATGTAACCCACTC<br>GTGCACCCAACTGATCTTCAGCATCTTTTACTTTCACCAGCGTTTC<br>TGGGTGAGCAAAAACAGGAAGGCAAAATGCCGCAAAAAAGGGAAT<br>AAGGGCGACACGGAAATGTTGAATACTCATACTCTTCCTTTTTCAA<br>TATTATTGAAGCATTTATCAGGGTTATTGTCTCATGAGCGGATACA<br>TATTTGAATGTATTTAGAAAAATAAACAAATAGGGGTTCCGCGCAC<br>ATTTCCCCGAAAAGTGCCACCTGACGTCTAAGAAACCATTATTATC<br>ATGACATTAACCTATAAAAATAGGCGTATCACGAGGCCCTTTCGTC<br>TTCAAGAATTCTGGCGAATCCTCTGACCAGCCAGAAAACGACCTTT<br>CTGTGGTGAAACCGGATGCTGCAATTCAGAGCGGCAGCAAGTGG |

| | |
|---|---|
| | GGGACAGCAGAAGACCTGACCGCCGCAGAGTGGATGTTTGACAT GGTGAAGACTATCGCACCATCAGCCAGAAAACCGAATTTTGCTGG GTGGGCTAACGATATCcgcctgatgcgtgaacgtgacggacgtaaccaccgcgacat gtgtgtgctgttccgctggctcggatacccttactctgttgaaaacgaatagataggttaaggaacg gttatttctgcgtagatctatcttacacagcatcacactggctcaccttcgggtgggcctttctgcgtttat atactagagagagaatataaaaagccagattattaatccggctttttattatttaggcaactgaaac gattcggatcctgtattactattctta |
